# Nucleoporin-binding nanobodies that either track or inhibit nuclear pore complex assembly

**DOI:** 10.1101/2023.09.12.557426

**Authors:** Mireia Solà Colom, Zhenglin Fu, Thomas Güttler, Sergei Trakhanov, Vasundara Srinivasan, Kathrin Gregor, Tino Pleiner, Dirk Görlich

## Abstract

Nuclear pore complex (NPC) biogenesis is a still enigmatic example of protein self-assembly. We now introduce several cross-reacting anti-Nup nanobodies for imaging intact NPCs from frog to human. We further report a simplified assay that directly tracks postmitotic NPC assembly by added labeled anti-Nup nanobodies. In interphase, NPCs are inserted into a pre-existing nuclear envelope. This makes it difficult to monitor this process as newly-assembled NPCs must be distinguished from pre-existing ones. We solved this problem by inserting *Xenopus* NPCs into human nuclear envelopes and using frog-specific anti-Nup nanobodies for detection. We also asked whether anti-nucleoporin (Nup) nanobodies could serve as NPC assembly inhibitors. A first generation, selected from immune libraries against *Xenopus* Nups, comprised only bright stainers of intact NPCs but no inhibitors, perhaps because the immune response was biased towards non-conserved and, thus, functionally-irrelevant epitopes. To overcome this bias, we selected for crossreactivity between *Xenopus* and human Nups and obtained anti-Nup93, Nup98, and Nup155 nanobodies that block Nup-Nup interfaces and arrest NPC assembly. We solved structures of nanobody-target complexes and identified roles for the Nup93-α-solenoid in recruiting Nup358 and the Nup214·88·62 complex, and for Nup155 and the Nup98 autoproteolytic domain in NPC-scaffold assembly. The latter suggests an assembly checkpoint linking pore formation to permeability barrier assembly.

## Introduction

The nucleus and cytoplasm of eukaryotic cells are separated by the nuclear envelope (NE), which comprises two concentric lipid bilayers: an outer nuclear membrane (ONM) and an inner nuclear membrane (INM). This separation protects the genome, allows proper handling of introns, and provides sophisticated control over gene expression. However, it also necessitates a dedicated infrastructure for communication and fine-tuned exchange between the two compartments (Cavalier-Smith, 1988). Nucleocytoplasmic transport proceeds through NE-embedded nuclear pore complexes (NPCs; reviewed in Görlich and Kutay, 1999; Wing, C.E., Fung, H.Y.J. & Chook, 2022). With a molecular mass of around 125 MDa and an outer diameter of ∼ 120 nm, NPCs are among the cells’ largest protein complexes. They are composed of about 30 different proteins called nucleoporins, or Nups for short (Fig1A; reviewed in Knockenhauer and Schwartz, 2016). The NPC scaffold has octagonal symmetry and is organized in three coaxial, stacked rings. The inner ring (IR) encloses the central transport channel, and it is flanked by two outer rings: the cytoplasmic ring (CR) and the nuclear ring (NR) (Fig 1A). Several Nups pre-assemble into sub-complexes. The Y-complex, named for its Y-shaped structure, is the largest Nup complex and the most prominent building block of the outer rings (Siniossoglou et al., 2000; Harel et al., 2003b; Walther et al., 2003a; Beck et al., 2004, 2007). In vertebrates, it consists of nine proteins, namely Nup160, Nup133, Nup107, Nup96, Nup75, Nup43, Nup37, Sec13, and SehI.

**Figure 1.**
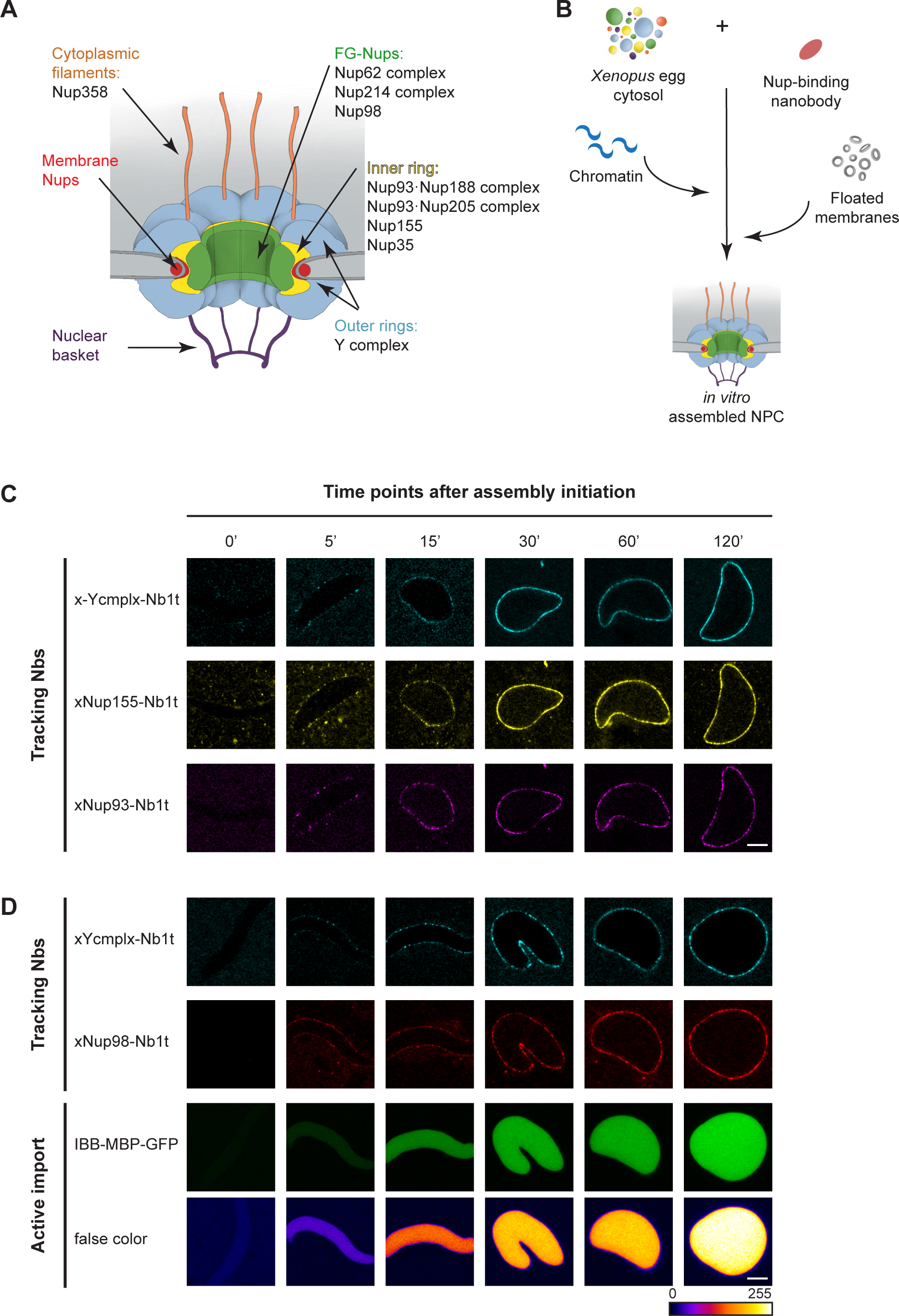
Anti-Nup tracking nanobodies allow to follow NPC assembly from *Xenopus* egg extracts. **(A)** Schematic representation of the NPC with locations of the major Nup complexes being indicated. **(B)** Schematic of the nuclear assembly process from *Xenopus* egg extracts in the presence of anti-Nup nanobodies. The soluble fraction of the egg extract is pre-incubated with an anti-Nup nanobody of choice. In the presence of chromatin and floated membranes, NPCs assemble in a test tube. **(C, D)** Nuclei were assembled from *Xenopus* egg extracts in the presence of 50 nM of the indicated tracking nanobodies (see Figure 2A), which is approximately stoichiometric to the endogenous Nups (Wühr et al., 2014) and/or 1 µM IBB-MBP-GFP. Next, sperm chromatin and the *Xenopus* membrane fraction were added, and the assembly reactions were allowed to proceed at 18°C. Confocal images of live nuclei were acquired at the indicated time points. **(C)** The Y-complex, Nup155, and Nup93 are recruited to the NE as nuclear assembly proceeds and nuclei grow in size. **(D)** Reconstituted nuclei become transport-competent as the Y- complex and Nup98 are recruited to assembly sites. IBB-MBP-GFP: a fusion of Importin β- binding domain of importin α to a maltose-binding protein-GFP module. Scale bar, 5 µm.

The Nup93·Nup188 complex (Miller et al., 2000) is a constituent of the inner ring, while recent NPC structures have shown that the (paralogous) Nup93·Nup205 complex is present in all three rings (Bley et al., 2022; Mosalaganti et al., 2022; Zhu et al., 2022). Nup35 (also called Nup53 or MP44) and Nup155 are further structural elements of the inner ring, connecting it through Aladin and the membrane-integral nucleoporin NDC1 to the pore membrane (Eisenhardt et al., 2014; Mosalaganti et al., 2022).

So-called FG Nups equip NPCs with a permeability barrier. Their intrinsically disordered FG repeat domains can engage in cohesive interactions and condense into a selective FG phase, which in turn is highly permeable to shuttling nuclear transport receptors (NTRs) and NTR·cargo complexes, while rejecting inert macromolecules that are not recognized as valid cargoes (Schmidt and Görlich, 2015; Labokha et al., 2013; Frey and Görlich, 2007).

Vertebrate NPCs contain 11 FG Nups, each anchored in a specific way. Nup98 provides the most cohesive and barrier-critical FG domain (Hülsmann et al., 2012). It is anchored by its autoproteolytic domain (APD) either to the flexible N-terminus of Nup96 in the Y-complex (Hodel et al., 2002) or to the β-propeller of Nup88 (Griffis et al., 2003). The N-terminus of Nup93 provides binding sites for two other FG complexes, namely the Nup62·58·54 coiled-coil trimer complex (Chug et al., 2015; Stuwe et al., 2015) and the Nup214·Nup88·Nup62 complex (Griffis et al., 2003; Bley et al., 2022). The Nup62·Nup214·Nup88 complex also appears to contact Nup85 (Fontana et al., 2022).

Y-complexes of the cytoplasmic ring also anchor Nup358/ RanBP2 (Beck et al., 2004, 2007), an FG Nup that forms the cytoplasmic filaments (Walther et al., 2002; Wu et al., 1995; Yokohama et al., 1995). In addition to providing several FG domain segments, Nup358 pentamerizes and functions as an architectural element of the cytoplasmic ring that stabilizes interactions between Y-complexes (Von Appen et al., 2015; Zhu et al., 2022; Bley et al., 2022; Mosalaganti et al., 2022).

In animals, NPCs form *de novo* from individual Nup complexes at two distinct stages of the cell cycle: upon exit from mitosis and during interphase. In mitosis, NPCs and NE disassemble prior to chromosome segregation, resulting in soluble Nups and Nup complexes as well as in the dispersal of membrane-integral Nups into vesicles/NE remnants (reviewed by Kutay et al., 2021). After cell division, NPCs re-assemble from these building blocks in a synchronous wave. Fusion events between membrane sheets and vesicles not only re-form a closed NE, but are also thought to directly establish the special topology of the pore membrane.

Extracts from activated *Xenopus* eggs have been widely used to reconstitute post-mitotic NPC assembly in the test tube (Newport, 1987; Finlay and Forbes, 1990; Marshall and Wilson, 1997; Bernis and Forbes, 2015). *Xenopus* eggs are laid in metaphase of meiosis II. Their nuclei are disassembled and contain large stockpiles of all NPC and NE components. Upon activation, they return to an interphase state and become assembly-competent. In a typical nuclear assembly experiment, sperm chromatin is added as an assembly template. Within an hour, nuclei form that contain functional NPCs, exclude inert macromolecules, and actively accumulate supplied import cargoes. This system thus recapitulates NPC and NE formation at the end of cell division and allows the analysis of phenotypes that would be lethal in a cellular setting ( 1B).

A powerful approach has been to deplete specific Nup components from *Xenopus* egg extracts and analyze the consequences for the assembly process (reviewed in Antonin et al., 2008; Schellhaus et al., 2016). This approach revealed essential functions of the Y-complex (Walther et al., 2003a; Harel et al., 2003b; Franz et al., 2007; Rasala et al., 2008), Nup35 (Vollmer et al., 2012), Nup93 (Sachdev et al., 2012), and Nup155 (Franz et al., 2005) in NPC scaffold assembly. This way, it has also been shown that a functional permeability barrier requires cohesive Nup98 FG repeat interactions (Hülsmann et al., 2012). However, Nup depletions are tedious; they may remain incomplete, may damage the extracts in a non-specific manner, or may not be feasible for all components. Furthermore, removing an entire protein or Nup complex abolishes multiple protein-protein interactions and introduces a very dramatic change in the system. Therefore, it is never straightforward to define which assembly step(s) are specifically affected or whether the observed phenotypes correspond to actual intermediates.

In contrast to the post-mitotic assembly mode, NPCs are inserted into an intact NE during interphase. This requires a pore-forming fusion event between the inner and outer nuclear membranes (reviewed in Otsuka and Ellenberg, 2018) and is also more fundamental as only this NPC biogenesis pathway is available in organisms with closed mitosis. Interphase NPC assembly has been followed by examining the increase in NPC numbers after the completion of postmitotic NPC assembly from *Xenopus* egg extracts (D’Angelo et al., 2006; Vollmer et al., 2015) and also visualized by live cell and electron microscopy (Goldberg et al., 1997; Dultz et al., 2008; Otsuka et al., 2016). The earliest assembly event has been proposed to be the direct binding of Nup153 to the INM. Nup153, in turn, is thought to recruit the Y complex to new assembly sites (Vollmer et al., 2015). Subsequently, additional Nups appear to be recruited, perhaps contributing mechanical force to deform the NE. The local fusion between the INM and ONM seems to require Torsins (Laudermilch et al., 2016; Rampello et al., 2020; Prophet et al., 2022), but is otherwise only poorly understood.

It is generally accepted that interphase and postmitotic NPC assembly proceed through a stepwise sequence of structurally defined intermediates, and fundamental mechanistic differences between the two pathways have been described (Antonin et al., 2008; Dultz and Ellenberg, 2010; Otsuka et al., 2016, 2018). Nevertheless, either pathway is still poorly understood, mainly because it has been very difficult to identify assembly intermediates, arrange them in temporal order, and characterize them both biochemically and structurally.

In the present study, we introduce the use of Nup-specific nanobodies to investigate the mechanisms of both NPC assembly modes. Nanobodies (Nbs) are the isolated variable domains of heavy chain-only IgGs (Casterman et al., 1993). They are small in size (∼12-15 kDa) and can be selected for very high affinity (with dissociation constants down to the picomolar range) and high solubility. In addition, functional nanobodies can be robustly produced in microorganisms at high yields and easily labeled with fluorescent dyes (reviewed in Cheloha et al., 2020; Helma et al., 2015; Ingram et al., 2018; Muyldermans, 2013). We (Pleiner et al., 2015; Chug et al., 2015) and others (Nordeen et al., 2020) have previously employed nanobodies to aid in biochemical and structural characterization of NPCs. Here, we extend the anti-Nup nanobody toolbox to investigate the mechanisms of NPC biogenesis and overcome the limitations of the currently available assembly assays.

Our first set of nanobodies, termed tracking Nbs (t), bind soluble Nup complexes and intact NPCs. These nanobodies allow for robust Nup tracking along the course of postmitotic NPC assembly by simply adding them in a labeled form to an assembly reaction and detecting them by direct fluorescence microscopy. We also present a novel approach to study the insertion of NPCs into an intact NE. It relies on human cell nuclei as NPC insertion-templates, a GFP-Nup107 fusion to mark pre-existing NPCs, interphase *Xenopus* egg extract to supply nucleoporins that indeed assemble new NPCs into human NEs, and tracking nanobodies that selectively stain and thus identify newly inserted *Xenopus* NPCs. The system is flexible in terms of which Nups are being tracked, and it allows for biochemical manipulation.

A second set, termed inhibitory Nbs (i), block essential Nup-Nup interactions and arrest the assembly of functional NPCs by targeting conserved Nup epitopes that get buried during NPCs assembly. All inhibitory nanobodies (i) recognize conserved Nup epitopes and thus cross-react between *Xenopus* (x) and human (h) Nups, whereas tracking nanobodies (t) can be either cross-reactive (xh) or *Xenopus*-specific (x). We describe an anti-Nup93 inhibitory nanobody that prevents the incorporation of Nup358 and the Nup214·88·62 complex, indicating a specific arrest in the assembly of the cytoplasmic ring. In addition, inhibitory nanobodies targeting distinct domains of Nup155 and the autoproteolytic domain (APD) of Nup98 block very early steps. The requirement of the Nup98 APD for the assembly of the NPC scaffold suggests that the formation of NE-perforating pores is closely linked to the establishment of the permeability barrier – perhaps to avoid assembly intermediates with non-selectively open pores.

## Results and Discussion

### Anti-Nup nanobodies for tracking post-mitotic NPC assembly from *Xenopus* egg extracts

We have previously generated a toolbox of *Xenopus*-specific anti-Nup nanobodies that recognize fully-assembled NPCs (Pleiner et al., 2015). These nanobodies were directly coupled to fluorophores, eliminating the need for a secondary reagent, and they yielded bright and specific fluorescent signals at the NE of *Xenopus* cells. We now asked whether these nanobodies could simplify the standard procedures for assaying nuclear pore formation from *Xenopus* egg extracts.

We initiated nuclear assembly reactions in the presence of an IBB-MBP-GFP fusion as a fluorescent import cargo (where MBP is the *E. coli* maltose-binding protein and IBB is a strong nuclear import signal, namely the importin β-binding domain of importin α; Görlich et al., 1996). We also added combinations of various anti-Nup nanobodies carrying compatible fluorophores. This allowed us to acquire confocal images of live nuclei at distinct assembly stages, avoiding additional sample preparation steps. The anti-Nup nanobodies did not interfere with the formation of functional NPCs, allowing the recruitment of different Nups at the reforming NE to be tracked (Fig 1C). Furthermore, the included fluorescent transport cargo allows to correlate Nup recruitment with the formation of functional NPCs (Fig 1D), as the accumulation of an import substrate requires a properly installed permeability barrier (reviewed in Knockenhauer and Schwartz, 2016). Therefore, the nanobodies shown in Fig 1C-D (x-Ycmplx-Nb1t, xNup155-Nb1t, xNup93-Nb1t, and xNup98-Nb1t) as well as the additional ones introduced here (*i.e.*, xhNup133- Nb2t, xhNup93-Nb3t, xhNup35-Nb1t, xNup62-Nb1t, xh-Nup214-Nb1t, xhNup358-Nb1t, and xhNup358-Nb2t) (Fig 2A-B, see below) allow for a straightforward analysis of the NPC assembly process, avoiding tedious staining and fixation protocols. From now on, we will refer to them as “tracking nanobodies” (indicated by a “t” following the nanobody number).

**Figure 2.**
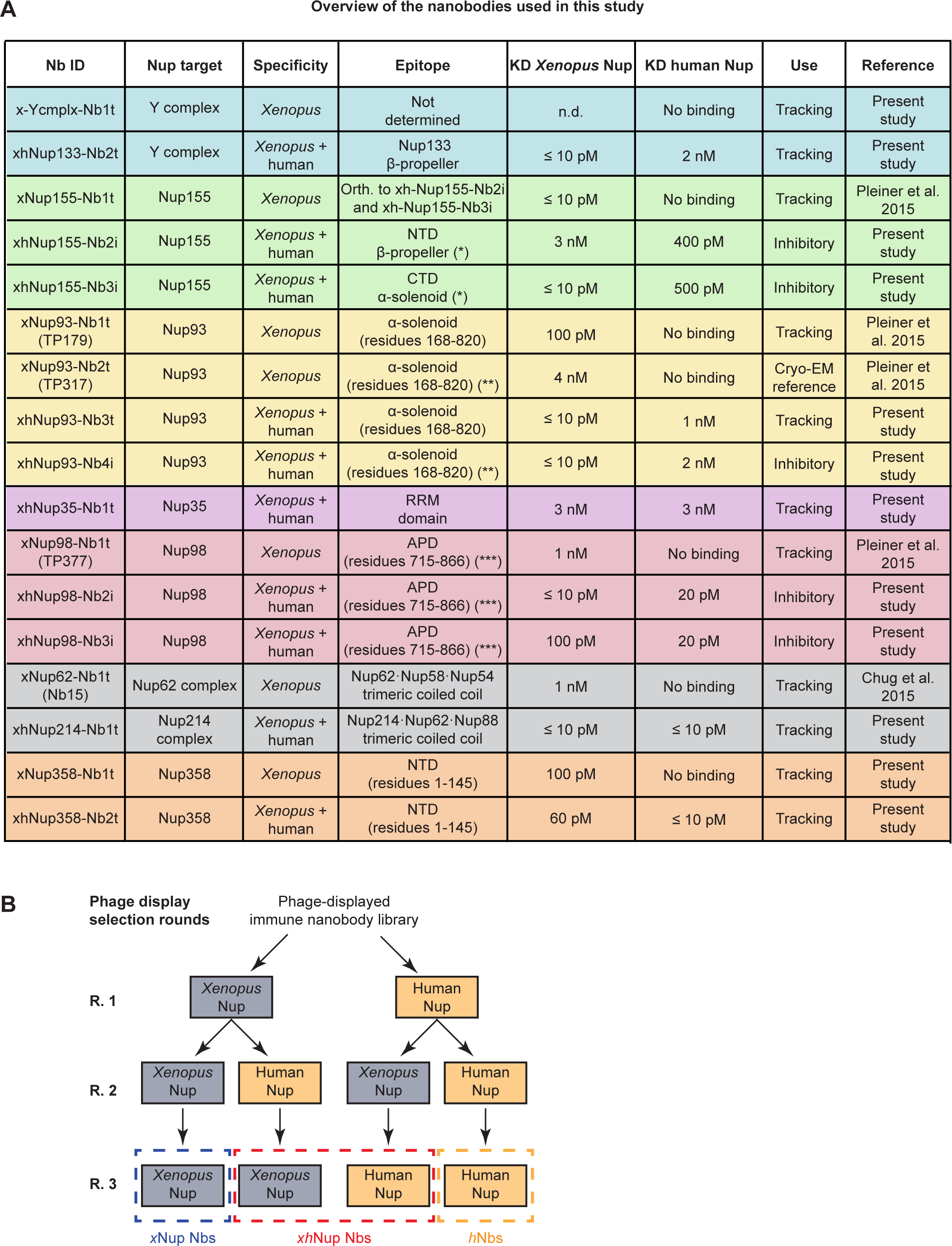
Overview of anti-Nup nanobodies and phage-display strategy used in this study. **(A)** List of anti-Nup nanobodies. The targeted Nup, specificity, affinities estimated by biolayer interferometry, epitope, and use of all nanobodies are indicated. NTD: N-terminal domain. CTD: C-terminal domain. APD: autoproteolytic domain of Nup98. orth.: orthogonal. n.d.: not determined. (*) See Fig S6; (**) see Fig 10; (***) see Figs. 11 and S1. **(B)** Scheme of the used phage display strategy to identify nanobodies with tailored specificities. *x*Nup Nbs: *Xenopus*- specific anti-Nup nanobodies, *xh*Nup: *Xenopus*-human cross-reactive anti-Nup nanobodies, *h*Nup Nbs: human-specific anti-Nup nanobodies.

### An *in vitro* system to track NPC formation during interphase

Compared to postmitotic NPC assembly, interphase NPC formation is more challenging to track because there are comparatively few sporadic assembly events spread over a relatively long period of time. Moreover, the newly-inserted NPCs are indistinguishable from the pre-existing (postmitotic) ones; they look identical (reviewed in Otsuka and Ellenberg, 2018; Weberruss and Antonin, 2016). Nevertheless, we wanted to establish a readily accessible experimental assay for interphase NPC assembly that allows direct biochemical manipulation. This requires not only that the process occurs in a test tube, but also that newly inserted pores can be distinguished from pre-existing ones. To this end, we considered assembling NPCs from a *Xenopus* egg extract into human NEs. Newly-inserted NPCs would then be of *Xenopus* origin and thus chemically distinct from the pre-existing human NPCs. *Xenopus*-specific tracking nanobodies should then label only newly assembled NPCs (see Fig 3A for a scheme). However, given the great evolutionary distance between frogs and humans, it was initially unclear whether the NPC components and assembly machineries would still be compatible enough for such an interspecies experiment to work.

**Figure 3.**
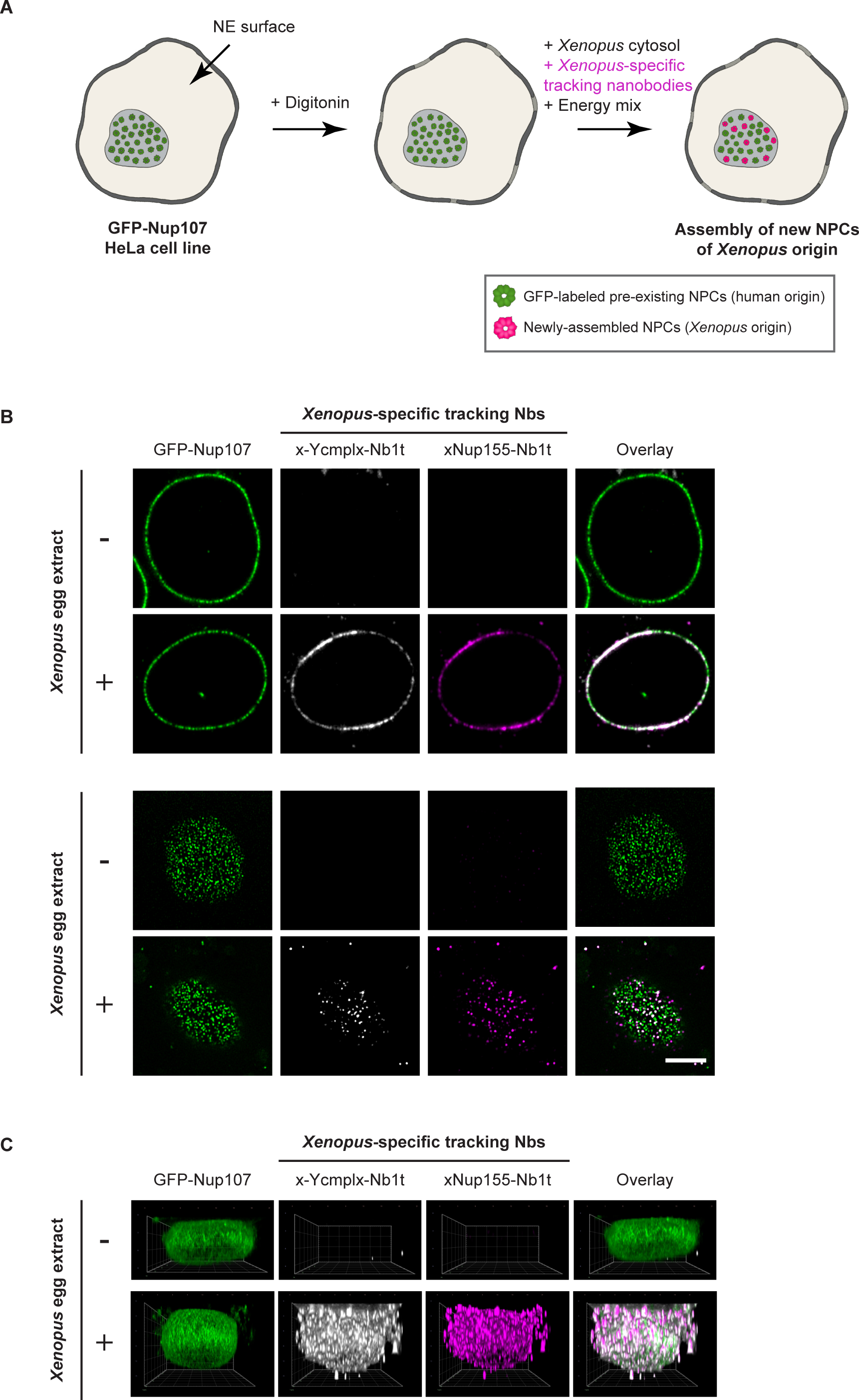
New NPCs composed of *Xenopus* Nups are inserted into human NEs, recapitulating the interphase mode of NPC assembly. **(A)** Scheme of the assay for tracking NPC assembly during interphase. *Xenopus* egg extract (soluble, high-speed fraction) as a Nup source, fluorophore-labeled tracking nanobodies, and energy mix are added to digitonin-permeabilized, GFP-tagged Nup107 HeLa cells to initiate NPC assembly into the intact human NE. In this setup, the newly inserted NPCs are of *Xenopus* origin and can be specifically stained by the *Xenopus*- specific nanobodies. **(B)** Either buffer (-) or *Xenopus* egg extract (+) was mixed with the indicated *Xenopus*-specific tracking nanobodies conjugated to Alexa 568 (x-Ycmplx-Nb1t) or Alexa 647 (xNup155-Nb1t) fluorophores at two or three engineered cysteines (Pleiner et al., 2015) and incubated with digitonin-permeabilized HeLa cells (with GFP-tagged Nup107) for 2 hours at room temperature (RT). Confocal images were acquired using a Zeiss LSM880 confocal microscope. The *Xenopus*-specific tracking nanobodies yielded a bright, specific signal at the midplane (upper panels) and a dot-pattern signal at the NE surface of HeLa nuclei (lower panels), which is consistent with the insertion of new NPCs. Scale bar, 5 µm. **(C)** 3D reconstructions of HeLa nuclei were obtained from the acquired 3-channel Z-stacks along the bottom NE using the Arivis Vision4D software (version 3.1.3; VisionVR, 2020).

To mark pre-existing human NPCs, we used a CRISPR/Cas9-edited HeLa cell line whose Nup107 carries an N-terminal sfGFP tag (Gunkel et al., 2021). We grew these cells on coverslips, permeabilized their plasma membranes with digitonin, and incubated them with a high-speed extract of activated *Xenopus* eggs, which is a rich source of assembly-ready Nups and Nup complexes. To track the assembly of new nuclear pores, we added fluorophore-labeled nanobodies that recognize, for example, the *Xenopus* Y-complex or Nup155.

In equatorial confocal sections of the incubated nuclei, the *Xenopus*-specific tracking nanobodies produced a bright fluorescent signal that coincided (at low resolution) with the GFP signal, indicating that *Xenopus* Nups had accumulated at the human NE (Fig 3B, top panel). In nuclear surface scans, the tracking nanobodies produced a fluorescent spot pattern similar to the GFP signal, which is also consistent with the formation of new NPCs (Fig 3B, bottom panel). Importantly, the nanobody signal was strictly dependent on the addition of egg extract, confirming the species-specificity of the tracking nanobodies and the *Xenopus* origin of the labeled NPCs.

During extract incubation, the initially flat HeLa nuclei became rounder; they grew in volume, suggesting that a massive import of *Xenopus* nuclear proteins had occurred, inflating nuclei through their osmotic-colloidal pressure (Fig 3C). Thus, when new NPCs were assembled, they also contained a functional permeability barrier that kept the imported material inside the nuclei.

To assess whether different *Xenopus* Nups colocalized with each other on the detected spots, we stained the samples with various combinations of tracking nanobodies and imaged them at higher resolution. Fig 4A shows that newly-assembled NPCs can be clearly identified as hNup107-GFP-free but *Xenopus* Y-complex, Nup155, and Nup93-colocalizing spots. *Xenopus* Nup358 and the Nup62 complex also colocalized with the *Xenopus* Y-complex, indicating that barrier-forming FG Nups and the central organizer of the cytoplasmic NPC ring are also recruited to the newly inserted pores. Strikingly, the *Xenopus*-specific xNup98-Nb1t and a *Xenopus*-specific anti-Nup153 antibody colocalized with *Xenopus* NPCs - as expected from a faithful assembly process - but also gave bright signals co-localizing with GFP-positive old human NPCs (Fig 4B). Since these reagents do not recognize the human proteins, as confirmed by the lack of staining in the absence of egg extract, this suggests either that the pre-existing NPCs still had vacant binding sites for Nup98/ Nup153, or that some exchange between NPC-bound and free populations occurred over time. The latter would be consistent with the reported mobility of these Nups (Griffis et al., 2002a, 2004). It should be noted, however, that the huge excess of these FG Nups in the added extract is likely to increase their off-rates - because their anchorage through multiple weak interactions is easily outcompeted stepwise by the excess molecules.

**Figure 4.**
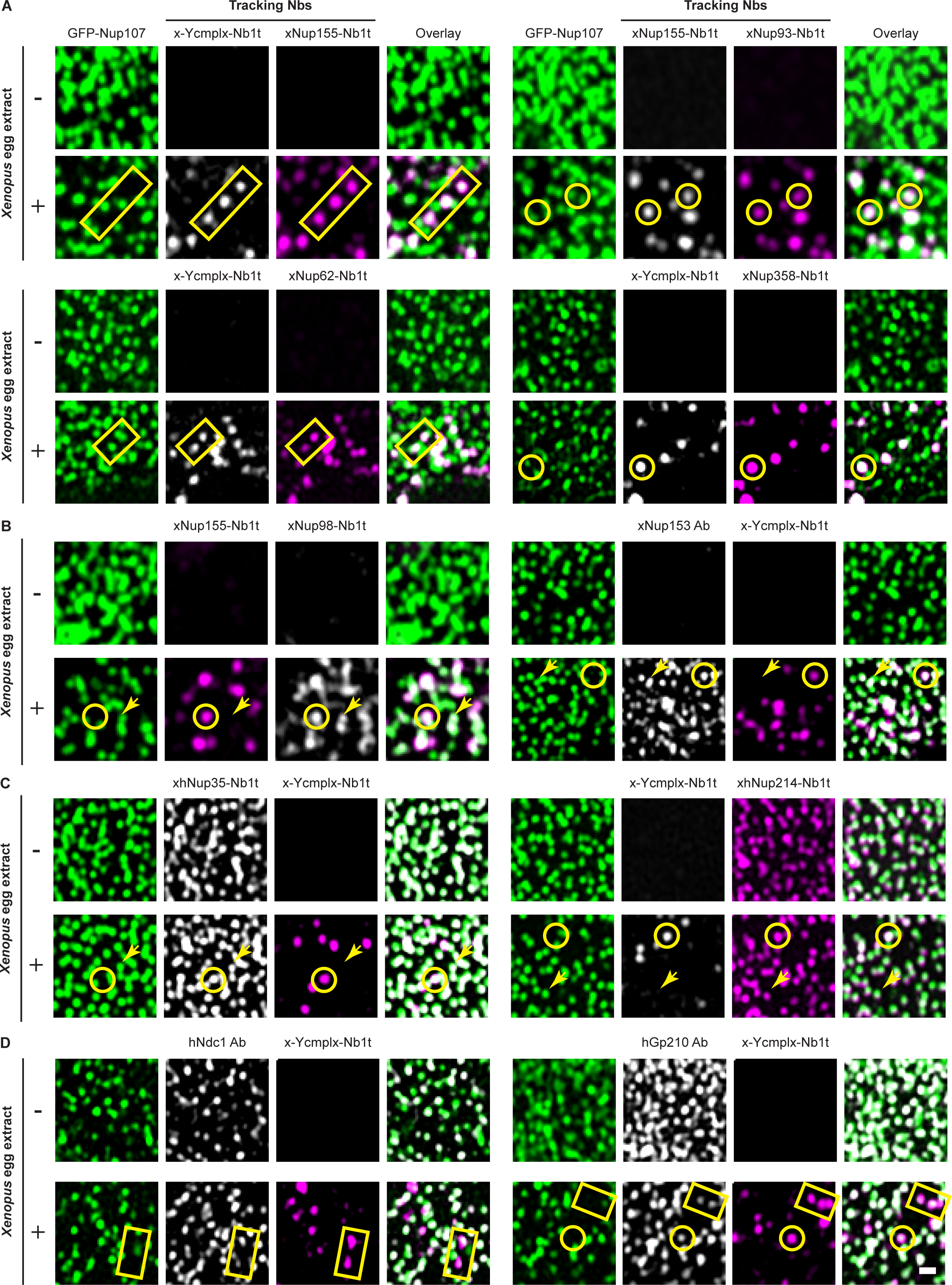
Tracking nanobodies with *Xenopus*-human species specificity distinguish between pre-existing NPCs, newly-assembled NPCs, and Nups that are exchanged between them. New pore insertion was initiated as described in Fig 3, in the presence of the indicated tracking nanobodies and either buffer (-) or *Xenopus* egg extract (+). (A) *Xenopus* Y-complex co-localizes with *Xenopus* Nup155, *Xenopus* Nup62, and *Xenopus* Nup358 in newly assembled pores. *Xenopus* Nup155 also co-localizes with *Xenopus* Nup93 in all cases. (B) *Xenopus* Nup98 and *Xenopus* Nup153 are detected in both pre-existing and newly-assembled NPCs. The minus extract controls indicate that this is not due to cross-reaction but to either complementing vacant binding sites in old pores or to subunit exchange. **(C)** Cross-specific xhNup35-Nb1t and xhNup214-Nb1t co-localize with both pre-existing and newly-assembled NPCs. **(D)** Human membrane Nups support the assembly of *Xenopus* NPCs. Cells were stained with x-Ycmplx-Nb1t coupled to Alexa647 and antibodies against human Ndc1 and human gp210 (Stavru et al., 2006a; b). Circles or rectangles mark newly-inserted pores; arrows point to pre-existing NPCs. Prefixes indicating species specificity: *x*: *Xenopus-*specific; *xh*: cross-reaction between *Xenopus* and human. Scale bar, 0.5 µm.

The presented approach relies on a high species-specificity of the used tracking nanobodies (Fig 2A), which can indeed be explained by the corresponding nanobody-target structures. In the case of xNup98-Nb1t, an arginine residue at CDR3 of the nanobody interacts with an asparagine residue at the APD of *Xenopus* Nup98, but would clash with a lysine at the same position in its human counterpart, preventing its binding (Pleiner et al., 2015; Fig S1A). Similarly, our newly solved cryo-EM structure shows that xNup93-Nb2t fits well into a pocket at the C-terminal of *Xenopus* Nup93, but finds obvious steric clashes with the backbone of human Nup93 at the same protein region (Fig S1B; Table S1).

As expected, cross-specific anti-Nup35 and anti-Nup214 complex nanobodies (see below) stained both pre-existing (GFP-positive) human and newly assembled (GFP-free) *Xenopus* pores as spots of similar size and brightness (Fig 4C). This further documents the architectural equivalence of old and new NPCs, and is thus a key control. Finally, by staining newly-assembled pores with anti-human Ndc1 and -Gp210 antibodies, we confirmed that human membrane Nups present in the HeLa cells support the assembly of the new *Xenopus* NPCs. As shown in Fig 4D, both membrane Nups colocalized with the new pores stained by the *Xenopus-*specific x-Ycmplx-Nb1t.

Interestingly, new NPCs detected in these assay did not cluster in specific membrane regions but maintained a constant and homogenous density on the human NE, which was also observed by Otsuka et al. (2016) in live cell microscopy studies. Furthermore, the fact that no GFP signal was detected in the *Xenopus* Nup structures indicates that the new pores assembled *de novo* and did not originate from pore-splitting events, which is also consistent with previous conclusions (D’Angelo et al., 2006; Dultz and Ellenberg, 2010).

Two hours after the addition of the *Xenopus* extract, *Xenopus* NPCs represented between 25 and 40% of the pre-existing (*i.e.,* human) NPCs and contained similar ratios of all tracked Nups. Longer incubations did not increase the number of assembled *Xenopus* NPCs, suggesting that the system was already saturated after two hours, perhaps because membrane Nups (in particular, the essential membrane Nup Ndc1) had become limiting. Nevertheless, this initial rate of NPC insertion is higher than in cultured HeLa cells (Maul et al., 1972), where the count of NPCs doubles over a period of ∼20 hours (G1+S+G2). We would explain this high rate by the embryonic origin of the added extract and the high concentration of available Nups.

To confirm the insertion of properly assembled NPCs, we acquired higher-resolution images of the HeLa NE containing newly-assembled pores, using two-colour STED (stimulation emission depletion) microscopy (Klar et al., 2000). The anti-GFP nanobodies (labeling Nup107 in the pre-existing human NPCs) allowed to resolve ring-like structures of ∼100-120 nm in diameter, which is consistent with the diameter of the NPC outer rings (Fig 5A). Similarly, the *Xenopus*-specific x- Ycmplx-Nb1t stained rings of a similar shape and diameter, which did not colocalize with the GFP-labelled old NPCs (Fig 5B). STED images of the NE equator confirmed that the *Xenopus* Nups are not only bound but actually embedded in the human NE (Fig S2).

**Figure 5.**
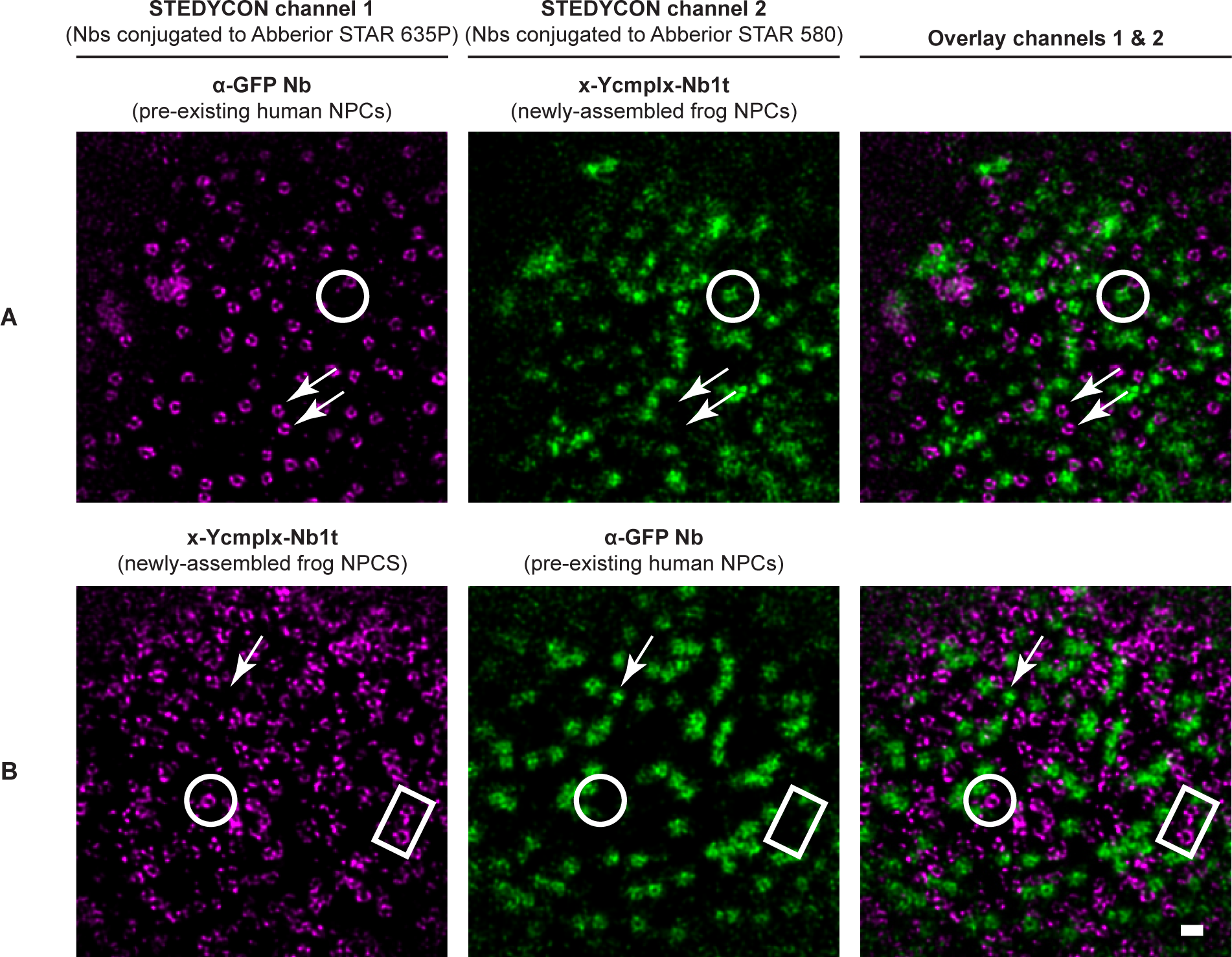
*Xenopus* NPCs newly inserted into human NEs exhibit a ring-like architecture. New pore insertion was initiated as described in Fig 3, in the presence of an anti-GFP nanobody (Kirchhofer et al., 2010) and the indicated tracking nanobodies. Next, images of the NE plane were acquired using a STEDycon microscope. **(A)** The anti-GFP nanobody yielded ring-like structures of ∼ 100-120 nm in diameter, which did not colocalize with the spots labeled by the *Xenopus*- specific x-Ycmplx-Nb1t. **(B)** The *Xenopus*-specific x-Ycmplx-Nb1t yielded ring-like structures of a similar size, which did not colocalize with the GFP-labelled spots. Note that the first channel of the STEDycon system provides a slightly higher resolution than the second; thus, NPC rings can only be resolved on images acquired using the first STED channel. The used nanobodies were labeled at two or three engineered cysteines (Pleiner et al., 2015) with the fluorophore Abberior STAR 635P maleimide for acquiring images on the first STED channel or the Abberior STAR 580 for acquiring images on the second STED channel. Newly-inserted pores are marked by circles or squares; pre-existing pores are indicated by arrows. Scale bar, 200 nm.

The insertion of frog NPCs into a human NE is not a trivial result. Instead, it implies that the ∼30 soluble different frog Nups can still cooperate smoothly with the still elusive human membrane fusion machinery and the at least three human membrane-integral Nups. Indeed, the fact that human Nup98 and human Nup153 can be exchanged by their *Xenopus* counterparts (Fig 4C) shows that even NPCs containing Nups from multiple species are operational. Given that the lineages leading to mammals and frogs diverged ∼350 million years ago (Benton et al., 2015), such inter-species compatibility is remarkable and suggests that functionally-essential Nup-Nup interfaces are well conserved through evolution.

### A cross-species phage display strategy for targeting functional Nup regions

Capturing and characterizing intermediate stages of NPC assembly has remained the main bottleneck in the field. As NPCs are essential structures, such capture can be complicated by lethal phenotypes and is thus particularly challenging in living cells. Biochemical alterations are easier to introduce in *Xenopus* cell-free extracts, where all potential assembly phenotypes are, in principle, accessible. In this system, only a few inhibitors of nuclear pore formation have been described, including wheat germ agglutinin (WGA) (Wiese et al., 1997), BAPTA (1,2-bis(O-aminophenoxy)ethane-N, N, N’, N’-tetraacetic acid) (Macaulay and Forbes, 1996; Wiese et al., 1997; Bernis and Forbes, 2015), and an excess of importin β (Harel et al., 2003a; Walther et al., 2003b). These treatments result in pore-free membranes and thus block NPC assembly at very early stages, providing only limited information on the mechanisms of this process.

We reasoned that nanobodies directed against functional Nup interfaces could prevent relevant Nup⋅Nup interactions and thus arrest the NPC assembly process at discrete intermediate steps. Such nanobodies would act through the smallest possible change in the system without having to deplete any protein. They would provide “epitope resolution” and not just reveal requirements for protein components or entire protein complexes. In addition, nanobodies could be added to the disassembled NPC components in non-limiting amounts, which would ensure that all targeted Nup molecules become trapped in Nup⋅nanobody complexes when NPC assembly is initiated.

To test this idea, we performed nuclear assembly reactions in the presence of various anti-Nup nanobodies selected by Pleiner et al., (2015) (Fig 1C, D); however, none of them affected the formation of functional nuclear pores (see below). While initially disappointing, this outcome became plausible when considering (*i*) that all tested nanobodies had been selected for their bright stain of fully-assembled NPCs (Pleiner et al., 2015), and thus for targeting well-accessible epitopes, and (*ii*) that all of them recognize non-conserved, *Xenopus*-specific epitopes. In contrast, functional protein regions, whose block can arrest NPC assembly, should get masked by protein-protein interactions and be well conserved through evolution. With this in mind, we set out to select nanobodies recognizing conserved Nup epitopes.

Since Pleiner et al., (2015) had immunized alpacas exclusively with *Xenopus* Nups, the antibody response was likely biased towards variable *Xenopus*-specific surfaces that the immune system did not recognize as “self”. To boost an immune response directed against conserved epitopes, we re-immunized the same animals simultaneously with *Xenopus* and human Nup homologs. We chose the same Nup domains that had already been administered by Pleiner et al., (2015), namely the autoproteolytic domain of Nup98, the Nup93 α-solenoid, and the full-length Nup155. In addition, we immunized the N-terminal domain (NTD) of Nup358 (Kassube et al., 2012), the RRM domain of Nup35 (Handa et al., 2006), the β-propeller of Nup133 from the Y-complex (Berke et al., 2004), and the trimeric (Δ FG) Nup62·Nup214·Nup88 complex (Fig 1A, 2A). This focus on well-folded immunogens considered that nanobodies only rarely bind linear epitopes.

We employed three parallel phage display strategies to select binders of a given Nup from the obtained immune libraries: *(i)* a *Xenopus*-specific (multi-round) selection using the *Xenopus* Nup ortholog as a bait, *(ii)* a selection using the human Nup ortholog, and *(iii)* a cross-specific selection alternating between the *Xenopus* and the human Nup orthologs in successive selection rounds, in order to exclusively enrich nanobodies recognizing Nup epitopes that are conserved among the two species (Fig 2B). The bait concentrations were gradually reduced in each selection round to a final concentration below 1 nM to only enrich high-affinity binders. Not surprisingly, we re-discovered the same nanobody classes as Pleiner et al., (2015) in the *Xenopus*-specific selections against Nup98, Nup93, and Nup155. However, all cross-specific pannings selected new nanobody classes – consistent with an enrichment of nanobodies against conserved Nup epitopes. Figure 2A provides an overview of all selected nanobodies used in this study, including estimates for their affinity as measured by biolayer interferometry. The prefix “x” indicates *Xenopus-*specific nanobodies, while “xh” refers to *Xenopus-*human cross-specific ones.

### Cross-specific tracking nanobodies stain human NPCs

We produced representative members of all nanobody classes obtained from the cross-species selections, conjugated them to fluorophores, and tested them for NPC-staining in HeLa cells. A subset, including xhNup133-Nb2t, xhNup93-Nb3t, xhNup35-Nb1t, and xhNup358-Nb2t, yielded bright and specific fluorescent signals at the NE of both fixed triton-permeabilized (Fig 6A) and non-fixed digitonin-semipermeabilized (Fig S3) HeLa cells, suggesting that their epitopes are well exposed in fully-assembled NPCs. Since these nanobodies originate from cross-specific phage display selections, they can also be used to track NPC assembly from *Xenopus* egg extracts, and they would probably stain NPCs from other vertebrate organisms as well.

**Figure 6.**
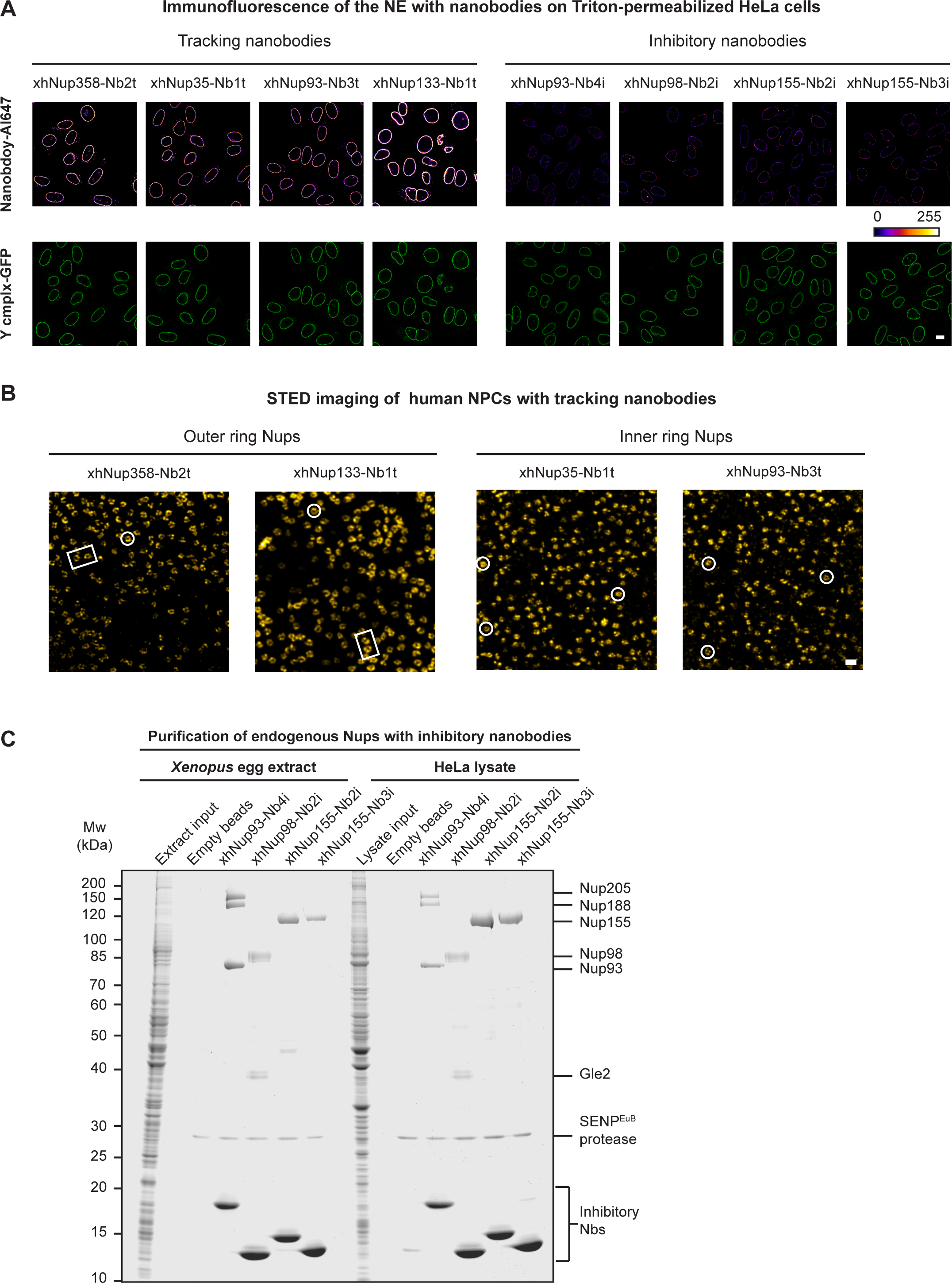
Staining of human NPCs and purification of endogenous Nup complexes with cross-specific (frog-human) nanobodies. **(A)** Four out of the eight tested nanobodies stain NPCs bright at the NE of HeLa cells, indicating that they recognize exposed Nup epitopes. GFP-Nup107 HeLa cells were PFA-fixed, triton-permeabilized, and incubated with 30 nM of the indicated nanobodies coupled to Alexa 647. Confocal cross-sections of the stained nuclei were acquired using a confocal Leica SP8 microscope and identical settings. The GFP signal was used as an internal reference to stain all cells; and a false-color representation (i.e., modified lookup table (LUT)) of the nanobody-stainings is given for a clearer comparison between the different nanobodies. Scale bar, 10 µm. **(B)** Tracking nanobodies allow to resolve the ring-like structure of human NPCs using STED microscopy. HeLa cells were stained with 30 nM of the indicated nanobody coupled to the Abberior STAR 635P fluorophore and imaged using a STEDycon system. Complete NPC rings are highlighted in white. Scale bar, 250 nm. **(C)** Nanobodies that failed to stain HeLa NPCs do purify their human and *Xenopus* Nup targets in a specific manner. The indicated nanobodies were produced with a cleavable N-terminal (biotin)-Avi-SUMO^EU^-tag, immobilized on streptavidin-agarose matrix, and incubated with either HeLa lysate or *Xenopus* egg extract to capture the targeted Nups. Next, the Nup-nanobody complexes were eluted by tag-cleavage with SENP^Eu1^ protease (Vera-Rodriguez et al., 2019) and analyzed by SDS-PAGE/Coomassie-staining.

In addition, these four tracking nanobodies are valuable tools for super-resolution microscopy since they allow to resolve NPC rings on HeLa cell NEs by STED. As expected, nanobodies recognizing Nups of the NPC outer rings (e.g., Nup358, Nup133) labeled rings with larger diameters than nanobodies against inner ring components (e.g., Nup93, Nup35) (Fig 6B).

### Cross-specific nanobodies that recognize buried NPC epitopes

The human-*Xenopus* cross-reacting nanobodies xhNup93-Nb4i, xhNup98-Nb2i, xhNup155-Nb2i, and xhNup155-Nb3i stained NPCs only very weakly, which is particularly evident when compared to the internal Nup107-GFP reference and the in parallel tested tracking nanobodies (Fig 6A, S3). These weak signals could be explained by (*i*) a generally poor affinity to their target Nups or (*ii*) by their respective epitopes being buried by protein-protein interactions in fully-assembled NPCs.

To distinguish between these scenarios, measured affinities for their Xenopus targets and found low nanomolar to low picomolar affinities (Fig 2A). These numbers indicate a tight or even very tight binding. As a complementary experiment, we used the weakly staining nanobodies as baits to purify soluble (unassembled) Nups from *Xenopus* egg extracts and HeLa cell lysates. To do this, we produced them in (biotin)-avi-SUMO^Eu1^-tagged form, immobilized them on streptavidin-agarose, incubated them with extracts, washed extensively, and eluted the nanobodies along with any bound prey by cleaving the SUMO^Eu1^-tag with the engineered SENP^Eu1^ protease (Vera-Rodriguez et al., 2019). This resulted in a clean one-step-purification of Nup93 (with co-purifying Nup188 and Nup205), Nup98 (with co-purifying Gle2), and Nup155 (Fig 6C). In addition, HeLa and *Xenopus* egg extract gave essentially identical results, confirming the cross-species recognition. Thus, the xhNup93-Nb4i, xhNup98-Nb2i, xhNup155-Nb2i, and xhNup155-Nb3i nanobodies recognize their targets specifically and tightly, which in turn is consistent with their low to sub-nanomolar affinities (Fig 2A). The weak staining of intact NPCs is therefore probably due to the burial of their epitopes.

### Some cross-specific nanobodies inhibit the assembly of functional NPCs

Next, we tested all cross-specific nanobodies for their effects on NPC assembly from *Xenopus* egg extracts. All nanobodies that brightly stained NPCs of HeLa cells also permitted the formation of import-competent nuclei and, thus, of functional nuclear pores (Fig 7A), which is not surprising considering that their epitopes are obviously exposed on the NPC surface.

**Figure 7.**
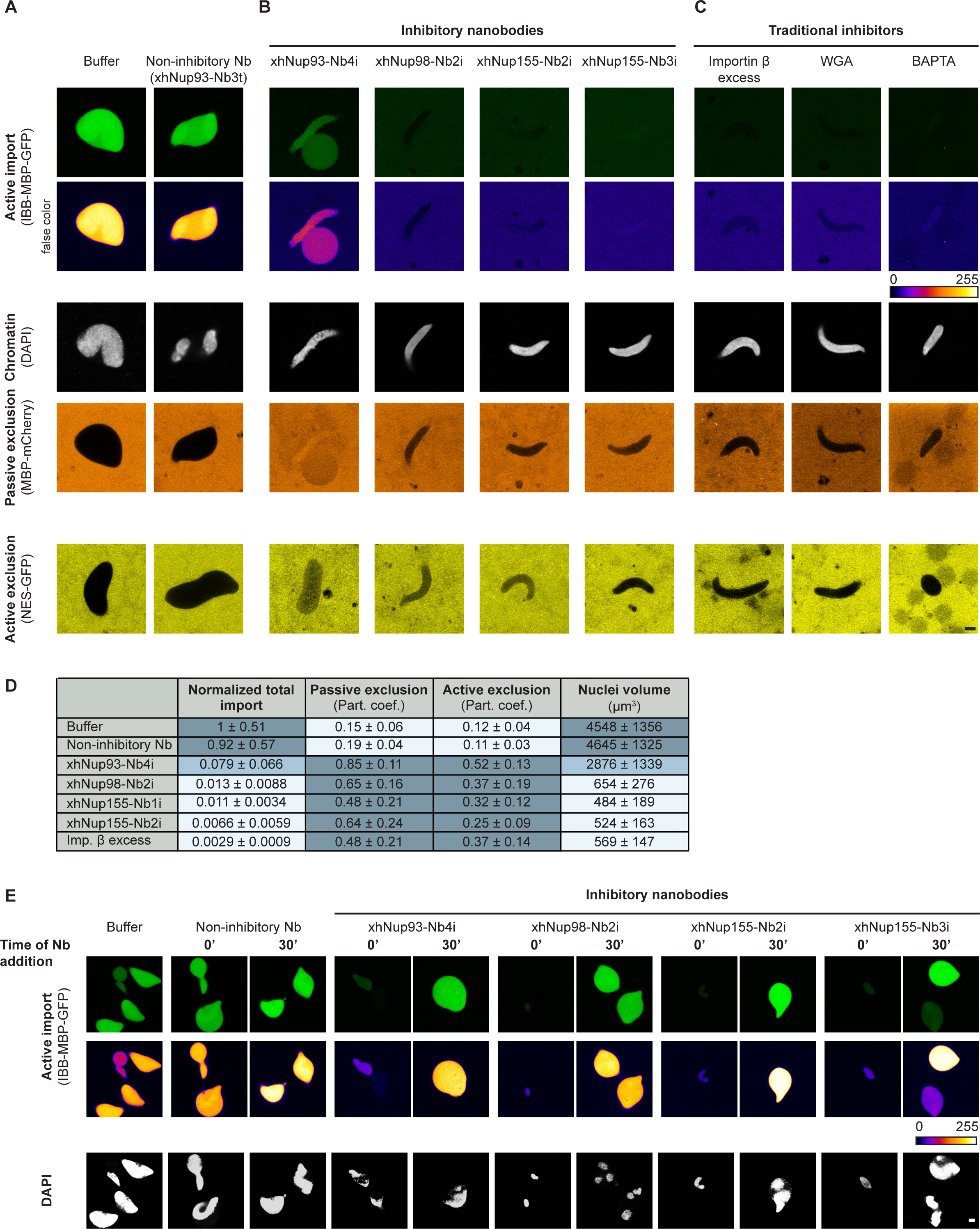
Inhibitory nanobodies arrest NPC assembly from *Xenopus* egg extracts. Nuclear assembly reactions were initiated in the presence of buffer, 2 µM of the indicated anti-Nup nanobody, 2 µM importin β, 2.5 µM WGA, or 5 mM BAPTA, and allowed to proceed for 1 hour at 18°C. Subsequently, DAPI, a fluorescent import cargo (IBB-MBP-GFP), and either a passive nuclear exclusion marker (MBP-mCherry) or an active (Xpo1-dependent) export cargo (NES- GFP) were added. Confocal scans were taken 30 minutes later. Scale bar, 5 µm. **(A)** None of the *Xenopus*-specific or cross-specific nanobodies that brightly stain HeLa cells (see Fig 6) impaired the assembly of functional NPCs, as exemplified by the xhNup93-Nb3t control. **(B)** The four cross-specific Nbs that failed to stain HeLa cells (i.e., inhibitory Nbs; see Fig 6A, S3) and **(C)** the traditional inhibitors of NPC assembly blocked the formation of functional nuclear pores and resulted in nuclei that failed to import the IBB-MBP-GFP fusion protein. **(D)** Total import was quantified as the partition coefficient (Part. coef.) between the mean intensity inside and outside the nuclei multiplied by the nuclear volumes. The obtained values were then normalized to the control values (*i.e.* nuclei assembled in the presence of buffer). Passive and active exclusion were quantified as the partition coefficient (Part. coef.) between the mean intensity outside and inside the nuclei. The nuclei volume was quantified from 3D reconstructions from acquired Z-stacks. In all cases, the mean and SD from 10-50 nuclei from at least four independent experiments are shown. **(E)** Nuclei were assembled as described above, and inhibitory nanobodies were added either prior to chromatin addition and assembly initiation (0’) or 30 minutes after membrane addition. Next, IBB-MBP-GFP and DAPI were added, and nuclei were imaged 90 minutes later. Scale bar, 5 µm.

In contrast, the xhNup93-Nb4i nanobody caused a quite diagnostic phenotype. Import of IBB-MBP-GFP into assembled nuclei was ≥10-fold reduced, and MBP-Cherry and NES-GFP fusions equilibrated between nucleus and cytoplasm – consistent with defects in the (passive) permeability barrier and in Xpo1-mediated nuclear export (Fig 7B, D) (for explanations, see below). Nevertheless, it appears that at least rudimentary NPCs can still assemble in the presence of this nanobody.

A very severe phenotype was evident when xhNup98-Nb2i, xhNup98-Nb3i, xhNup155-Nb2i, or xhNup155-Nb3i were present during the nuclear assembly reaction. Nuclear import of IBB-MBP-GFP was drastically reduced – 100-fold or even to the extent of nuclear exclusion (Figs. 7B, D). In addition, the nuclei remained very small, probably because there was no nuclear import to drive nuclear growth. Indeed, “nuclei” trapped by these nanobodies resemble those assembled in the presence of WGA, an excess of importin β, or BAPTA (Fig 7C), which are known to inhibit nuclear pore formation completely and to result in a pore-free nuclear envelope around the added chromatin (Wiese et al., 1997; Harel et al., 2003a; Bernis and Forbes, 2015).

Strikingly, these nanobodies had no inhibitory effect when added 30 minutes after assembly initiation (Fig 7E). Thus, they do not block nuclear import directly, supporting the notion that they arrest NPC assembly by targeting Nup epitopes that are only transiently accessible. Furthermore, this control is another strong argument against off-target effects (besides their exquisite binding specificities, as documented in Fig 6).

Although all inhibitory nanobodies recognize conserved Nup epitopes, not all cross-species anti-Nup nanobodies induce an assembly phenotype. A clear example is xhNup35-Nb1t, which binds in two copies to a highly conserved epitope of the homodimeric RRM domain of Nup35 (see Fig S4 and Table S1 for the crystal structure of the nanobody·RRM complex). This nanobody stains fully assembled human and frog NPCs (Figs. 4C, 6, 8, S5) and is compatible with the assembly of functional nuclear pores.

**Figure 8.**
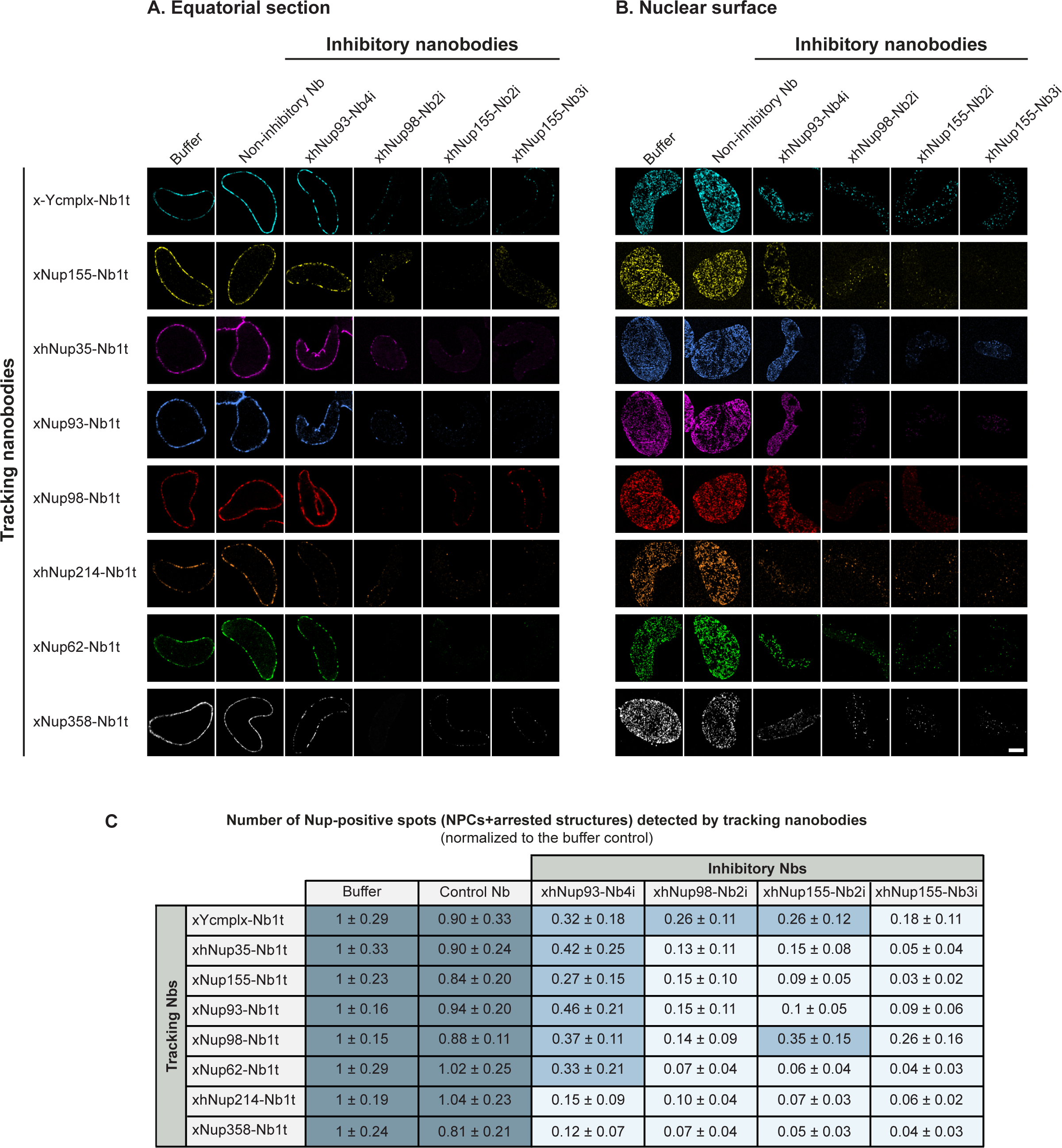
The assembly-inhibited nuclei contain a decreased number of Nup-positive NE structures. Nuclei were assembled as described above, fixed, triton-permeabilized, and incubated with 50 nM of the indicated tracking Nbs. Images were acquired using a ZEISS LSM880 confocal microscope and deconvolved by Airyscan processing. Equatorial planes **(A)** and nuclear surface planes **(B)** of the nuclei are shown. Note that the fixation conditions can alter the morphology and apparent volume of the nuclei; the volume phenotype is thus better appreciated in live samples (*i.e.* Fig 7). Scale bar, 5 µm. **(C)** All inhibitory nanobodies result in the formation of nuclei with a reduced number of NPCs or arrested structures. The number of NPCs and arrested structures was counted from acquired images of the nuclear surface.

The mechanisms by which xhNup93-Nb4i, xhNup93-Nb3i, and xhNup98-Nb2i interfere with NPC biogenesis will be discussed below after mapping their epitopes by structural analyses.

### Assembly-arrested nuclei contain only NPC-like structures in reduced numbers and with derailed Nup compositions

To better characterize the assembly-arrested phenotypes, we repeated the assembly reactions in the presence of the inhibitory nanobodies and stained the resulting (pseudo-) nuclei with tracking nanobodies that recognize different Nups (Fig 8, 9, S5).

**Figure 9.**
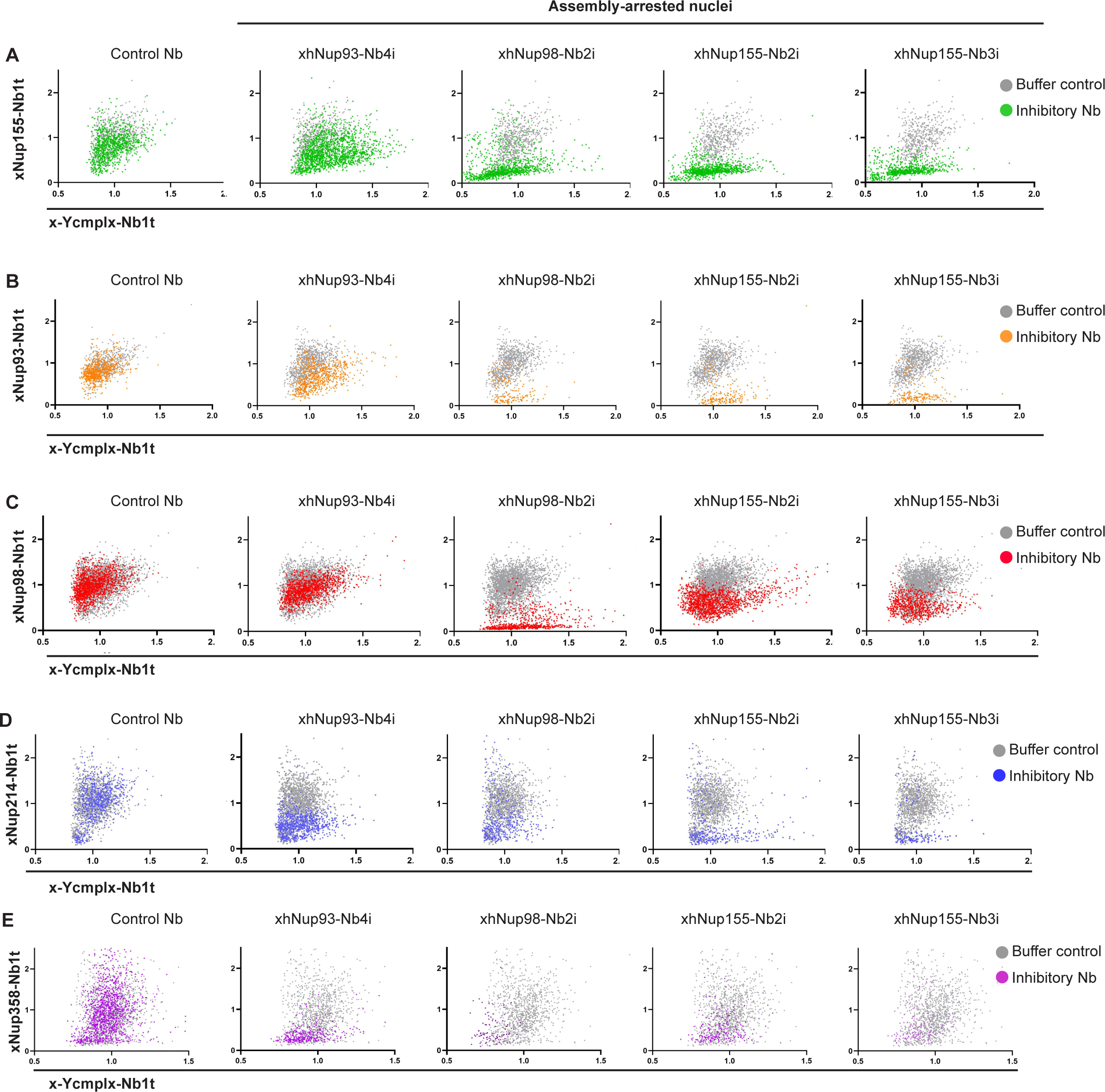
Assembly-arrested pore-like structures contain a derailed and heterogeneous Nup composition that points to off-pathway intermediates. Individual pore-like structures were detected from 3-channel images of assembled nuclei (see Fig 8, S5) using the Y-complex channel (stained by the x-Ycmplx-Nb1t) as the reference channel. Next, the signal intensities of the x- Ycmplx-Nb1t on detected individual spots were plotted on the x axis and the signal intensities of the same spots detected by a different tracking nanobody were plotted on the y axis. Pore-like structures assembled in the presence of the indicated inhibitory nanobodies (labelled in different colours according to the tracking nanobody on the y axis) were overlayed with control nuclear pores assembled in the presence of buffer (labeled in gray). In most cases, assembly-arrested nuclei show a lower intensity signal on the y axis tracking nanobody as compared to the control, with some outliers of higher intensity that might correspond to off-pathway intermediates. Therefore, the inhibitory nanobodies not only decrease the number of assembled pore-like structures as detected by the different tracking nanobodies (see Fig 8C), but also cause a derailed recruitment of the different Nups. Importantly, plots only show pore-like structures in which the Y-complex was detected; thus, potential structures with a low Y-complex signal and a high signal for any of the other Nups are missing from the analysis. Data points were plotted using GraphPad Prism, version 9.

Nuclei arrested by the inhibitory nanobodies showed clearly reduced Nup-signals at the nuclear rim. This phenotype was more dramatic in nuclei arrested by the early-stage inhibitory nanobodies (*i.e.,* xhNup98-Nb2i, xhNup155-Nb2i, and xhNup155-Nb3i) than in those arrested by the xhNup93-Nb4i (Fig 8). A reduced Nup signal on the nuclear surface could be due to either a reduced number of fully-assembled NPCs and/or an unchanged number of pore-like structures containing fewer Nups. To distinguish between these scenarios, we acquired confocal planes of the nuclear surface to resolve individual pores (Fig S5, 9).

Compared to the control, all assembly-arrested nuclei contained a moderately (anti-Nup93- arrested) to dramatically (anti-Nup98- and 155- arrested) reduced number of pore-like structures as stained by x-Ycmplx-Nb1t (Fig S5). Since the Y-complex is the first NPC component to be recruited to assembly sites (Harel et al., 2003b; Otsuka et al., 2018), this indicates that all inhibitory nanobodies reduced the number of initiated assembly events.

### Inhibitory nanobodies targeting distinct Nup155 domains block NPC scaffold assembly

xhNup155-Nb2i recognizes the N-terminal β-propeller of Nup155, wheras xhNup155-Nb3i recognizes its α-solenoid domain (Fig 2A, S6). These two inhibitory nanobodies are not only orthogonal to each other but also orthogonal to our tracking nanobody (xNup155-Nb1t; Fig S6). This implies that xNup155-Nb1t can also track nanobody-inhibited Nup155 molecules.

The two inhibitory nanobodies blocked the incorporation of Nup155 into Y-complex-positive structures in the NE (Fig 8, 9, S5) and prevented the assembly of transport-competent NPCs (Fig 7). Nup155 interacts with Nup160 of the Y-complex, Nup205, and unstructured regions of Nup98 (R3), as well as of Nup35. In addition, it forms an interaction hub with NDC1 and Aladin that anchors the inner ring to the nuclear membrane (Lin et al., 2016; Fontana et al., 2022; Mosalaganti et al., 2022). So far, we cannot tell which of these interactions are directly blocked by the nanobodies. However, their strong phenotypes reaffirm the central role of Nup155 in scaffold assembly and demonstrate that both its β-propeller and its α-solenoid domain are required.

Even though the assembly of functional NPCs was fully blocked, it was evident that some larger NPC sub-complexes of varying composition still assembled in the presence of the inhibitory nanobodies (Fig 9, S5). For example, we observed objects that stained bright for Nup358, the Y- and Nup214 complexes but lacked Nup155. Others stained positive for Nup35, Nup93, and the Y-complex. These rudimentary structures alter the correlation between the intensity of the Y-complex and later-recruited Nups in the imaged pore-like structures (Fig 9) and likely represent off-pathway intermediates that are heterogeneously assembled. Their occurrence illustrates that NPC building blocks have an intrinsic affinity for each other and suggests that proper NPC assembly is kinetically controlled – probably by cooperative binding events.

### The inhibitory anti-Nup93 nanobody impedes the assembly of the NPC cytoplasmic ring

xhNup93-Nb4i reduced the number of assembled NPCs to ∼30% and altered their composition. The Y-complex, Nup35, Nup93, Nup155, the Nup62 complex, and Nup98 still co-localized at approximately wild-type ratios. In contrast, the Nup214·Nup88·Nup62 complex and Nup358 were detected only in some pore-like structures, with clearly reduced intensities (Fig 8, 9, S5). Since the Nup214 complex and Nup358 reside exclusively on the cytoplasmic side of the NPC, this points to defects in the assembly of the cytoplasmic ring.

The xhNup93-Nb4i epitope is only poorly accessible in properly assembled NPCs (Fig 6A, S3), but it becomes accessible in Nup358-depleted NPCs (Fig 10A). Thus, there is indeed an antagonism between this nanobody and Nup358 incorporation.

**Figure 10.**
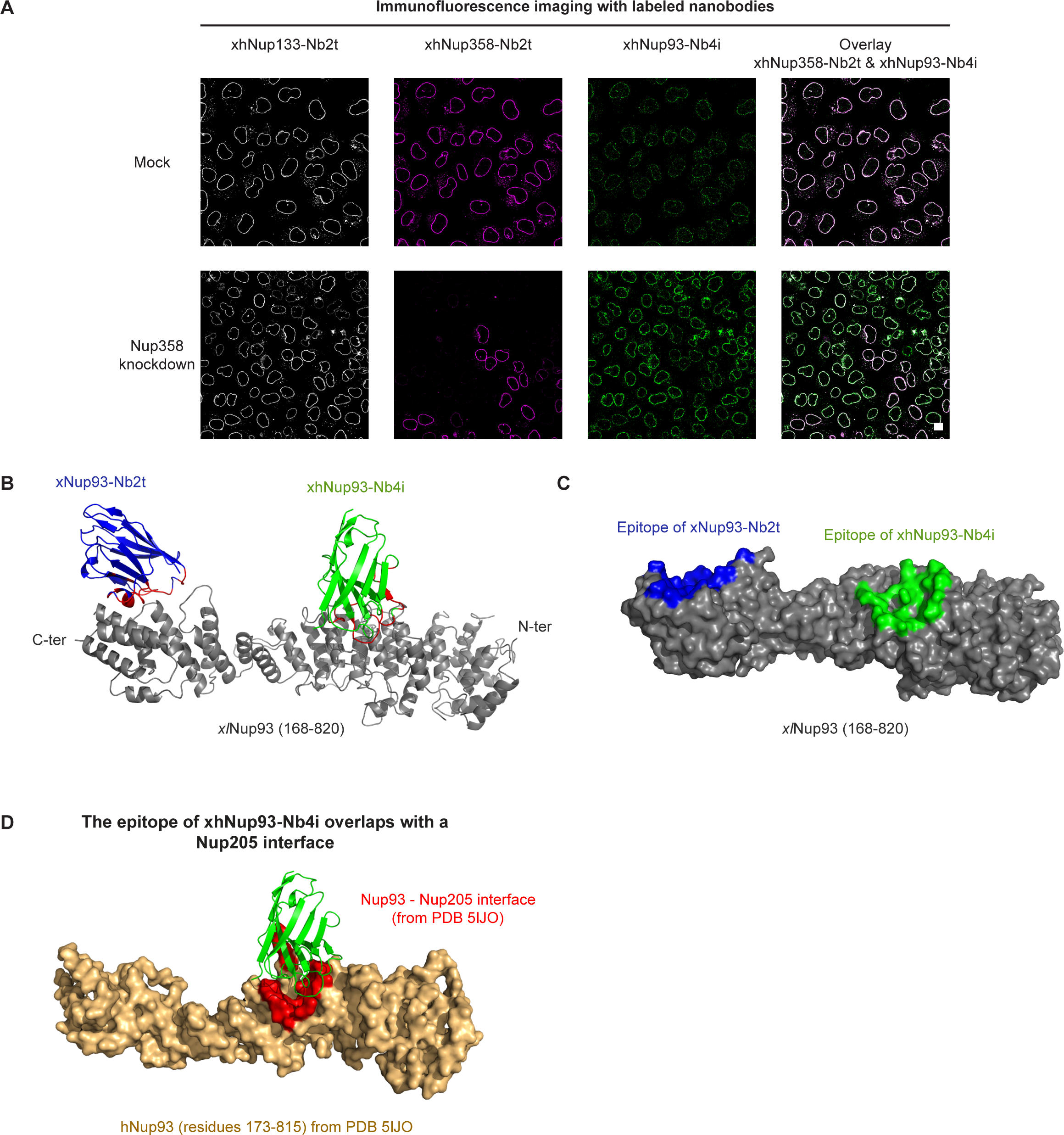
xhNup93-Nb4i recognizes a conserved epitope in the α-solenoid domain of Nup93 and clashes with Nup205 from a neighbouring Nup93·Nup205 dimer. **(A)** The epitope of xhNup93-Nb4i becomes more exposed in NPCs depleted of Nup358. HeLa cells were transfected with siRNAs targeting Nup358, fixed and stained with the indicated tracking nanobodies. Scale bar, 10 µm. **(B)** Cryo-EM structure of *Xenopus laevis (xl)*Nup93(168-820) (gray) in complex with xhNup93-Nb4i (green) and xNup93-Nb2t (blue). The paratope residues of the nanobody are highlighted in red. **(C)** A representation of the Nup93 surface with the nanobody epitopes highlighted in green and blue, respectively. **(D)** xhNup93-Nb4i blocks the interface with a neighboring Nup205 molecule (orange). See Methods for details of crystallization, cryo-EM, and structure solving as well as Tables S2 and S3 for crystallographic statistics.

To map the functional epitope of xhNup93-Nb4i, we solved its structure in complex with the α- solenoid domain of *xl*Nup93(residues 168-820) and xNup93-Nb2t by x-ray crystallography and cryo-EM (Fig 10B-D; Tables S2, S3). xNup93-Nb2t binds to the C-terminal end of the *Xenopus* Nup93 α-solenoid, it is frog-specific (Fig 2A) and did not inhibit NPC assembly. In contrast, xhNup93-Nb4i recognizes an epitope in the middle of the Nup93 α-solenoid (*i.e.,* residues 450- 536). Aligning our structure with recent cryo-EM maps of the NPC revealed that the xhNup93- Nb4i clashes with a Nup205 molecule from a neighboring Nup93·Nup205 heterodimer (Fig 10 D), which would be a plausible explanation for a failed assembly of the cytoplasmic ring (Lin et al., 2016; Bley et al., 2022; Mosalaganti et al., 2022). This consideration further suggests that the occasional Nup214 and Nup358 signals seen in some of the xhNup93-Nb4i distorted NPCs represent off-pathway intermediates (Fig 9, S5).

The failure to recruit Nup358 and the Nup214·Nup88·Nup62 complex is consistent with the observed functional deficits (Figs. 7-9, S5). The Nup214 complex directly contributes two FG domains (from Nup214 itself and from Nup62) to fully-assembled NPCs. In addition, its Nup88 β-propeller anchors a subset of Nup98 molecules, whose FG domains would also not be available. Furthermore, Nup358 is normally present in 40 copies, each contributing several FG sub-domains (Kosinski et al., 2016; Bley et al., 2022; Mosalaganti et al., 2022). Therefore, the anti-Nup93 nanobody-distorted NPCs suffer a cumulative loss of ∼5 megadaltons of FG mass. This explains the increased passive leakage through the FG-based permeability barrier and the defect in Xpo1/NES-mediated nuclear export (Fig 7), which heavily relies on Nup214 and Nup358 (Hutten and Kehlenbach, 2006).

### Assembly-inhibiting anti-Nup98 nanobodies block Nup98-Nup96/ -Nup88 interactions

Nup98 has a conserved domain structure: the unusually conserved N-terminal FG domain, interrupted by a Gle2-binding (GLEBS) domain, makes a key contribution to the NPC permeability barrier (Bailer et al., 1998; Hülsmann et al., 2012). It is followed by an intervening disordered domain with conserved linear motifs (R1, R2, and R3) and a C-terminal autoproteolytic domain (APD) (Hodel et al., 2002; Bley et al., 2022). The catalytic activity of the APD is used to cleave the initially synthesized Nup98-Nup96 fusion protein, which happens during or immediately after translation (Fontoura et al., 1999). Therefore, the egg extract contains the already cleaved and soluble entities: Nup98 and Nup96, which is a subunit of the Y-complex.

The second APD function is to anchor Nup98 to NPCs – either through the N-terminus of Nup96 (Hodel et al., 2002) or the β-propeller of Nup88 (Griffis et al., 2003; Stuwe et al., 2012a; Bley et al., 2022). We have now characterized three nanobodies against the Nup98 APD. One of them (xNup98-Nb1t) stains intact NPCs, permits normal NPC biogenesis, and allows tracking of the assembly process (Fig 1D; Pleiner et al., 2015). By contrast, xhNup98-Nb2i and -Nbi3 are inhibitory and cause a severe and early NPC assembly defect - comparable to a block of Nup155 (Figs. 7-9, S7). To understand this difference, we solved their crystal structures in complex with the Nup98-APD (Fig 11A, Table S4).

**Figure 11.**
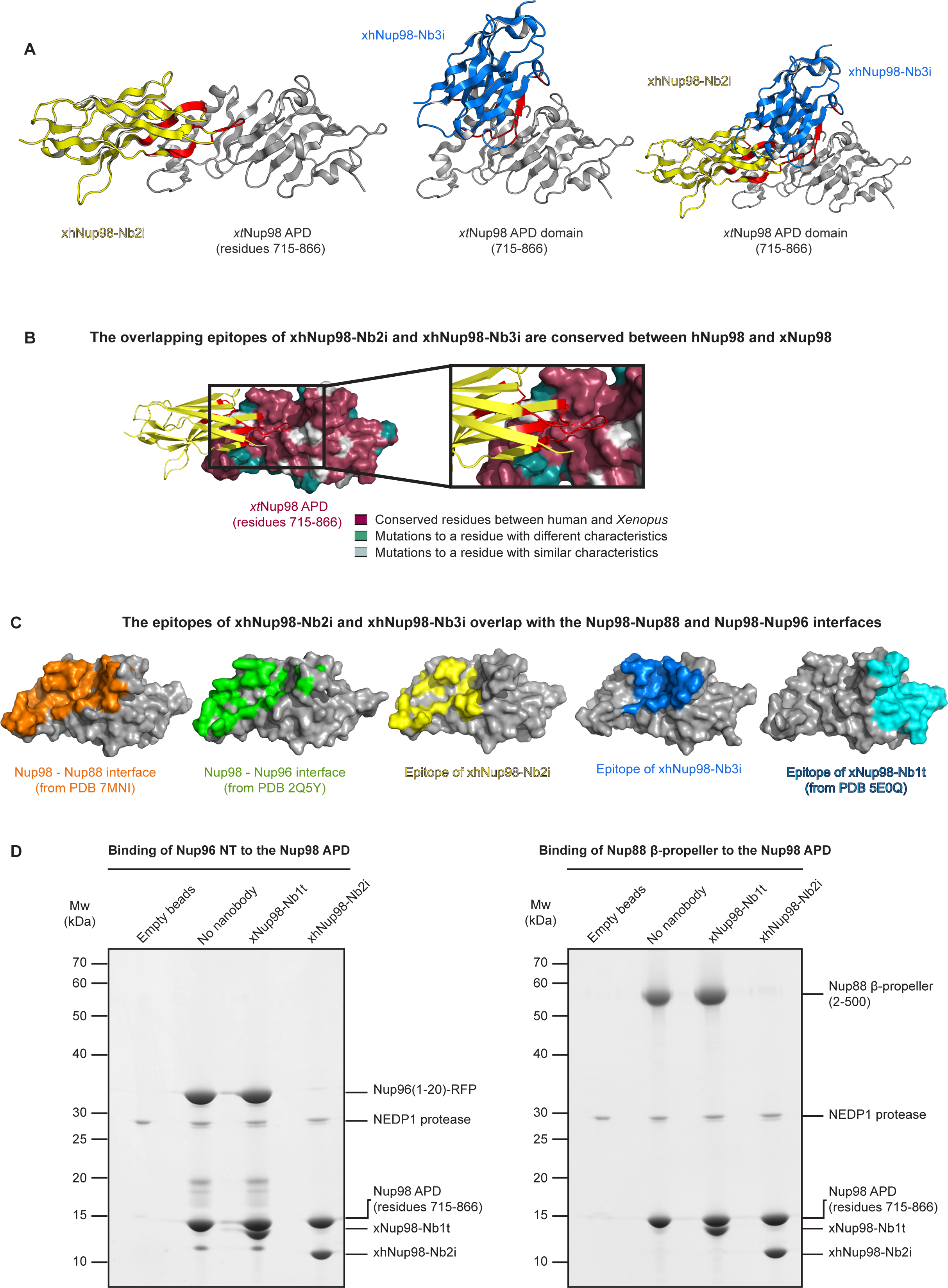
xhNup98-Nb2i recognizes a conserved region of the autoproteolytic domain (APD) of Nup98 and blocks the Nup98-Nup96 and Nup98-Nup88 interactions. **(A)** Crystal structures of the APD of *Xenopus tropicalis (xt)*Nup98 (gray) in complex with xhNup98-Nb2i (yellow) and xhNup98-Nb3i (blue). The epitope-contacting residues of the nanobodies are highlighted in red. Both nanobodies recognize overlapping epitopes. Both nanobodies are equally effective in blocking NPC assembly (see Fig S7 for a comparison). See Methods for details of crystallization and structure solving and Table S4 for crystallographic statistics. **(B)** Surface representation of Nup98 APD color-coded according to amino acid conservation between human and *Xenopus*. The epitope recognized by xhNup98-Nb2i and xhNup98-Nb3i is highly conserved among vertebrates. (**C**) Surface representations of the APD of Nup98 showing its interfaces with Nup88 (orange), Nup96 (green), xhNup98-Nb2i (yellow), xhNup98-Nb2i (blue) and xNup98-Nb1t (cyan). **(D)** The His14-NEDD8 tagged APD of *xt*Nup98(715-866) was immobilized on a Ni^2+^ chelate matrix and incubated with either the xNup98-Nb1t or the xhNup98-Nb2i. Next, either the N-terminus of xlNup96 (aa 1-20) fused to RFP (left) or the N-terminal β propeller of xtNup88(aa 2-500) (right) were added, and the immobilized APD was eluted along with its binding partners by the addition of NEDP1 protease (Frey and Görlich, 2014). Eluted fractions were analyzed by SDS- PAGE/Coomassie-staining.

The obtained structures show that the inhibitory nanobodies xhNup98-Nb2i and -Nbi3 each recognize an overlapping and highly conserved epitope (Fig 11A, B), validating our cross-species phage display strategy to select binders against functionally relevant protein regions. Furthermore, both epitopes overlap with the binding sites of the Nup98-APD for Nup96 and Nup88 (Figs. 11C), suggesting that these nanobodies fully block these interactions. Indeed, xhNup98-Nb2i abolishes the binding of Nup98 to Nup96 and also to Nup88, as tested in a biochemical assay (Fig 11D). By contrast, the tracking nanobody xNup98-Nb1t binds to a non-overlapping epitope on the Nup98- APD and is compatible with both interactions (see structural analysis in Fig 11C and direct interaction assay in Fig 11D). So far, it was assumed that Nup98 gets recruited to a pre-assembled NPC scaffold – the argument being that the bulk of Nup98 arrives at NPC assembly sites rather late (Dultz et al., 2008). Our observation that preventing a Nup98-Nup96 interaction causes a much earlier defect, already at scaffold assembly, suggests a different scenario, namely a tight coupling between scaffold assembly and the establishment of the Nup98-dependent permeability barrier of NPCs.

### xhNup98-Nb2i allows for Nup98 incorporation into already assembled NPCs

To test whether the same phenotype would be observed following the interphase mode of NPC assembly, we assembled *Xenopus* NPCs into an intact human NE. In the presence of the traditional assembly inhibitors or the anti-Nup98 inhibitory nanobodies, new NPC formation was abolished (Fig 12 A). In contrast, the xhNup98-Nb2i nanobody did not prevent the recruitment of *Xenopus* Nup98 from the extract to old human NPCs (Fig 12 B). This can be explained by the fact that the Nup98-APD is not only NPC interaction site. Indeed, cohesive interactions of the Nup98 FG repeats with FG domains of other molecules/FG Nups and of additional linear motifs of Nup98 with Nup205 and Nup155 should all contribute to the NPC anchorage of Nup98 (Griffis et al., 2002a, 2004; Stuwe et al., 2012b; Chatel et al., 2012; Hülsmann et al., 2012; Bley et al., 2022).

**Figure 12.**
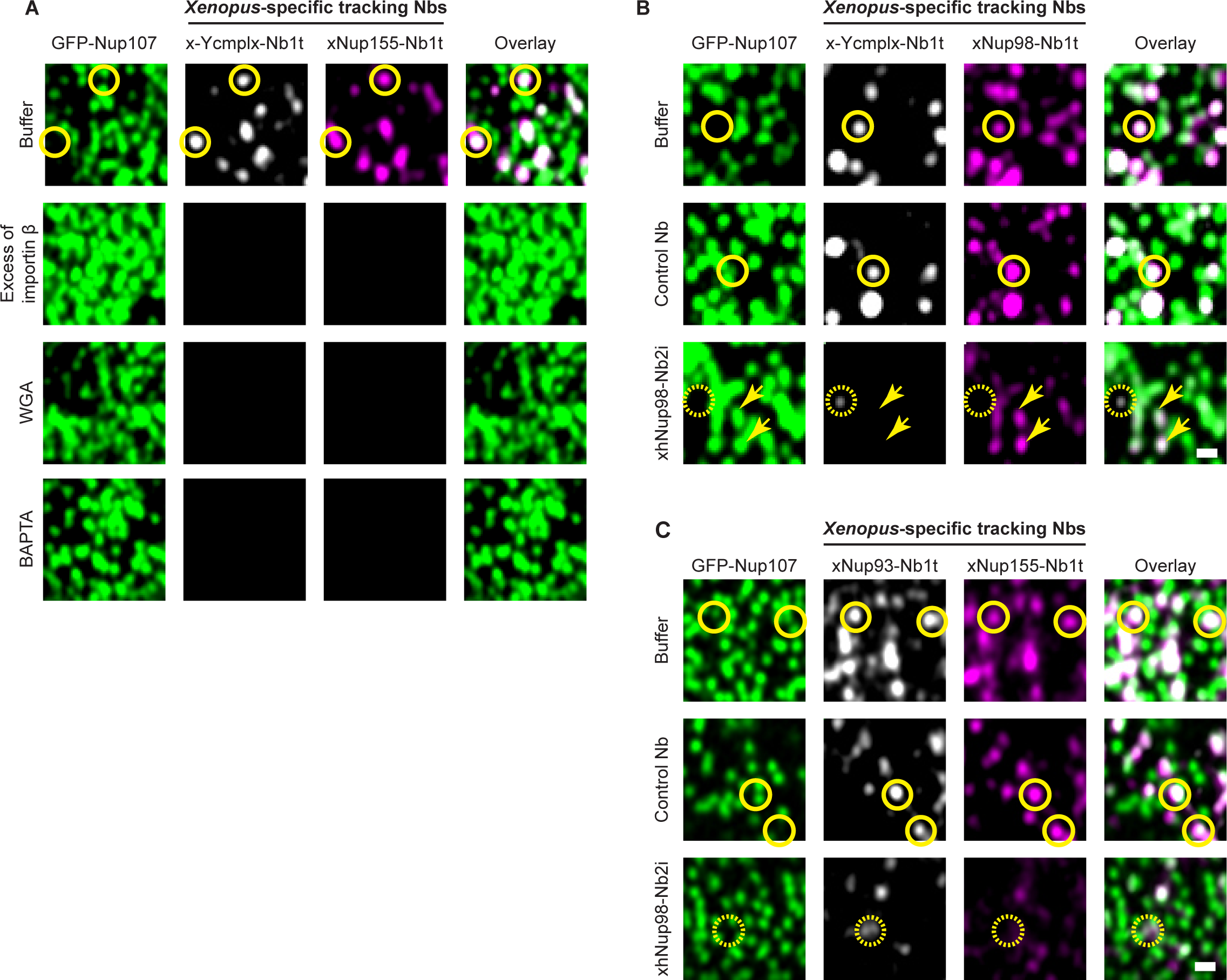
xNup98-Nb2i blocks interphase assembly of new NPCs but allows the exchange of Nup98 into pre-existing pores. New pore insertion was initiated without inhibitor as in Fig 3, or in the presence of 2 µM importin β, 2.5 µM WGA, 5 mM BAPTA or 2 µM of the indicated (unlabeled) inhibitory nanobodies. After 2 hours at RT, cells were washed and imaged using a ZEISS LSM880 confocal microscope. Newly-assembled NPCs are marked by solid lines, incomplete pore-like structures by dashed lines, and pre-existing NPCs by arrows. Scale bar, 0.5 µm. **(A)** An excess of importin β, WGA and BAPTA completely abolished the insertion of new pores. **(B)** xhNup98-Nb2i severely reduced NPC insertion and blocked Nup98 incorporation into new NPC-like structures, while still allowing pre-existing human NPCs to recruit *Xenopus* Nup98. **(C)** xhNup98-Nb2i also reduced the number and intensity of Nup93- and Nup155- containing spots. The control nanobody in **(B)** and **(C)** was xNup93-Nb2t.

Nevertheless, the inhibitory nanobody had the striking effect of blocking the insertion of new NPCs (Fig 12 B). This block appears very early in the assembly pathway, as the appearance of *Xenopus* Y-complex-positive spots at the NE was already suppressed, while the occasionally observed weakly Y-complex-positive structures lacked any Nup98 signal (in contrast to old NPCs). Consistent with this, the detected spots containing other scaffold Nups, such as xNup155 and xNup93, were also fewer in number and weaker in intensity (Fig 12 C). We interpret this as the Nup98 APD mediating interactions that are essential for the assembly of the NPC scaffold. Given the inhibitory effect on the early Y-complex structures, we assume that the APD-Nup96 interaction is the relevant one. This requirement applies to both postmitotic and interphase NPC assembly, suggesting that these two processes are, after all, more similar than currently believed. Given that the final NPC product is the same, this is a plausible scenario.

## Discussion

We present here a toolbox of anti-Nup nanobodies and demonstrate their use as specific trackers or inhibitors of the NPC assembly process. To track post-mitotic assembly, we use the well-established *Xenopus* extract system. The key challenge in tracking interphase assembly is to distinguish newly-assembled from pre-existing NPCs. We solved this problem by using human NEs as templates and inserting chemically distinct frog NPCs from an activated egg extract into them. The described experimental setup opens new avenues to study NPC assembly during interphase and represents a significant advance in the biochemical dissection of this pathway.

Through AddGene, all nanobody plasmids will be made available to the community. Earlier, we reported the first generation of nanobodies directed against frog nucleoporins (Pleiner et al. 2015). We now demonstrate their utility for tracking NPC assembly from *Xenopus* egg extracts, which represents a major simplification of the assay. In addition, we present a new set of tracking nanobodies with broad cross-reactivity that stain *Xenopus* as well as human NPCs. These include nanobodies against Nup358, Nup133, Nup93, Nup35, and the Nup214·88·62 complex. They are suitable for super-resolution microscopy and can resolve NPC rings.

We also report the first NPC assembly-inhibiting nanobodies against three targets: Nup93, Nup155, and Nup98. All of them bind to epitopes that are initially accessible in soluble Nup complexes but become buried in protein-protein interaction interfaces during NPC assembly. The nanobodies thus block critical protein-protein interactions. Inhibitory nanobodies perturb nuclear pore formation by introducing a very small and specific change and are thus excellent tools to complement the structural data of fully-assembled NPCs (Fontana et al., 2022; Bley et al., 2022; Mosalaganti et al., 2022) with insights into the function and assembly mechanisms of the different Nup components. We obtained them by a direct cross-species (frog-human) immunization and phage display strategy for binders of conserved and, thus, functionally relevant epitopes. This strategy is generally applicable and can be used to generate highly selective, protein-based inhibitors of also other molecular machines. For the future, we have planned to systematically extend the anti-NPC toolbox to all relevant subunits, also in the hope of arresting in-pathway-intermediates, such as the immediately pre-fusion pore state of NPCs.

The inhibitory anti-Nup93 nanobody reduced the number of assembly events and imposed a late block that impairs cytoplasmic ring assembly. The resulting NPCs lack Nup358 and the Nup214·88·62 complex and thus suffer a cumulative loss of FG mass, causing defects in the permeability barrier and consequently in active import and export. However, this inhibitory nanobody still allowed rudimentarily functional NPCs.

By contrast, the inhibitory anti-Nup155 nanobodies prevented scaffold assembly, resulting in nuclei without nuclear import activity and an NE containing only a few off-pathway NPC assembly intermediates with derailed Nup compositions. This was not unexpected as Nup155 is a critical architectural element of NPCs. Surprisingly, however, the same scaffold phenotype was caused by anti-Nup98 nanobodies targeting its autoproteolytic domain (APD) and blocking interactions with Nup96 and the Nup88 β-propeller. If these inhibitory anti-APD nanobodies only interfered with the anchoring of Nup98, one would expect that the NPC scaffold (including the Y-complex, Nup93, Nup155, and Nup35) and other FG Nups would still assemble correctly. Instead, they impose an obvious block at early stages of assembly (Figs. 7, 8, S5). This suggests that Nup98 is not just a “late addition” to a pre-assembled NPC scaffold but is already involved in the earliest steps of NPC biogenesis.

This scenario further suggests the existence of multiple Nup98 populations that are recruited at different stages of assembly and through different qualities of Nup-Nup interactions. Some copies of Nup98 are likely recruited at early stages of assembly through obligatory contacts of the Nup98 APD domain with Nup96, engage in additional contacts with other scaffold Nups, and thereby triggering the aforementioned assembly checkpoint for the NPC scaffold. We assume that these Nup98 molecules remain very stably attached.

Indeed, one candidate for such early inter-complex interactions can be prominently detected by affinity chromatography from nuclear assembly extracts (Fig S8), where a Nup98^395-866^ fragment (lacking 394 residues of its FG domain) retrieves not only the Y-complex but also the Nup188·Nup93 complex with similar efficiency. The interaction with the Nup188·Nup93 complex was lost when the remaining 100 residues of the FG domain were also deleted (in Nup98^486-866^). This is consistent with previous reports of GLFG-repeat interactions of the yeast Nup188 homologs (Andersen et al., 2013; Onischenko et al., 2017). In our data, however, this interaction appears to be more sequence-specific and to favor an interaction with Nup188 over the paralogous Nup205. In addition, Nup98 is known to interact through its R3 motif with Nup155 (Lin et al., 2016; Bley et al., 2022; Mosalaganti et al., 2022).

This multitude of interactions renders the APD actually dispensable for steady-state Nup98 anchorage, which explains why the inhibitory anti-APD nanobodies still allow efficient Nup98 incorporation into pre-existing NPCs (Fig 12B). Considering, however, the essential role of the APD during the first steps of NPC biogenesis, it is tempting to speculate that early Nup98-Nup96 interactions are special: they might serve as an assembly checkpoint, allowing scaffold assembly and pore-formation to proceed only after a sufficient number of barrier-forming Nup98 molecules have been recruited, which in turn connect (at least) three key architectural elements of the NPC scaffold: the Y-complex, Nup155 and the Nup188·Nup93 dimer.

The bulk Nup98 population may arrive later in assembly to complete the “sealing” of the NPC central channel with its permeability barrier. This population is likely recruited through fewer contacts and thus weaker interactions – at least this is suggested by the reported mobility of Nup98 in photobleaching experiments (Griffis et al., 2002b) and by the ease with which they are exchanged in the presence of egg extract (Fig 4B). We would assume that the later-arriving population is still heterogeneous, anchored by interactions between the Nup98 unstructured regions and Nup155, Nup188 and/or Nup205, by cohesive interactions between FG domains, and by binding of the Nup98 APD to still vacant Nup96 sites and to the later-recruited Nup88 (Lin et al., 2016; Bley et al., 2022; Mosalaganti et al., 2022).

It can be estimated that a barrier-less NPC with a channel diameter of 60 nm would cause 1000 times more leakage for a GFP-sized molecule than a functional NPC (for derivation, see Ribbeck and Görlich, 2001). For larger species, this leakage ratio would be even higher. This implies that non-selective leakage is dominant and that even a very small fraction of barrier-free NPCs would already deteriorate the performance of the entire nuclear transport system. The Nup98 checkpoint sketched here might be crucial to avoid such a situation, as it would ensure that only NPCs with selective permeability assemble and that NEs with non-selectively open pores are avoided.

## Figure legends

**Figure S1.**
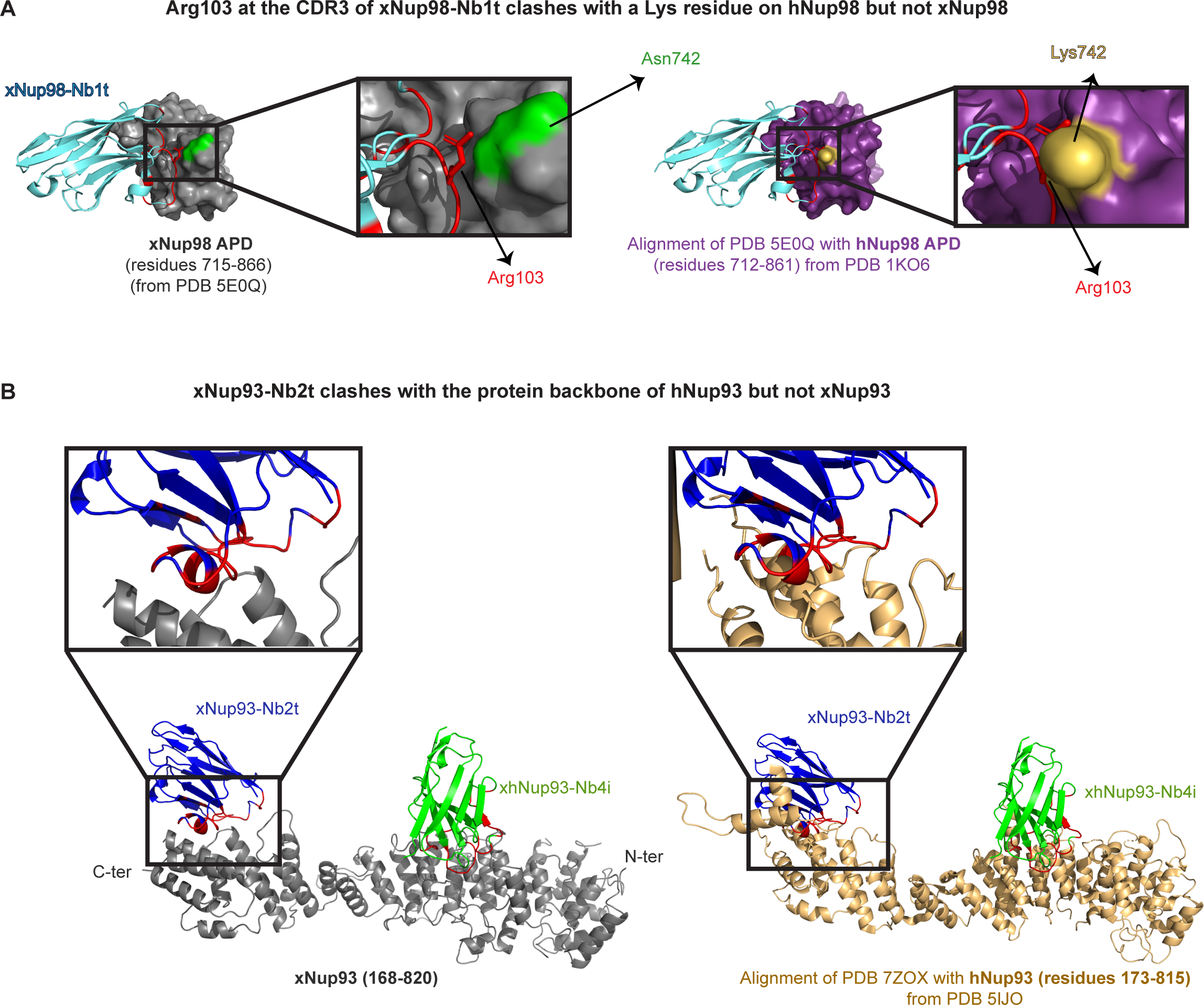
*Xenopus*-specific anti-Nup nanobodies clash with the corresponding human Nup targets. **(A)** The crystal structure of the xNup98-xNup98-Nb1t complex shows that an arginine residue at the nanobody’s CDR3 interacts with an asparagine residue at the APD of *Xenopus* Nup98 (left) (PDB 5E0Q; Pleiner et al., 2015). However, this arginine would clash with a lysine at the same position in hNup98, preventing its binding (right). **(B)** xNup93-Nb2t fits well to a pocket at the C-terminal of *Xenopus* Nup93 (Fig 10), but would clash with the backbone of human Nup93 at the same protein region.

**Figure S2.**
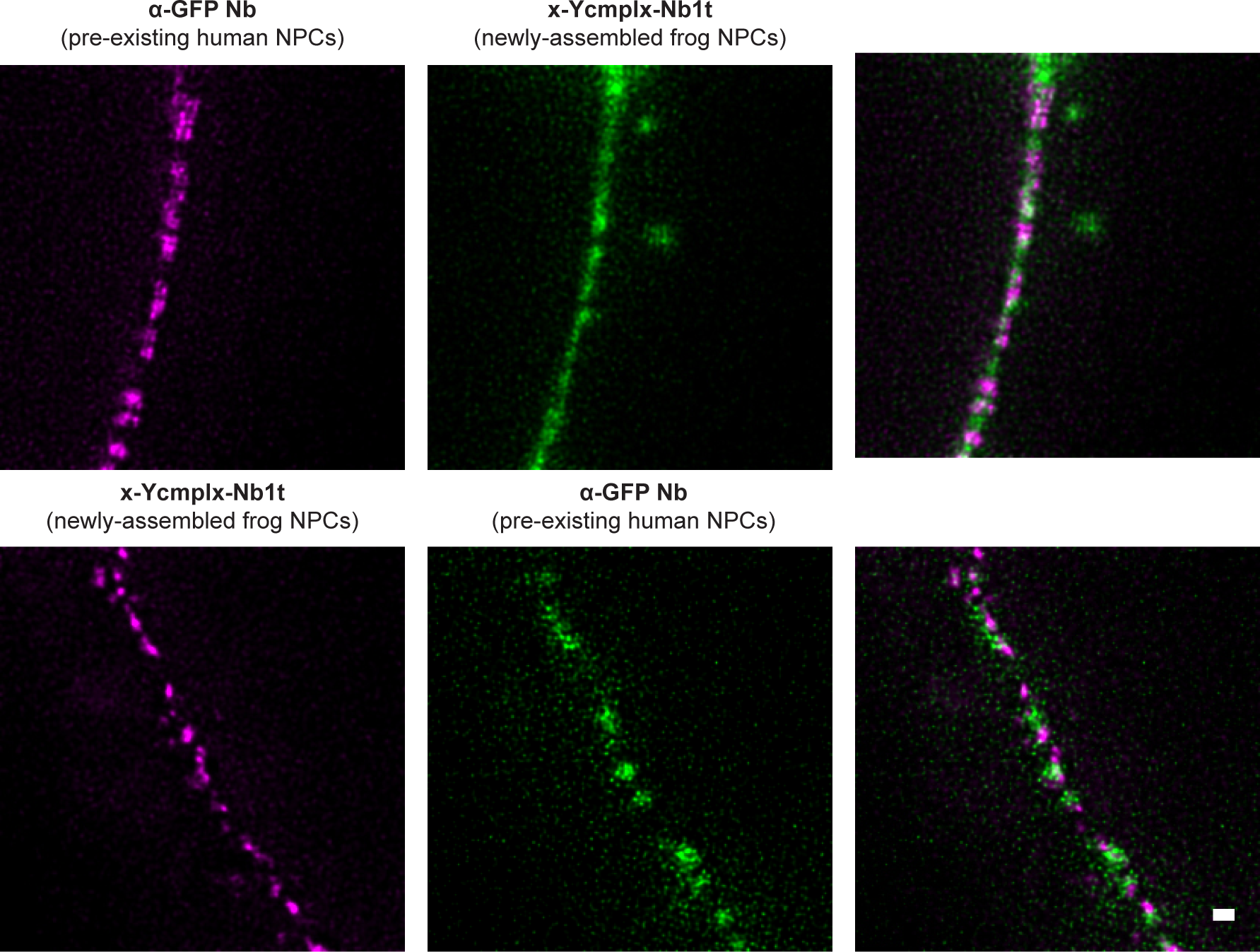
The newly-assembled *Xenopus* NPCs are inserted into human NEs. New pore insertion was initiated as described above, in the presence of an anti-GFP nanobody (Kirchhofer et al., 2010) and the indicated tracking nanobodies labeled with Abberior STAR 580 or Abberior STAR 635P. Next, equatorial planes of the nuclei were imaged using a STEDycon microscope.

**Figure S3.**
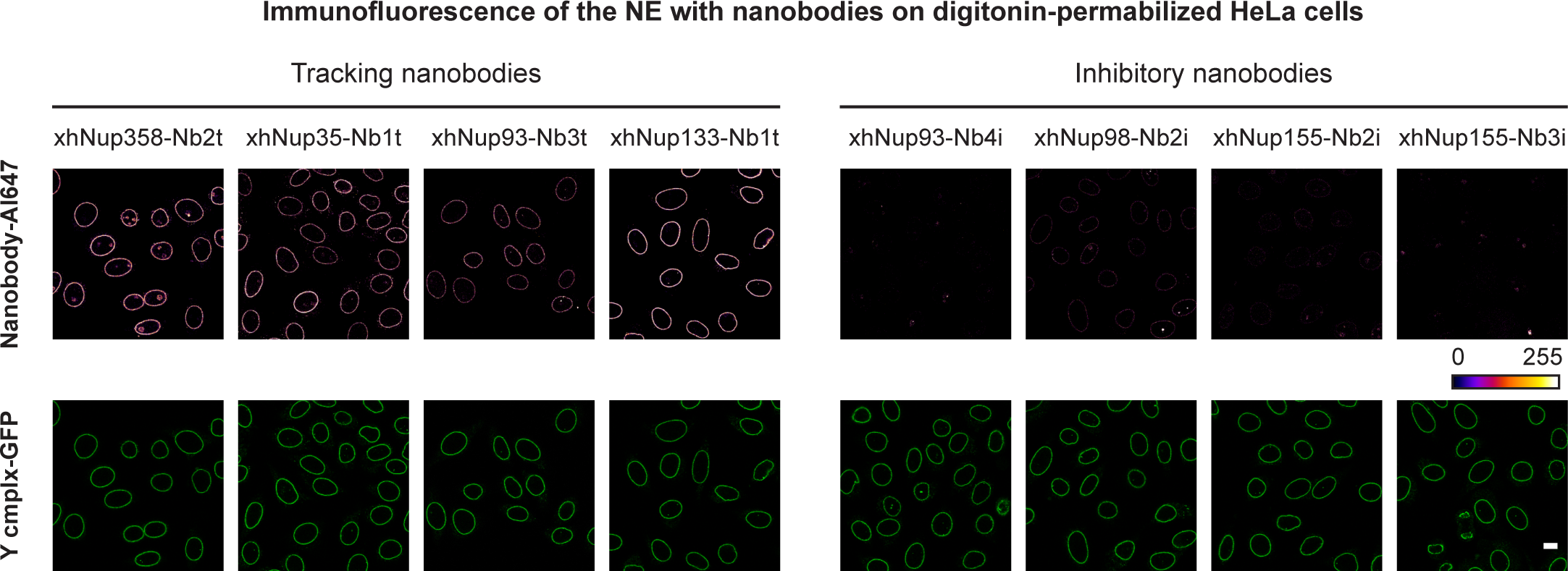
Tracking but not inhibitory nanobodies produced bright and specific fluorescent signals at the NE of digitonin-permeabilized, unfixed HeLa cells. Same experiment as in Fig 6A but analyzing unfixed cells. HeLa cells were permeabilized with digitonin and incubated with 30 nM of the indicated tracking and inhibitory nanobody. After washing off excess of nanobodies, live cells were imaged using a Leica SP8 confocal microscope. All images were acquired using identical microscope settings and are shown as false-color representations (i.e. modified lookup tables (LUTs)) to facilitate a comparison between the different nanobodies. Scale bar, 10 µm.

**Figure S4.**
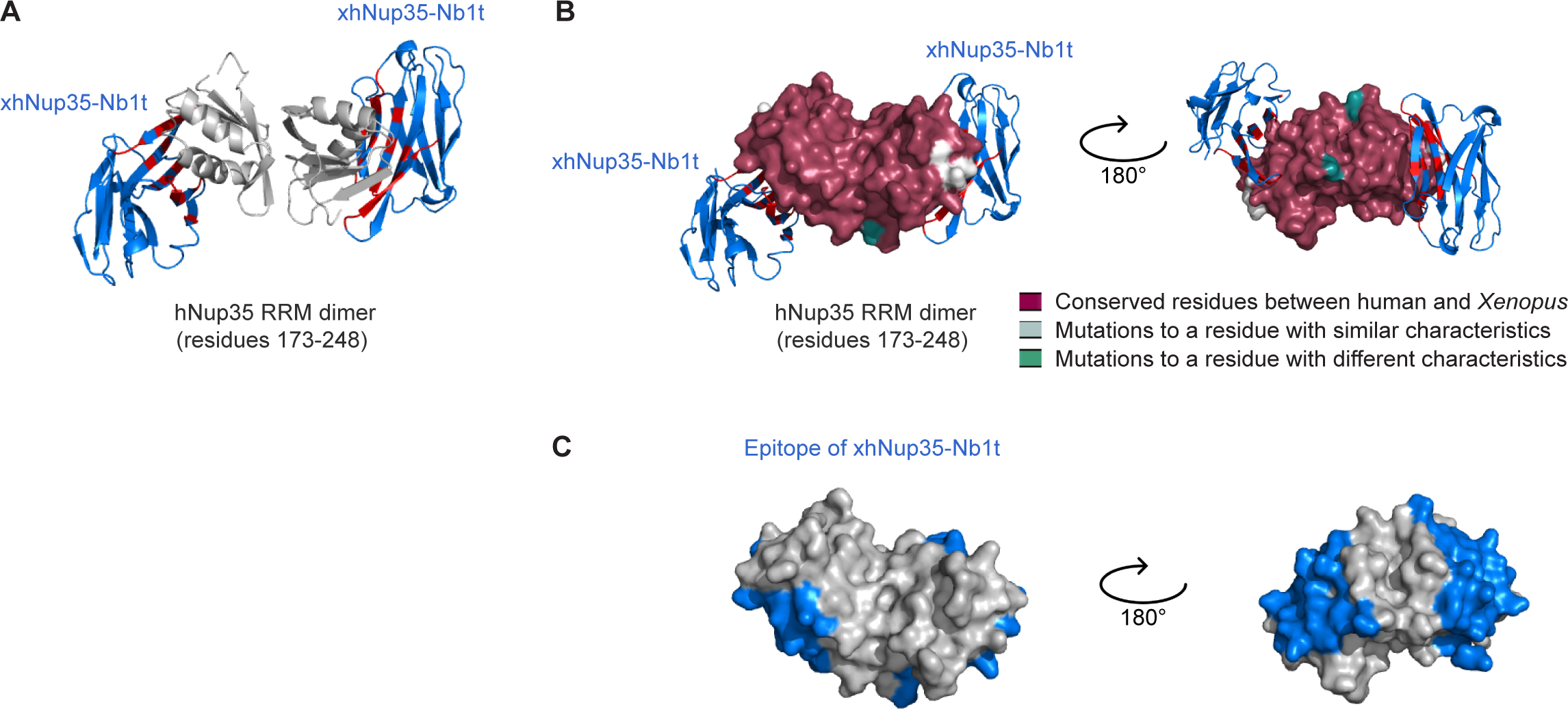
The xhNup35-Nb1t recognizes a conserved epitope at the RRM domain of Nup35. **(A)** Crystal structure of the RRM domain of *homo sapiens (h)*Nup35 (gray) in complex with xhNup35-Nb1t (blue). The nanobody contacting residues are highlighted in red. **(B)** Surface representations of the Nup35 RRM domain color-coded according to amino acid conservation between human and *Xenopus* Nup35. The RRM domain and the nanobody epitope are highly conserved among vertebrates, as also indicated by the nanobody’s cross-reaction between human and *Xenopus* Nup35 (see Fig 4, 6A, B). **(C)** The xhNup35-Nb1t epitope is highlighted in blue to allow for a straightforward comparison with (B). Although the Nup35 RRM domain mediates Nup35 homodimerization, which is necessary for the assembly of functional NPCs (Vollmer et al., 2012), xhNup35-Nb1t recognizes an epitope distant from the dimerization interface and is compatible with the Nup35 dimerization, explaining why it does not interfere with the formation of functional NPCs. In addition, the Nup35 dimer associates with membranes directly or through Ndc1 at very early assembly steps (Vollmer et al., 2012; Eisenhardt et al., 2013), and it links Nup93 to the β-propeller of Nup155, which is essential for the organization of the NPC inner ring and the assembly of the NPC scaffold (Hawryluk-Gara et al., 2008; De Magistris et al., 2018). However, these critical interactions occur through short linear motifs located at the Nup35 disordered termini (Vollmer et al., 2012; Eisenhardt et al., 2013; Mosalaganti et al., 2022), again explaining why xhNup35-Nb1t does not impede NPC assembly. Indeed, the lack of a phenotype is consistent with homodimerization being the only essential function of this RRM domain.

**Figure S5.**
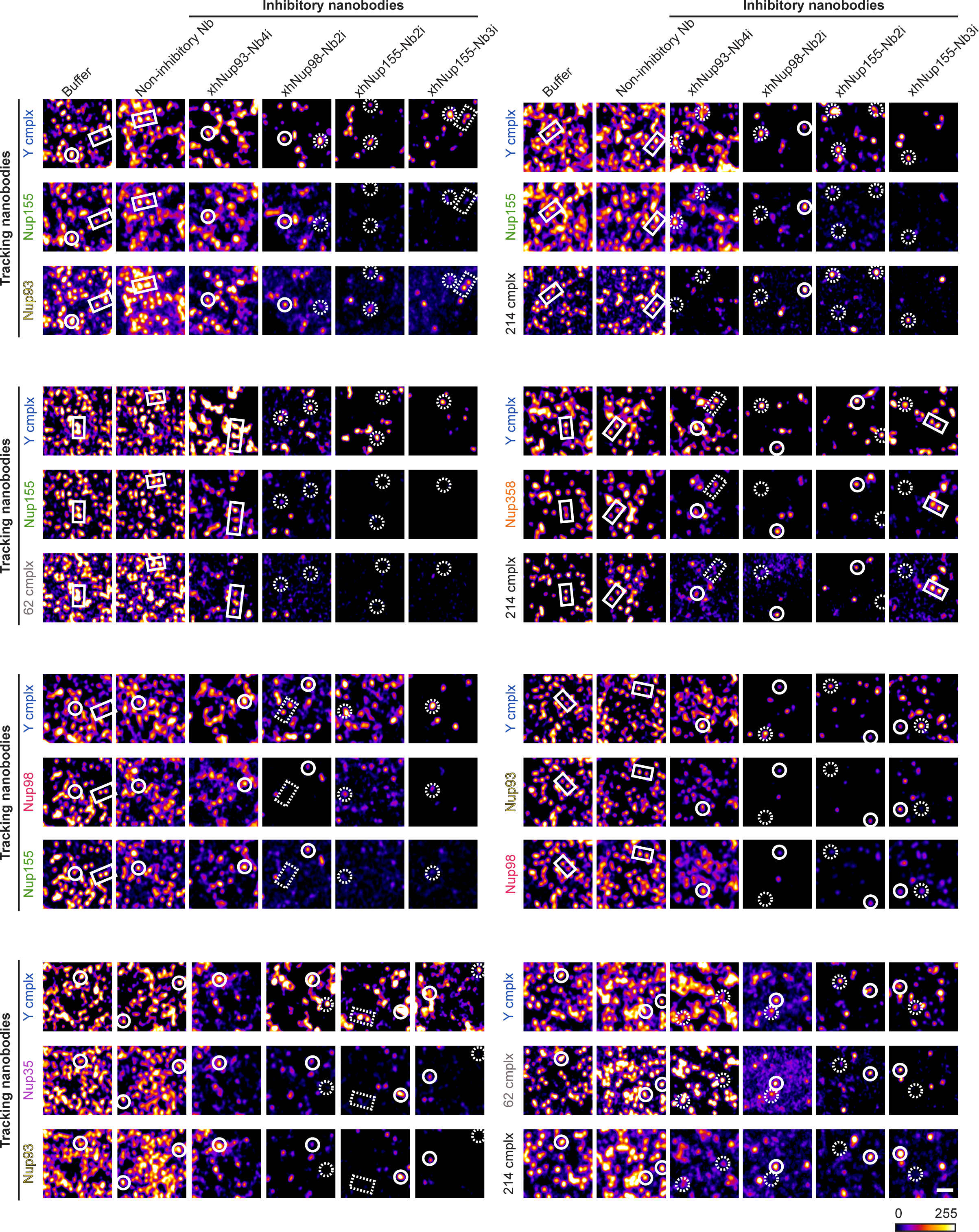
The assembly-inhibited ‘NPCs’ are decreased in number and altered in composition. Representative close-up views of NE assembled in the presence of indicated inhibitory nanobodies. Indicated Nup components were stained by the x-Ycmplx-Nb1t, xNup155- Nb1t, xNup93-Nb1t, xhNup35-Nb1t, xNup98-Nb1t, xNup62-Nb1t, xhNup214-Nb1t or xNup358- Nb1t tracking nanobodies. Pore-like structures in which all probed Nup components were detected are highlighted with white solid lines; arrested structures with missing Nup components are highlighted with white dashed lines. A false-color representation (LUT) is shown to facilitate a comparison between the different images. Scale bar, 1 µm.

**Figure S6.**
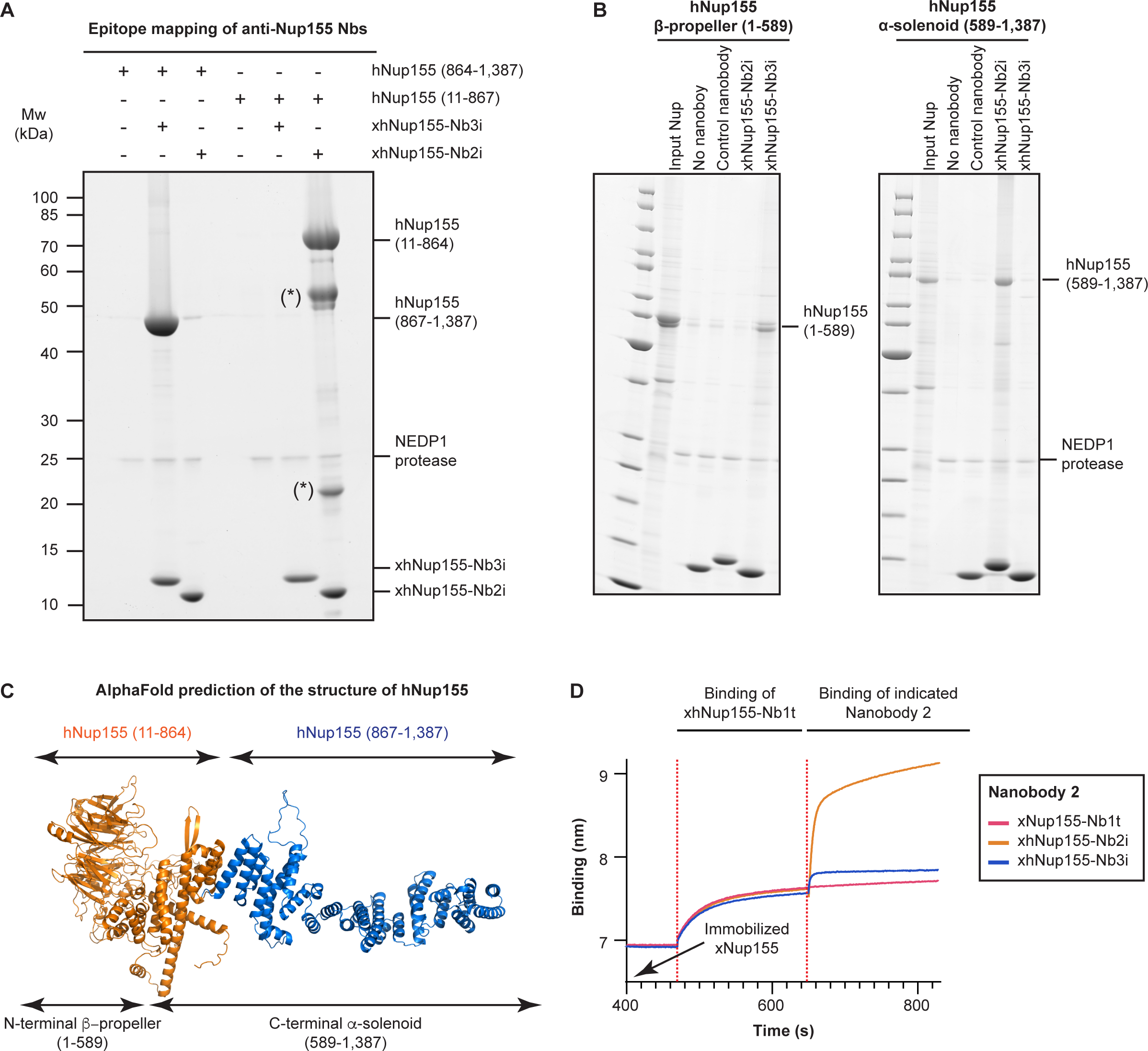
Epitope mapping of the anti-Nup155 nanobodies used in this study. **(A, B)** xhNup155-Nb2i binds to the N-terminal β- propeller domain of Nup155, whereas xhNup155-Nb3i binds to its C-terminal α-solenoid. His14-NEDD8 nanobody fusions were immobilized onto a Ni^2+^ chelate matrix and incubated with the indicated Nup155 fragments. Next, nanobodies were eluted along with their bound targets by the addition of NEDP1 protease (Frey and Görlich, 2014) and the eluates were analyzed by SDS-PAGE/Coomassie-staining. (*) Corresponds to fragments of the N-terminal β-propeller of Nup155 that were spontaneously cleaved during expression/purification from *E. coli*. **(C)** AlphaFold prediction of human Nup155, in which the N-terminal β- propeller and C-terminal α-solenoid domains are colored according to the binding assay in (D). **(D)** Epitope binning experiments by bio-layer interferometry (BLI) confirmed that xhNup155-Nb2i and xhNup155-Nb3i are each orthogonal to xNup155-Nb1t. Biotinylated full-length *xl*Nup155 was loaded onto streptavidin-coated sensors. The loaded sensors were then dipped into wells containing saturating amounts of the xNup155-Nb1t and subsequently into wells containing one of the three different indicated Nbs (Nb2).

**Figure S7.**
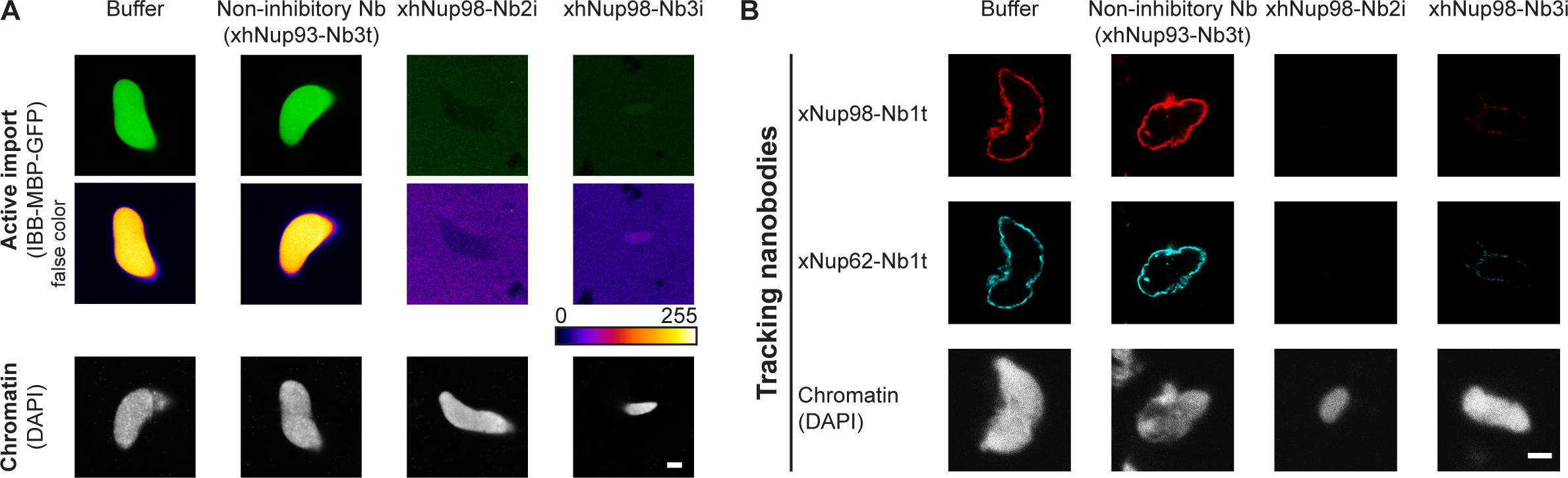
Nanobodies xhNup98-Nb2i and xhNup98-Nb3i interfere with the formation of functional NPCs. As described for xhNup98-Nb2i, the presence of xhNup98-Nb3i during nuclear assembly results in nuclei that fail in active protein import **(A)** and in accumulating NPC markers at the NE **(B)**. This is consistent with the fact that both nanobodies recognize overlapping epitopes that clash with Nup96 and Nup88 (see Fig 11). Since xhNup98-Nb2i an xhNup98-Nb3i cause the same phenotypes, and for space economy, the main manuscript focused on only one of them (xhNup98-Nb2i). Nevertheless, having two independent nanobodies with different paratopes is an excellent control for the specificity of their drastic phenotypes.

**Figure S8.**
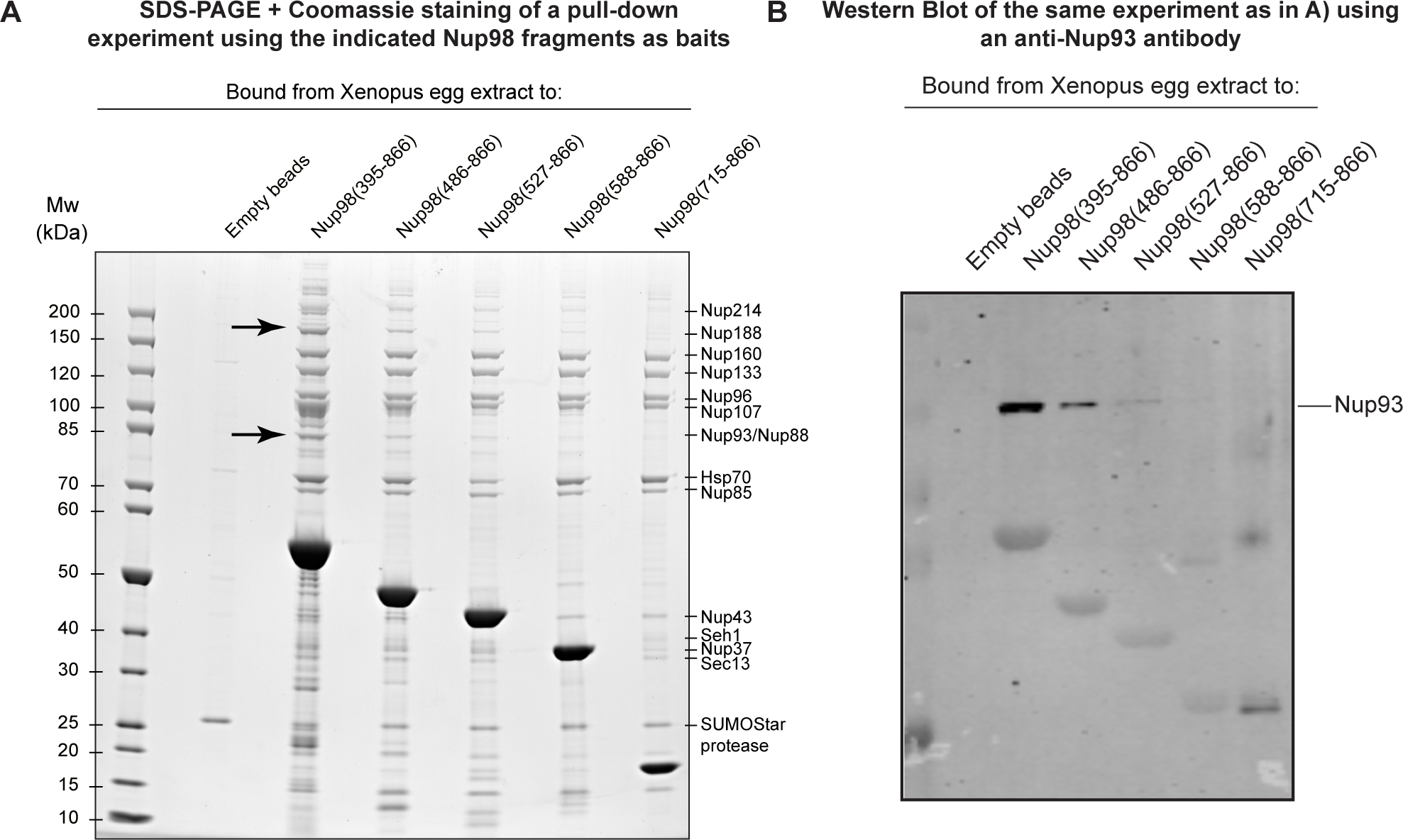
Recombinant Nup98 fragments interact with the Nup188·Nup93 complex from *Xenopus* egg extract. **(A)** The indicated Nup98 truncations were purified from *E. coli* as H14-Avi-biotin SumoSTAR fusions and immobilized on streptavidin-agarose. Next, the soluble fraction from *Xenopus* egg extract was added. After washing, Nup98·Nup complexes were eluted by SumoSTAR protease (Peroutka et al., 2008). Eluted fractions were analyzed by SDS- PAGE/Coomassie-staining. Identity of protein bands was confirmed by mass spectrometry. Bands corresponding to Nup93 and Nup188 are indicated with arrows. **(B)** The binding assay was performed as described in figure (A), but analysis was an immune blot using an anti-Nup93 antibody.

## Materials and methods

### Alpaca immunization

Three female alpacas, held at the Max Planck Institute for Multidisciplinary Sciences, were immunized with 0.5 - 1.0 mg of the human and *Xenopus* orthologues of the respective Nups at 3- 4-week intervals. Nups that had already been immunized by Pleiner *et al*., (2015) were used for two additional immunization boosts, whereas Nups that were used for the first time as immunization antigens were injected 3-4 times.

Immunogens were expressed recombinantly in *E. coli* as H14-NEDD8, His14-scSUMO or His14- SUMO^EU^ fusions, purified by Ni-chelate affinity chromatography with protease elution and subsequent size exclusion chromatography. Next, Nup antigens were buffer-exchanged to a physiological, isotonic, isoosmolar buffer (150 mM NaCl, 20 mM Tris/HCl pH 7.4, 250 mM Sorbitol), mixed with the commercial adjuvant Fama (GERBU Biotechnik GmbH, Cat. No.: 3030) or a squalene-based oil-in-water emulsion, and injected subcutaneously.

### Construction and selections of phage-displayed immune nanobody libraries

The generation of phage-displayed nanobody immune libraries and selection of anti-Nup Nbs with biotinylated baits was performed as previously described (Pleiner *et al*., 2015, 2018). The bait concentration was gradually decreased in subsequent panning rounds to a concentration below 1 nM. Specific nanobody inserts were cloned into H14-NEDD8, His14-scSUMO or His14-SUMO^EU^ expression vectors and transformed into *E. coli* cells. Individual clones were sequenced and representative nanobody sequences were chosen for expression and further characterization.

### Expression and purification of recombinant proteins in *E. coli*

Recombinant Nups were expressed in NEB Express *E. coli* cells (New England Biolabs, C2523). Nanobodies were expressed in *E. coli* SHuffle cells (New England Biolabs, C3026J) for disulfide bond formation and higher stability (Lobstein *et al*., 2016). Protein expression was induced by 30 - 100 μM isopropyl-β-D-thiogalactoside (IPTG) at 18-30 °C for 4 - 20 hours. Next, cells were harvested, resuspended in resuspension buffer (50 mM Tris/HCl pH 7.4, 300 mM NaCl, 20 mM imidazole, 2 mM DTT), and lysed by sonication. His-tagged proteins were immobilized onto pre-equilibrated Ni^2+^ chelate matrixes, washed with resuspension buffer and eluted by the addition of resuspension buffer supplemented with either 400 mM imidazole or the appropriate protease (500 nM bdNEDP1, 50 nM bdSENP1, or 50 nM SUMO^EU^) (Frey and Görlich, 2014; Vera-Rodriguez *et al*., 2019).

### Nanobody labelling with maleimide dyes

Fluorophore maleimides were conjugated to nanobodies through two or three engineered surface cysteines (including N- and C- terminal ones), as previously described (Pleiner *et al*., 2015, 2018). Briefly, purified nanobodies were supplemented with 15 mM dithiothreitol (DTT) for 15 mins on ice. Next, nanobodies were buffer exchanged to maleimide-labeling buffer (300 mM NaCl, 50 mM potassium phosphate (pH 6.8) and rapidly mixed with a 1.2 x molar excess (over the ectopic cysteines) of the respective maleimide fluorescent dye. Labelling reactions proceeded for 40 mins at 4°C and the excess dye was removed using NAP-5 desalting columns (17-0853-01, GE Healthcare). Nanobodies to be used for confocal microscopy were labelled with Alexa488 (Thermo Fisher, A10254), Alexa568 (Thermo Fisher, A20341) or Alexa647 (Thermo Fisher, A20347), and nanobodies to be used for STED microscopy were labelled with the STED fluorophores Abberior STAR 580 (Abberior, ST580-0003) or Abberior STAR 635P (Abberior, ST635P-0003) (Wurm *et al*., 2012). Labelling efficiencies (degree of labelling; DOL) were determined by UV-vis spectroscopy (at 280 nm and the absorbance peak of the respective fluorophore) and validated by size-shifts on SDS-PAGE.

### Nuclear assembly reactions from *Xenopus* egg extracts (post-mitotic assembly assay) (Figs. 1, 7-9, S5, S7)

*Xenopus* egg extracts were prepared as described in Hülsmann, *et al*., (2012). To initiate an assembly reaction, 10 μl egg cytosol were pre-incubated with energy mix (10 mM creatine phosphate, 0.5 mM ATP, 0.5 mM GTP, 50 μg/ml creatine kinase) and sperm chromatin for 15 minutes at 20 °C to allow for chromatin decondensation. Next, membranes (isolated by high-speed centrifugation during the egg extract preparation) were added.

To follow NPC assembly in real time (Fig 1C, D), 50 nM of the corresponding tracking nanobody was added, and confocal images were acquired at the indicated time points using a confocal LSM880 microscope (Zeiss, Germany). To screen for nanobodies that inhibit the formation of functional NPCs (Figures 7-9, S5, S7), 2 μM of each nanobody candidate was added to the egg cytosol fraction prior to chromatin and membrane addition or at the indicated time point. Alternatively, 2.5 μM WGA, 2 μM importin β, or 5 mM BAPTA were added to the egg cytosol fraction to inhibit NPC assembly. After adding the membrane fraction, nuclear assembly proceeded for 1 hour at 18 °C, without shaking. The formation of NPCs bearing an intact permeability barrier was tested by adding 1 μM IBB-MBP-GFP, 5 μg/ml DAPI, and either active (NES-GFP) or passive (MBP-mCherry) exclusion cargoes at a final 0.5 μM concentration (Fig 7). Transport cargoes were incubated for another 30 mins at 18°C, and the resulting nuclei were analysed using a Leica SP8 confocal microscope (Leica, Germany). Notably, the strength of the inhibition might vary depending on the precise time point in which the inhibitory nanobodies are added. Egg extracts are arrested to interphase and activated by the addition of calcium. For a complete inhibitory effect, nanobodies need to be added at a very early time point and before the first interactions between Nups and Nup sub-complexes occur.

To resolve single pores on acquired images of assembled nuclei by confocal microscopy (Figs. 8, 9, S5), it is critical to ensure maximum epitope occupancy and use imaging medium to minimize bleaching and match the diffraction index of the objective immersion oil. Thus, a slightly modified staining protocol was used: 15 μl assembled nuclei were fixed with 25 μl 2.4 % PFA (w/v) for 5 mins at RT, layered on top of 250 μl gradients (10% glycerol/ 11% sucrose in 100 mM NaCl, 50 mM Tris/HCl (pH 7.4)) and centrifuged onto polylysine-coated coverslips (swingout rotor at 1,400 x g for 4 mins). Purified nuclei were washed with PBS, permeabilized with 0.3% v/v triton X-100 for 3 mins at RT, and blocked with 0.5 % BSA. Next, nuclei were incubated with three tracking nanobodies recognizing different Nups and coupled to compatible fluorophores (*i.e.* Alexa 488, Alexa 568 and Alexa 647) at a final 50 nM concentration for 30 mins on ice. After washing off the excess nanobodies, images were acquired using a confocal LSM880 microscope (ZEISS, Germany) and deconvolved by Airyscan Processing (Huff, 2015).

### Permeabilized cell assay for interphase assembly (Figs. 3-5, 12)

The interphase assembly assay was established using a HeLa cell line CRISPR/Cas9 modified to express Nup107 homozygously tagged with sfGFP at its N-terminus (Gunkel *et al*., 2021). HeLa cells were grown at 37°C in High-Glucose Dulbecco’s Modified Eagle Medium (DMEM) supplemented with 10% fetal bovine serum (FBS) (v/v). One day prior to use, HeLa cells were seeded in a 10-well glass slide at a density such that they would reach 70-80% confluence by the time of the experiment.

GFP-Nup107 HeLa cells were permeabilized by the addition of transport buffer (TRB) (110 mM potassium acetate, 3.5 mM magnesium acetate, 20 mM Hepes/KOH (pH 7.5), 1mM EGTA, 250 mM sucrose) supplemented with 30 μg/ml digitonin for 3 mins at RT, with mild shaking. After washing the digitonin off, a *Xenopus* egg extract mixture was added to the cell wells. *Xenopus* egg extract mixtures contained 15 μl of the soluble fraction of *Xenopus* egg extracts, 1.8 μl energy mix (supplying final concentrations of 10 mM creatine phosphate, and each 0.5 mM ATP and GTP), two tracking nanobodies (recognizing different Nups and conjugated to compatible fluorophores) at a 35 nM final concentration each, and TRB to a final volume of 75 μl. To arrest interphase NPC assembly, final concentrations of 2 μM inhibitory nanobodies, 2.5 mM WGA, 2 μM importin β or 5 mM BAPTA were included. The *Xenopus* egg extract mixture was incubated with the HeLa cells for 2 hours at RT, with mild shaking. Next, cells were washed 3 times with TRB and either directly imaged in transport buffer or fixed by the addition of TRB supplemented with 2.4% (w/v) paraformaldehyde for 5 mins at RT.

For an optimal quality of the acquired microscopy images, fixed cells were permeabilized by the addition of TRB supplemented with 0.3% Triton (v/v) for 3 mins at RT. Cells were then blocked with PBS supplemented with 1% BSA (w/v) and subsequently stained with 35 nM of the same tracking nanobodies for 30 mins at RT, with mild shaking. Next, the excess tracking nanobodies were washed off with TRB and the imaging media SlowFade Gold Antifade Mountant (Thermo Scientific) was added to the cells to protect the dyes from photobleaching. For confocal microscopy imaging (Figs. 3, 4 and 12), a ZEISS LSM880 microscope coupled to an Airyscan detector (ZEISS, Germany) was used. Three-channel images of nuclear cross-sections or 3-channel Z-stacks along the bottom NE with a Δz of 0.2 μm were acquired. Next, acquired images were Airyscan processed with a processing strength of 6 (Huff, 2015). Z-stack projections are shown. For super-resolution microscopy (Figs. 5 and S2), cells were imaged using the STEDycon system and deconvolved using the Huyggens Professional software (version 19.10) (Schoonderwoert *et al*., 2013).

### Immunofluorescence of human NPCs (Figs. 6 A, B, S3)

For staining of human NPCs, the same GFP·Nup107 HeLa cell line as above was used (Gunkel *et al*., 2021). GFP·Nup107 HeLa cells were grown on 10-well glass slides (543078, Greiner Bio One) for 24 hours in DMEM high glucose medium supplemented with 10% (v/v) fetal bovine serum (FBS). For staining without prior fixation, cells were semi-permeabilized by the addition of 30 μg/ml digitonin diluted in transport buffer (110 mM KAc, 3.5 mM MgAc, 20 mM Hepes/KOH (pH 7.4), 1mM EGTA, 250 mM sucrose) for 3 mins at room temperature (RT) (Fig S3). Alternatively, cells were fixed with 2.4% PFA (w/v) and permeabilized with 0.3 % Triton/PBS (v/v) for 3 mins at RT (Fig 6A). In both cases, cells were then blocked with 1% (w/v) BSA for 30 mins at RT and stained with 35 nM of the indicated Alexa 647-labeled tracking nanobodies for 30 mins at RT. Finally, the excess tracking nanobodies was washed off and cells were imaged using a Leica SP8 confocal microscope (Leica, Germany). For STED microscopy (Fig 6B), cells were imaged in SlowFade Gold (Thermo Fisher Scientific) imaging medium to protect the dyes from photobleaching. Images were acquired with a STEDycon microscope (Abberior Instruments) using the 640 nm laser. Acquired STED images were deconvoluted using the Huyggens Professional software as described above.

### Native purification of endogenous Nups using nanobodies (figure 6C)

Biotinylated Nups containing a SENP^EU1^ cleavage site were immobilized to a Sepharose-streptavidin matrix (53113, Thermo Scientific^TM^) and incubated with the soluble fraction of either *Xenopus* egg extracts or HeLa lysates for 40 mins at 4 °C. Next, unspecific binders were washed off and the Nup-nanobody complexes were eluted by the addition of 50 nM SENP^EU1^ (Vera-Rodriguez *et al*., 2019). The eluted proteins were analysed by SDS-PAGE followed by Coomassie staining.

### Binding assays with recombinant Nups and nanobodies (figures 11D and S6 A, B)

Purified nanobodies carrying an N-terminal His-tag and a protease cleavage site were immobilized onto pre-equilibrated Ni^2+^ chelate beads for 45 mins at 4°C. Next, excess nanobody was washed off with washing buffer and equimolar amounts of the untagged Nup target were added for 30 mins at 4°C. Excess Nups were then washed off and nanobodies were eluted along with their bound Nup targets by the addition of the appropriate protease. The eluted protein fractions were analysed by SDS-PAGE followed by Coomassie staining.

### Image analysis of the assembled nuclei (Figs. 7-9)

To measure the volume of the reconstituted nuclei (Fig 7D), DAPI and IBB-MBP-GFP were added onto unfixed nuclei assembly reactions. Next, 2-channel Z-stacks were acquired using a Leica SP8 microscope with a Δz of 0.2 μm, and the acquired Z-stacks were thresholded and segmented using the software KNIME version 4.1.0 (Berthold *et al*., 2009). Segmented images were used to measure the nuclei volume using the “Particle Analyzer” tool of the MorpholibJ Plugin (Legland *et al*., 2016) from FiJi/ Image J (Schindelin *et al*., 2012). Finally, the average and SD of at least 20 nuclei per sample from three independent experiments were calculated.

To quantify the active import, active exclusion, and passive exclusion (Fig 7D), acquired images were filtered using FiJi/ Image J (Schindelin *et al*., 2012), thresholded using the Otsu algorithm (Otsu, 1979) and segmented. Segmented images were then used as a mask to measure the mean pixel intensity inside and outside the nuclei using the “Analyze particles” function from FiJi. Partition coefficients were calculated as the ratio between the mean intensity inside and outside each nucleus. The mean and SD of at least 10 nuclei per sample from 4 independent experiments were calculated. To calculate the total import, the partition coefficient was multiplied by the volume of each nucleus.

To count the number of assembled NPCs (or Nup-positive structures) (Fig 8C) the “Analyze particles” function from FiJi was used on segmented 3-channel images and the spots detected on each channel were counted. For each tracking nanobody, the average and SD of at least 5 images from 3 different experiments were measured.

To analyze the correlation between the signal intensities of different Nups (Fig 9), the x-Ycmplx- Nb1t was used as a reference mask to detect nuclear pores and pore-like structures. Then, the signal intensities of the x-Ycmplx-Nb1t were plotted on the x axis, and the signal intensities of the same detected spot as stained by a different tracking nanobody were plotted on the y axis. On each plot, the signal intensities of each phenotype are overlayed with the signal intensities of the buffer control. The plots were obtained using GraphPad Prism, version 9.

### Crystallization and structure determination of the *h*Nup35·xhNup35-Nb1t complex (Fig S4)

Purified untagged *h*Nup35(173-248, the RRM domain) was incubated with a molar excess of His14-NEDD8- xNup35-Nb1t. Next, the formed complexes were immobilized onto a Ni^2+^ chelate matrix and eluted by the addition of 0.5 µM NEDP1 protease. The obtained complex was further purified by size exclusion chromatography using a Hi-Load Superdex 75 16/60 column ÄKTA column, eluted in 100 mM NaCl, 10 mM Tris/HCl (pH 7.5), and concentrated to 10 mg/ml.

Crystallization conditions were screened by robot-assisted screens performed at the Protein Crystallization facility at the Max Planck Institute for Multidisciplinary Sciences. The optimal crystallization condition was 0.05 M HEPES (pH 6.5); 25% v/v PEG400; 0.05 M NaCl, 0.01 M MgCl2. The crystals were cryoprotected by soaking in reservoir solution with increasing amounts of PEG400 up to 40% for 30 minutes, directly harvested from the robotic plate, mounted in loops and flash frozen in liquid nitrogen for data collection. Data sets were collected at EMBL beamline P14, PETRA III storage ring, DESY, Hamburg, Germany using a EIGER X 16M detector (Dectris). The obtained data were processed with XDS (Kabsch, 2009), *XSCALE* (Diederichs, 2006) and *POINTLESS*/*AIMLESS* (Evans and Murshudov, 2013). Initial phases were obtained by molecular replacement in PHASER (McCoy, 2006) using the model coordinates of PDB 4LIR and 2X1O as references. Automatic model building was done using warp (Langer *et al*., 2008) followed by iterative manual building in *Coot* (Emsley *et al*., 2010). The structure of the complex was refined using REFMAC5 (Murshudov *et al*., 2011). Data collection and final refinement statistics are summarized in Table S1.

### Crystallization and structure determination of the Nup93 nanobodies (Fig 10, S1B)

xhNup93-Nb4i and xNup93-Nb2t were expressed as His14-bdSUMO genetic fusions and purified by Ni^2+^ chelate chromatography and on-column protease cleavage. Nanobodies were further purified by size exclusion chromatography using a Hi-Load Superdex 75 16/60 column ÄKTA column, eluted in 100 mM NaCl, 20 mM Tris/HCl (pH 7.4) and concentrated to 10 mg/ml.

Crystallization conditions were screened by vapor diffusion in sitting drops (100 nl protein + 100 nl reservoir) using a Gryphon liquid handling system (Art Robbins Instruments) at the Protein Crystallization facility at the Max Planck Institute for Multidisciplinary Sciences. The crystallization conditions were 100 mM imidazole (pH 8.0), 30% (w/v) PEG 8000, 200 mM NaCl for xhNup93-Nb4i (PDB-ID 8CDS), and 100 mM CAPS (pH 10.5), 1.2 M sodium phosphate, 0.8 M potassium phosphate, 200 mM lithium sulfate for xNup93-Nb2t (PDB-ID 8CDT) at 20 °C. Crystals were cryo-protected in the precipitant solution containing 15 or 25% (v/v) glycerol, respectively, and plunge-frozen in liquid nitrogen.

Diffraction data were collected at the PXII-X10SA beamline at the Swiss Light Source (SLS, Paul-Scherrer-Institut, Villigen, Switzerland) at 100 K, using an EIGER2 X 16M detector (DECTRIS). The datasets were processed in XDS (Kabsch, 2009) and the Phenix Package (Liebschner *et al*., 2019) was used throughout structure solving. The structures were solved by molecular replacement in Phaser (McCoy, 2006). The search model was generated in Sculptor (Bunkóczi and Read, 2011) with the nanobody Re5D06 (PDB-ID 7ON5) as a structural template (Güttler *et al*., 2021). Iterative cycles of model building/adjustment in Coot (Emsley *et al*., 2010) and refinement in phenix.refine (Afonine *et al*., 2018; Liebschner *et al*., 2019) were performed until the refinements converged at the final models (Table S2). Water molecules were built manually in Coot, where the 2Fo−Fc and Fo−Fc maps (contoured at 1σ or 3σ, respectively) showed a peak within a 2.2−3.5 Å distance to oxygen or nitrogen atoms of the protein molecules or other water molecules/ligands. Secondary structure restraints (with isotropic B-factor refinement) were used in early cycles, and restraints were relaxed in later iterations, for which B-factors of all non-water atoms were refined anisotropically. The coordinates and structure factors have been deposited in the PDB with accession code 8CDS for the xhNup93-Nb4i and 8CD7 for the xNup93-Nb2t.

### Structure determination of the Nup93·xhNup93-Nb4i·xNup93-Nb2t complex by Cryo-EM (Fig 10, S1B)

Purified untagged *xl*Nup93(168-820) was incubated with a molar excess of purified untagged xNup93-Nb2t and His14-bdNEDD8- xhNup93-Nb4i. After immobilization to a Ni chelate matrix and washing off excess components, the equimolar trimeric *xl*Nup93(168-820) xhNup93- Nb4i·xNup93-Nb2t complex was eluted by the addition of NedP1 protease. The eluted complex was further purified by size exclusion chromatography using a HiLoad 26/60 Superdex 200 column in 50 mM NaCl, 20mM Tris/HCl (pH 7.5), 2 mM DTT.

For cryo-EM analysis, the purified complex was diluted to 1.7 mg/ml with 25 mM Tris pH 8.0, 50 mM NaCl, 1mM DTT. Two μl of the diluted complex were applied to freshly glow-discharged R 2/2 holey carbon grids (Quantifoil), which were blotted with a force of 6 for 5 seconds and immediately vitrified by plunge-freezing in liquid ethane using a Vitrobot Mark IV (ThermoFisher) operating at 4 °C and 100% humidity. The samples were initially screened using a Glacios cryo-electron microscope (Thermo Fisher) operated at 200 kV with a Falcon-III direct electron detector (Thermo Fisher). Next, a Titan Krios transmission electron microscope (Thermo Fisher) operating at 300 keV with a K3 Summit direct detector (GATAN) and a GIF Quantum Filter was used to obtain the final cryo-EM data. Data were acquired using SerialEM (Mastronarde, 2005) in EFTEM mode with an energy filter slit width of 20 eV and recorded as counting image stacks of 40 movie frames, with a total electron dosage of 54.41 e^−^/Å^2^ at a magnification of 105,000×, equivalent to a calibrated pixel size of 0.834Å. 13 001 movie stacks were imaged with a defocus range of -1.5 to -2.5 μm. Warp (Tegunov and Cramer, 2019) was used for motion correction, dose weighting, contrast-transfer function estimation, and particle selecting on the fly. RELION 3.1. (Zivanov *et al*., 2020) was used for the rest of the image processing. Particles were extracted in a 128-pixel box, binned by 2 at 1.668 pixels per pixel, and subjected to reference-free two-dimensional (2D) classifications. A *de novo* model was created from the good 2D class averages using the 3D Initial Model function, and it was then lowpass filtered and utilized as a low-resolution reference for two rounds of three-dimensional classification, yielding 202,725 particles. These particles were re-extracted in a box of 256 pixels without binning and subjected to global 3D auto-refinement, yielding a 4.8 Å map. The set of particles was subject to one round of CTF refinement and Bayesian polishing. The final global 3D refinement was carried out using the polished particles, which resulted in a 4.4 Å reconstruction.

To build an initial model for the trimeric complex, the AlphaFold structure of Nup93(AF- Q7ZX96) and the crystal structures of xNup93-Nb2t and xhNup93-Nb4i were rigid-body fitted in the 4.4 Å map using UCSF Chimera (Pettersen *et al*., 2004). The fitted models were subsequently subjected to several rounds of MDFF flexible fitting in Namdinator (Kidmose *et al*., 2019). The final model was modified using Coot (Casañal *et al*., 2020). PHENIX (Afonine *et al*., 2018) was used to refine the model in real space and MolProbity (Williams *et al*., 2018) was used to assess it. The structure details are listed in models Table S3. The coordinates and structure factors of the complex have been deposited in the PDB with accession code 7ZOX and to the EMDB with accession code EMD-14849.

### Crystallization and structure determination of the *xl*Nup98-APD·xhNup98-Nb2i and *xl*Nup98-APD·xhNup98-Nb3i complexes (Fig 11, S1A)

Purified untagged *xl*Nup98(714-864) was incubated with a molar excess of either His14-NEDD8- xhNup98-Nb2i or His14-NEDD8-xhNup98-Nb3i. Next, the formed complexes were immobilized onto a Ni^2+^ chelate matrix and eluted by the addition of 0.5 7M NEDP1 protease. The obtained complexes were further purified by size exclusion chromatography on a Hi-Load Superdex 75 16/60 column ÄKTA column, equilibrated in 100 mM NaCl, 10 mM Tris/HCl (pH 7.5), concentrated to 10 mg/ml, and screened for crystallization.

The optimal crystallization condition was 2.5% (w/v) PEG 6000, 25% (v/v) PEG MME 500, 100 mM Tris-HCl (pH 9). Crystals were flash-frozen in liquid nitrogen without additional cryoprotection. Diffraction data was collected remotely from the beamline PXII at the Swiss Light Source (SLS) (Paul Scherrer Institute, Switzerland) and the structure was solved by molecular replacement using the previously published structure of the Nup98 APD·xNup98-Nb1t complex (PDB: 5E0Q) as a search model. Data collection and final refinement statistics are detailed in models Table S4. The coordinates and structure factors have been deposited in the PDB with accession code 7NQ for the *xl*Nup98(714-864)·xhNup98-Nb2i complex and 7NOW for the *xl*Nup98(714-864)·xhNup98-Nb3i complex.

### Nup silencing in HeLa cells (Fig 10A)

HeLa cells were seeded at a low density on 10-well glass slides (543078, Greiner Bio One). Next day, cells were transfected with 10 nM of either mock or Nup358 (s11774, Ambion) siRNAs pre-mixed with 20 μl of HiPerFect transfection reagent (301704, Qiagen) on free-serum media. RNA silencing proceeded for 72 h at 37 °C, and cells were then fixed, blocked, and stained with combinations of the indicated tracking nanobodies. Stained cells were imaged using a confocal SP8 microscope (Leica, Germany).

### Bio-layer interferometry (BLI) for affinity measurements (Fig 2A)

BLI experiments were performed using an Octet RED96e instrument (ForteBio/Sartorius) and High Precision Streptavidin biosensors at 25°C and using PBS (pH 7.4), 0.02% (w/v) Tween-20 and 0.1% (w/v) BSA as assay buffer. KDs listed in Figure 2A were determined by modifying nanobodies (containing two ectopic cysteines) with Biotin-[PEG]3-maleimide (Iris Biotec), immobilizing them to a binding signal of about 1-3 nm on High Precision Streptavidin sensors, allowing 30 nM of their (Xenopus or human) target proteins to bind for 600 seconds and to dissociate for 600-3000 seconds. Apparent KDs were fitted using the Octet Analysis HT 12.0 software.

### Bio-layer interferometry (BLI) for epitope binning (S6)

For orthogonality assays with anti-Nup155 nanobodies (Fig S6), sensors with immobilized full-length xNup155 (biotinylated) were dipped into wells containing saturating amounts of xNup155- Nb1t (untagged) and subsequently into wells containing either the xNup155-Nb1t, the xhNup155- Nb2i or the xhNup155-Nb3 (untagged). Data were reference-subtracted, aligned, and plotted using GraphPad Prism, version 9.

### Animal welfare

All animal experiments have been prior approved by the responsible authority (LAVES; Niedersächsisches Landesamt für Verbraucherschutz und Lebensmittelsicherheit) with the following file numbers: 33.9- 42502-05-13A351 and 33.9-42502-05-17A220 (for immunizations of and blood collection from alpacas) as well as 33.42502-05/A-005/07 and 33.19-4502-05-181246 (for collection of unfertilized *Xenopus laevis* eggs after hormone stimulation).

### Data availability

The coordinates and structure factors of the here reported six structures have been deposited in the PDB with the following accession codes: 8OZB (xhNup35-Nb1t complexed to the human Nup35 RRM homodimer); 8CDS (xhNup93-Nb4i); 8CD7 (xNup93-Nb2t); 7ZOX (cNup93·xhNup93- Nb4i·xNup93-Nb2t complex); 7NQA (*xl*Nup98-APD·xhNup98-Nb2i complex); and 7NOW (xlNup98-APD·xhNup98-Nb3i complex).

## Acknowledgments

We thank Ulrike Teichmann and her team for alpaca care, immunizations, and blood samplings, Jens Krull and Bianka Mußil for helping in the preparation of the nanobody libraries, Renate Rees for providing the Nup133 and Dr. Mamta Jaiswal for providing the Nup214 complex antigens. We also thank Jens Krull, Dr. Bastian Hülsmann, and Dr. Lareen Gräser for the preparation of *Xenopus* egg extracts. We thank Dr. Philip Gunkel and Dr. Volker Cordes for the GFP-Nup107 cell line and *Xenopus*-specific anti-Nup153 Antibodies, as well as Dr. Antonio Politi (Imaging facility of the institute) for assistance with image analysis. We are grateful to the crystallization and cryo-EM facilities of our institute for their excellent support. We thank Paloma Tarrío Alves for the scientific illustrations. This work received funding from the Max Planck Society, the Boehringer Ingelheim Fonds (BIF Ph.D. fellowship to MSC), and an ERC Synergy grant (StuDySARCOMERE to DG).

---

**Table S1.** X-ray data collection and refinement statistics for the *hs*Nup35(173-248)·xhNup35-Nb1t complex a) Values in parentheses are for the highest resolution cell.

|  |  |
| --- | --- |
| **Structure** | ***hsNup35(173-248)*·*xhNup35-Nb1t*** |
| **PDB data entry** | 8OZB |
| **Space group** | P2221 |
| **Unit cell dimensions** |  |
| a, b, c (Å) | 49.12, 77.22, 127.47 |
| $\alpha$, $\beta$, $\gamma$ (°) | 90.00, 90.00, 90.00 |
| **Resolution (Å)** | 77.02 - 2.09 (2.15-2.09)<sup>a</sup> |
| **Wavelength (Å)** | 0.9763 |
| **R<sub>merge</sub>** | 0.193 (1.89) |
| **R<sub>factor</sub>** | 0.2416 |
| **R<sub>free</sub>** | 0.2579 |
| **<I/σ(I)>** | 8.1 (1.4) |
| **CC<sub>1/2</sub>** |  |
| **Reflections** |  |
| Measured | 363326 (17256) |
| Unique | 29113 (1993) |
| **Completeness** | 98.6 (89.7) |
| **Multiplicity** | 12.5 (8.7) |
| **Number of atoms** | 3053 |
| **Protein/Solvent** | 2983/70 |
| **R.m.s. deviation from ideal** |  |
| Bond lengths (Å) | 0.010 |
| Bond angles (°) | 1.739 |
| **Ramachandran statistics (%)** |  |
| Favored | 96.6 |
| Allowed | 3.2 |
| Outliers | 0.0 |
| **<B-factor> (Å<sup>2</sup>)** | 38.23 |

a) Values in parentheses are for the highest resolution cell.

**Table S2.** X-ray data collection and refinement statistics for the nanobodies xhNup93-Nb4i and x- Nup93-Nb2t.

| **Structure** | **xhNup93-Nb4i** | **xNup93-Nb2t** |
| --- | --- | --- |
| **PDB data entry** | 8CDS | 8CDT |
| **Space group** | P12<sub>1</sub>1 | P6<sub>3</sub> |
| **Unit cell dimensions**<br>a, b, c (Å)<br>$\alpha$, $\beta$, $\gamma$ (°) | 26.88, 42.10, 44.95<br>90.00, 93.06, 90.00 | 99.97, 99.97, 30.71<br>90, 90, 120 |
| **Resolution (Å)** | 30.71–1.40 (1.45–1.40) | 28.94–1.33 (1.38–1.33) |
| **Mosaicity (°)** | 0.572 | 0.512 |
| **R<sub>meas</sub>** | 5.6 (174.9) | 4.6 (241.4) |
| **R<sub>p.i.m.</sub>** | 2.4 (82.6) | 1.1 (56.0) |
| **<I/σ(I)>** | 15.09 (0.77) | 32.02 (1.14) |
| **CC<sub>1/2</sub>** | 99.7 (50.1) | 100 (75.5) |
| **Reflections**<br>Measured<br>Unique | 115156 (9490)<br>18991 (1814) | 804408 (78031)<br>40855 (4036) |
| **Completeness** | 95.37 (89.91) | 99.64 (96.89) |
| **Multiplicity** | 6.1 (5.2) | 19.7 (19.3) |
| **No of reflections**<br>**R<sub>work</sub>**<br>**R<sub>free</sub>** | 14.68 (55.16)<br>18.00 (58.40) | 16.62 (33.09)<br>19.04 (31.97) |
| **R.m.s. deviation from ideal**<br>Bond lengths (Å)<br>Bond angles (°) | 0.008<br>0.98 | 0.014<br>1.50 |
| **Ramachandran statistics (%)**<br>Favored<br>Allowed<br>Outliers | 97.58<br>2.42<br>0.00 | 99.16<br>0.84<br>0.00 |
| **<B-factor> (Å<sup>2</sup>)**<br>Average<br>Protein<br>Ligands<br>Solvent | 24.10<br>22.30<br>37.97<br>36.19 | 28.94<br>26.71<br>52.64<br>43.93 |

**Table S3.** Cryo-EM data collection and refinement statistics of the Nup93·xhNup93- Nb4i·xNup93-Nb2t complex.

|  |  |
| --- | --- |
| **Structure** | **xNup93·xhNup93-Nb4i·xNup93-Nb2t complex** |
| **PDB ID** | 7ZOX |
| **EMDB ID** | EMD-14849 |
| **Data collection and Processing** |  |
| Detector | Gatan K3 |
| Magnification | 105,000 |
| Voltage (Kv) | 300 |
| Camera Mode | Counting |
| Electron exposure (e<sup>-</sup>/Å<sup>2</sup>) | 54 |
| Defocus range (μm) | 1.0-2.5 |
| Pixel size (Å) | 0.834 |
| Symmetry imposed | C1 |
| Initial particle images (no.) | 808,840 |
| Final particle images (no.) | 202,725 |
| Map resolution (Å) | 4.4 |
| FSC threshold | 0.143 |
| Map resolution range (Å) | 4.3-6.5 |
| **Refinement** |  |
| Initial model used (PDB code) | - |
| Model Resolution (Å) | 4.4 |
| FSC threshold | 0.5 |
| Map sharpening *B* factor (Å<sup>2</sup>) | -223 |
| Model composition |  |
| Non-hydrogen atoms | 67,778 |
| Protein residues | 852 |
| Ligand/ion | 0 |
| *B* factors (Å<sup>2</sup>) |  |
| Protein | 131.79 |
| Ligand/ion | - |
| R.m.s. deviations |  |
| Bond lengths (Å) | 0.005 |
| Bond angles (°) | 0.708 |
| **Validation** |  |
| MolProbity score | 2.2 |
| Clashscore | 16.59 |
| EMRinger score | 1.89 |
| Map CC (CC mask) | 0.8 |
| Poor rotamers (%) | 0.14 |
| Ramachandran analysis (%) |  |
| Preferred | 92.34 |
| Allowed | 7.66 |
| Outliers | 0 |

**Table S4.**
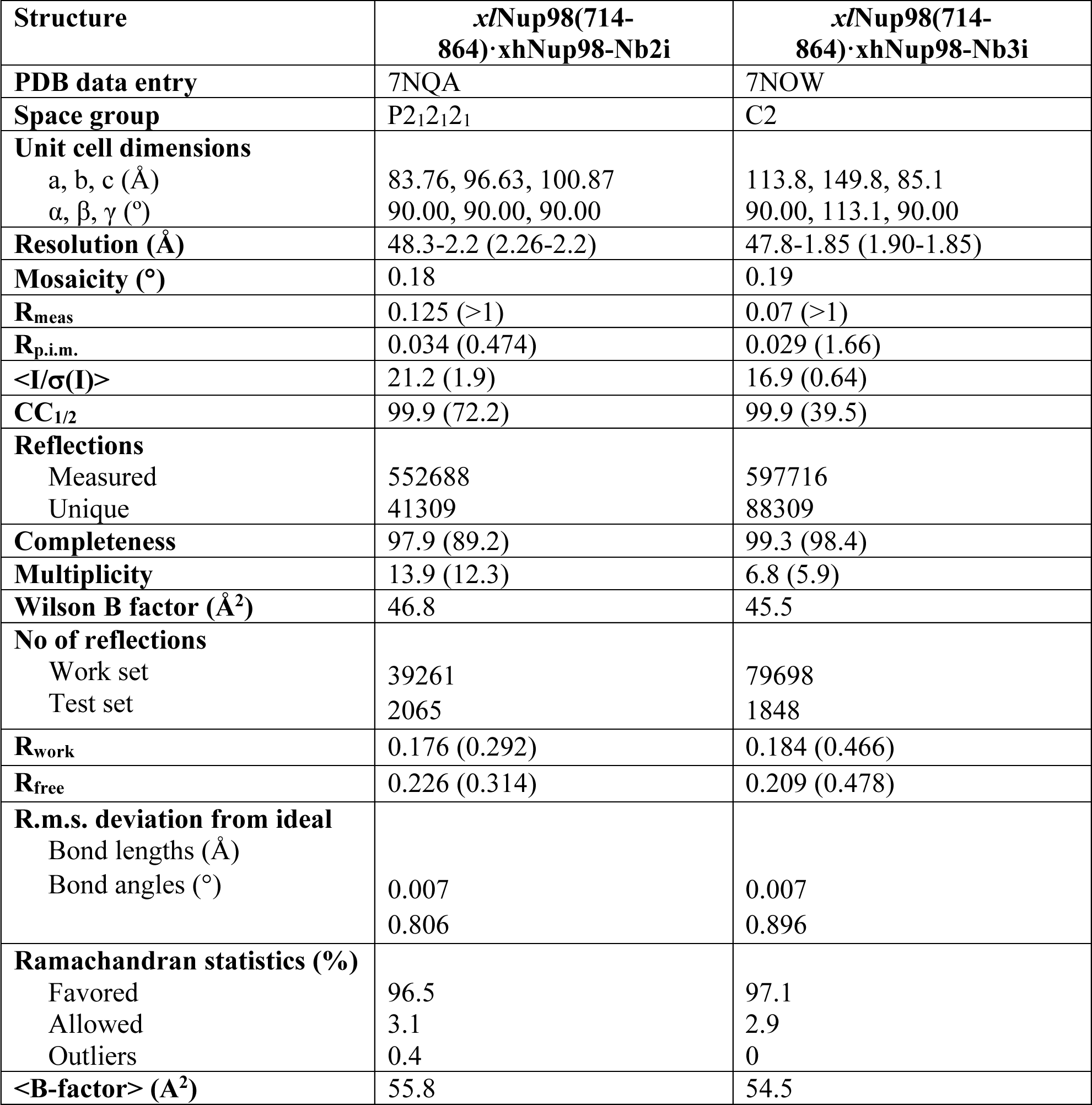
X-ray data collection and refinement statistics for the *xl*Nup98(714-864)·xhNup98-Nb2i and *xl*Nup98(714-864)·xhNup98-Nb3i complexes.

**Table S5.** Plasmid constructs used in this study.

| Plasmid ID | Construct | Use | Addgene entry |
| --- | --- | --- | --- |
| pTP290 | H14-brSUMO-xtNup98(486-715) | Crystallization |  |
| pMSC93 | H14-MBP-brSUMO-Gly-Ser-xlNup93(168-end) | Cryo-EM |  |
| pMSC96 | H14-brNedd8-Gly-anti-Nup93-x-Nup93-Nb1t | Crystallization |  |
| pMSC98 | H14-brNedd8-Gly-Ser-xh-Nup93-Nb4i | Crystallization |  |
| pMSC129 | H14-brNedd8-Gly-xh-Nup98-Nb2i | Crystallization /binding assays/assembly inhibition |  |
| pMSC130 | H14-brNedd8-Gly-xh-Nup98-Nb3i | Crystallization and assembly inhibition |  |
| pMSC100 | H14-brNedd8-Gly-Ser-xh-Nup155-Nb2i | Assembly inhibition and binding assays |  |
| pMSC99 | H14-brNedd8-Gly-Ser-xh-Nup155-Nb3i | Assembly inhibition and binding assays |  |
| pMSC262 | H14-Avi-SUMO<sup>EU1</sup>-xh-Nup93-Nb4i | Assembly inhibition and affinity chromatography |  |
| pMSC264 | H14-Avi- SUMO<sup>EU1</sup>- xh-Nup98-Nb2i | Assembly inhibition and affinity chromatography |  |
| pMSC261 | H14-Avi- SUMO<sup>EU1</sup>-xh-Nup155-Nb3i | Assembly inhibition and affinity chromatography |  |
| pMSC263 | H14-Avi- SUMO<sup>EU1</sup>-xh-Nup155-Nb2i | Assembly inhibition and affinity chromatography |  |
| pTP487 | H14-Sp-brNEDD8-Cys-x-Nup62-Nb1t (S7C)-Spacer-1xCys | Immunofluorescence |  |
| pMSC168 | H14-Sp-brNEDD8-Cys-Sp-xh-Nup35-Nb1t-Spacer-1xCys | Immunofluorescence |  |
| pMSC227 | H14-Nedd8-xh-Nup35-Nb1t (NT, CT, S7 Cys) | Immunofluorescence |  |
| pMSC234 | H14-brNedd8-CGS-xh-Nup93-Nb3t-TSC | Immunofluorescence |  |
| pMSC233 | H14-brNedd8-CGS- xh-Nup93-Nb4i-TSC | Immunofluorescence |  |
| pMSC235 | H14-brNedd8-CGS-xh-Nup155-Nb3i-TSC | Immunofluorescence |  |
| pMSC236 | H14-brNedd8-CGS- xh-Nup155-Nb2i-TSC | Immunofluorescence |  |
| pTP789 | H14-NEDD8-Cys-x-Ycmplex-Nb1t (S7C) | Immunofluorescence |  |
| pTP416 | H14-Sp-brNEDD8-x-Nup93-Nb1t-Spacer-1xCys | Immunofluorescence |  |
| pTP443 | H14-Sp-brNEDD8-Cys-x-Nup98-Nb1t | Immunofluorescence |  |
| pTP489 | H14-Sp-brNEDD8-Cys-x-Nup155-Nb1t (S7C)-Spacer-1xCys | Immunofluorescence |  |
| pMSC226 | H14-Nedd8-xh-Nup358-Nb2t (NT, CT, S7 Cys) | Immunofluorescence |  |
| pTP695 | H14-Nedd8-Cys-x-Nup358-Nb1t (S7C)-Cys | Immunofluorescence |  |
| pDG04163 | acHis-scSUMO-Cys-xh-Nup214-Nb1t-Cys | Immunofluorescence |  |

## References

Afonine, P. V., B.P. Klaholz, N.W. Moriarty, B.K. Poon, O. V. Sobolev, T.C. Terwilliger, P.D. Adams, and A. Urzhumtsev. 2018. New tools for the analysis and validation of cryo-EM maps and atomic models. Acta Crystallogr. Sect. D Struct. Biol. 74:814–840. doi:10.1107/S2059798318009324.

Allegretti, M., C.E. Zimmerli, V. Rantos, F. Wilfling, P. Ronchi, H.K.H. Fung, C.-W. Lee, W. Hagen, B. Turonova, K. Karius, X. Zhang, C. Müller, Y. Schwab, J. Mahamid, B. Pfander, J. Kosinski, and M. Beck. 2020. In cell architecture of the nuclear pore complex and snapshots of its turnover. bioArxiv. preprint:1–35.

Andersen, K.R., E. Onischenko, J.H. Tang, P. Kumar, J.Z. Chen, A. Ulrich, J.T. Liphardt, K. Weis, and T.U. Schwartz. 2013. Scaffold nucleoporins Nup188 and Nup192 share structural and functional properties with nuclear transport receptors. Elife. 2013:1–20. doi:10.7554/eLife.00745.

Antonin, W., J. Ellenberg, and E. Dultz. 2008. Nuclear pore complex assembly through the cell cycle: Regulation and membrane organization. FEBS Lett. 582:2004–2016. doi:10.1016/j.febslet.2008.02.067.

Von Appen, A., J. Kosinski, L. Sparks, A. Ori, A.L. DiGuilio, B. Vollmer, M.T. Mackmull, N. Banterle, L. Parca, P. Kastritis, K. Buczak, S. Mosalaganti, W. Hagen, A. Andres-Pons, E.A. Lemke, P. Bork, W. Antonin, J.S. Glavy, K.H. Bui, and M. Beck. 2015. In situ structural analysis of the human nuclear pore complex. Nature. 526:140–143. doi:10.1038/nature15381.

Bailer, S.M., S. Siniossoglou, A. Podtelejnikov, A. Hellwig, M. Mann, and E. Hurt. 1998. Nup116p and Nup100p are interchangeable through a conserved motif which constitutes a docking site for the mRNA transport factor Gle2p. EMBO J. 17:1107–1119. doi:10.1093/emboj/17.4.1107.

Beck, M., F. Förster, M. Ecke, J.M. Plitzko, F. Melchior, G. Gerisch, W. Baumeister, and O. Medalia. 2004. Nuclear pore complex structure and dynamics revealed by cryoelectron tomography. Science. 306:1387–1390. doi:10.1126/science.1104808.

Beck, M., V. Lŭí, F. Förster, W. Baumeister, and O. Medalia. 2007. Snapshots of nuclear pore complexes in action captured by cryo-electron tomography. Nature. 449:611–615. doi:10.1038/nature06170.

Benton, M.J., P.C.J. Donoghue, R.J. Asher, M. Friedman, T.J. Near, and J. Vinther. 2015. Constraints on the timescale of animal evolutionary history. *Palaeontol*. Electron. 18:1–107. doi:10.26879/424.

Berke, I.C., T. Boehmer, G. Blobel, and T.U. Schwartz. 2004. Structural and functional analysis of Nup133 domains reveals modular building blocks of the nuclear pore complex. J. Cell Biol. 167:591–597. doi:10.1083/jcb.200408109.

Bernis, C., and D. Forbes. 2015. Analysis of Nuclear Reconstitution, Nuclear Envelope Assembly, and Nuclear Pore Assembly Using Xenopus In Vitro Assays. Methods Cell Biol. 33:395–401. doi:10.1038/nbt.3121.ChIP-nexus.

Berthold, M.R., N. Cebron, F. Dill, T.R. Gabriel, T. Kötter, T. Meinl, P. Ohl, K. Thiel, and B. Wiswedel. 2009. KNIME - the Konstanz information miner: version 2.0 and beyond. Assoc. Comput. Mach. 11.

Bley, C.J., S. Nie, G.W. Mobbs, S. Petrovic, A.T. Gres, X. Liu, S. Mukherjee, S. Harvey, F.M. Huber, D.H. Lin, B. Brown, A.W. Tang, E.J. Rundlet, A.R. Correia, S. Chen, S.G. Regmi, T.A. Stevens, C.A. Jette, M. Dasso, A. Patke, A.F. Palazzo, A.A. Kossiakoff, and A. Hoelz. 2022. Architecture of the cytoplasmic face of the nuclear pore. Science. 376:1–51. doi:10.1126/science.abm9129.

Bunkóczi, G., and R.J. Read. 2011. Improvement of molecular-replacement models with Sculptor. Acta Crystallogr. Sect. D Biol. Crystallogr. 67:303–312. doi:10.1107/S0907444910051218.

Casañal, A., B. Lohkamp, and P. Emsley. 2020. Current developments in Coot for macromolecular model building of Electron Cryo-microscopy and Crystallographic Data. Protein Sci. 29:1069–1078. doi:10.1002/pro.3791.

Casterman, C.H., T. Atarhouch, S. Muyldermans, G. Robinson, C. Hamers, E.B. Songa, N. Bendahman, and R. Hamerst. 1993. Naturally occurring antibodies devoid of light chains. 363:446–448.

Cavalier-Smith, T. 1988. Origin of the cell nucleus. BioEssays. 9:72–78.

Chatel, G., S.H. Desai, A.L. Mattheyses, M.A. Powers, and B. Fahrenkrog. 2012. Domain topology of nucleoporin Nup98 within the nuclear pore complex. J. Struct. Biol. 177:81–89. doi:10.1016/j.jsb.2011.11.004.

Cheloha, R.W., T.J. Harmand, C. Wijne, T.U. Schwartz, and H.L. Ploegh. 2020. Exploring cellular biochemistry with nanobodies. J. Biol. Chem. 295:15307–15327. doi:10.1074/jbc.REV120.012960.

Chug, H., S. Trakhanov, B.H. Hülsmann, T. Pleiner, and D. Görlich. 2015. Crystal structure of the metazoan Nup62*Nup58*Nup54 nucleoporin complex. Science. 350:106–110. doi:10.1016/S0005-2736(99)00064-4.

D’Angelo, M.A., D.J. Anderson, E. Richard, and M.W. Hetzer. 2006. Nuclear pores form de novo from both sides of the nuclear envelope. Science. 312:440–443. doi:10.1126/science.1124196.

Diederichs, K. 2006. Some aspects of quantitative analysis and correction of radiation damage. Acta Crystallogr. Sect. D Biol. Crystallogr. 62:96–101. doi:10.1107/S0907444905031537.

Dultz, E., and J. Ellenberg. 2010. Live imaging of single nuclear pores reveals unique assembly kinetics and mechanism in interphase. J. Cell Biol. 191:15–22. doi:10.1083/jcb.201007076.

Dultz, E., E. Zanin, C. Wurzenberger, M. Braun, G. Rabut, L. Sironi, and J. Ellenberg. 2008. Systematic kinetic analysis of mitotic dis- and reassembly of the nuclear pore in living cells. J. Cell Biol. 180:857–865. doi:10.1083/jcb.200707026.

Eisenhardt, N., J. Redolfi, and W. Antonin. 2014. Interaction of Nup53 with Ndc1 and Nup155 is required for nuclear pore complex assembly. J. Cell Sci. 127:908–921. doi:10.1242/jcs.141739.

Emsley, P., B. Lohkamp, W.G. Scott, and K. Cowtan. 2010. Features and development of Coot. Acta Crystallogr. Sect. D Biol. Crystallogr. 66:486–501. doi:10.1107/S0907444910007493.

Evans, P.R., and G.N. Murshudov. 2013. How good are my data and what is the resolution? Acta Crystallogr. Sect. D Biol. Crystallogr. 69:1204–1214. doi:10.1107/S0907444913000061.

Finlay, D.R., and D.J. Forbes. 1990. Reconstitution of biochemically altered nuclear pores: Transport can be eliminated and restored. Cell. 60:17–29. 10.1016/0092-8674(90)90712-N.

Fontana, P., Y. Dong, X. Pi, A.B. Tong, C.W. Hecksel, L. Wang, T.M. Fu, C. Bustamante, and H. Wu. 2022. Structure of cytoplasmic ring of nuclear pore complex by integrative cryo-EM and AlphaFold. Science. 376. doi:10.1126/science.abm9326.

Fontoura, B.M.A., G. Blobel, and M.J. Matunis. 1999. A conserved biogenesis pathway for nucleoporins: Proteolytic processing of a 186-kilodalton precursor generates Nup98 and the novel nucleoporin, Nup96. J. Cell Biol. 144:1097–1112. doi:10.1083/jcb.144.6.1097.

Franz, C., P. Askjaer, W. Antonin, C.L. Iglesias, U. Haselmann, M. Schelder, A. De Marco, M. Wilm, C. Antony, and I.W. Mattaj. 2005. Nup155 regulates nuclear envelope and nuclear pore complex formation in nematodes and vertebrates. EMBO J. 24:3519–3531. doi:10.1038/sj.emboj.7600825.

Franz, C., R. Walczak, S. Yavuz, R. Santarella, M. Gentzel, P. Askjaer, V. Galy, M. Hetzer, I.W. Mattaj, and W. Antonin. 2007. MEL-28/ELYS is required for the recruitment of nucleoporins to chromatin and postmitotic nuclear pore complex assembly. EMBO Rep. 8:165–172. doi:10.1038/sj.embor.7400889.

Frey, S., and D. Görlich. 2007. A Saturated FG-Repeat Hydrogel Can Reproduce the Permeability Properties of Nuclear Pore Complexes. Cell. 130:512–523. doi:10.1016/j.cell.2007.06.024.

Frey, S., and D. Görlich. 2014. A new set of highly efficient, tag-cleaving proteases for purifying recombinant proteins. J. Chromatogr. A. 1337:95–105. doi:10.1016/j.chroma.2014.02.029.

Goldberg, M.W., C. Wiese, T.D. Allen, and K.L. Wilson. 1997. Dimples, pores, star-rings, and thin rings on growing nuclear envelopes: evidence for structural intermediates in nuclear pore complex assembly. J. Cell Sci. 110 (Pt 4:409–20.

Görlich, D., P. Henklein, R.A. Laskey, and E. Hartmann. 1996. A 41 amino acid motif in importin- α confers binding to importin-β and hence transit into the nucleus. EMBO J. 15:1810–1817. doi:10.1002/j.1460-2075.1996.tb00530.x.

Görlich, D., and U. Kutay. 1999. Transport between the nucleus and the cytoplasm. Annu. Rev. Cell Dev. Biol. 15:607–660.

Griffis, E., N. Altan, J. Lippincott-Schwartz, and M.A. Powers. 2002a. Nup98 is a mobile nucleoporin with transcription-dependent dynamics. Mol. Biol. Cell. 13:2170–2179. doi:10.1091/mbc.01.

Griffis, E.R., N. Altan, J. Lippincott-Schwartz, and M.A. Powers. 2002b. Nup98 is a mobile nucleoporin with transcription-dependent dynamics. Mol. Biol. Cell. 13:1282–1297. doi:10.1091/mbc.01.

Griffis, E.R., B. Craige, C. Dimaano, K.S. Ullman, and M.A. Powers. 2004. Distinct Functional Domains within Nucleoporins Nup153 and Nup98 Mediate Transcription-dependent Mobility. Mol. Biol. Cell. 15:1991–2002. doi:10.1091/mbc.E03.

Griffis, E.R., S. Xu, and M.A. Powers. 2003. Nup98 localizes to both nuclear and cytoplasmic sides of the nuclear pore and binds to two distinct nucleoporin subcomplexes. Mol. Biol. Cell. 14:600–610. doi:10.1091/mbc.E02.

Gunkel, P., H. Lino, S. Krull, and V.C. Cordes. 2021. Zc3hc1 is a novel inherent component of the nuclear basket, resident in a state of reciprocal dependence with tpr. Cells. 10:1–40. doi:10.3390/cells10081937.

Güttler, T., M. Aksu, A. Dickmanns, K.M. Stegmann, K. Gregor, R. Rees, W. Taxer, O. Rymarenko, J. Schünemann, C. Dienemann, P. Gunkel, B. Mussil, J. Krull, U. Teichmann, U. Groß, V.C. Cordes, M. Dobbelstein, and D. Görlich. 2021. Neutralization of SARS-CoV- 2 by highly potent, hyperthermostable, and mutation-tolerant nanobodies. EMBO J. 40:1–26. doi:10.15252/embj.2021107985.

Handa, N., M. Kukimoto-Niino, R. Akasaka, S. Kishishita, K. Murayama, T. Terada, M. Inoue, T. Kigawa, S. Kose, N. Imamoto, A. Tanaka, Y. Hayashizaki, M. Shirouzu, and S. Yokoyama. 2006. The Crystal Structure of Mouse Nup35 Reveals Atypical RNP Motifs and Novel Homodimerization of the RRM Domain. J. Mol. Biol. 363:114–124. doi:10.1016/j.jmb.2006.07.089.

Harel, A., R. Chan, A. Lachish-Zalait, E. Zimmerman, M. Elbaum, and D.J. Forbes. 2003a. Importin beta negatively regulates nuclear membrane fusion and nuclear pore complex assembly. Mol. Biol. Cell. 14:4387–4396. doi:10.1091/mbc.E03.

Harel, A., A. V. Orjalo, T. Vincent, A. Lachish-Zalait, S. Vasu, S. Shah, E. Zimmerman, M. Elbaum, and D.J. Forbes. 2003b. Removal of a single pore subcomplex results in vertebrate nuclei devoid of nuclear pores. Mol. Cell. 11:853–864. 10.1016/S1097-2765(03)00116-3.

Helma, J., M.C. Cardoso, S. Muyldermans, and H. Leonhardt. 2015. Nanobodies and recombinant binders in cell biology. J. Cell Biol. 209:633–644. doi:10.1083/jcb.201409074.

Hodel, A.E., M.R. Hodel, E.R. Griffis, K.A. Hennig, G.A. Ratner, S. Xu, and M.A. Powers. 2002. The three-dimensional structure of the autoproteolytic, nuclear pore-targeting domain of the human nucleoporin Nup98. Mol. Cell. 10:347–358. doi:10.1016/S1097-2765(02)00589-0.

Huff, J. 2015. The Airyscan detector from ZEISS: confocal imaging with improved signal-to-noise ratio and super-resolution. Nat. Methods. 12:i–ii. doi:10.1038/nmeth.f.388.

Hülsmann, B.B., A.A. Labokha, and D. Görlich. 2012. The permeability of reconstituted nuclear pores provides direct evidence for the selective phase model. Cell. 150:738–751. doi:10.1016/j.cell.2012.07.019.

Hutten, S., and R.H. Kehlenbach. 2006. Nup214 Is Required for CRM1-Dependent Nuclear Protein Export In Vivo. Mol. Cell. Biol. 26:6772–6785. doi:10.1128/mcb.00342-06.

Ingram, J.R., F.I. Schmidt, and H.L. Ploegh. 2018. Exploiting Nanobodies’ Singular Traits. Annu. Rev. Immunol. 36. doi:10.1146/annurev-immunol-042617-053327.

Kabsch, W. 2009. XDS. Acta Crystallogr. Sect. D Biol. Crystallogr. 125–132. doi:10.1107/s090744499201028x.

Kassube, S.A., T. Stuwe, D.H. Lin, C.D. Antonuk, J. Napetschnig, G. Blobel, and A. Hoelz. 2012. Crystal structure of the N-terminal domain of Nup358/RanBP2. J. Mol. Biol. 423:752–765. doi:10.1016/j.physbeh.2017.03.040.

Kidmose, R.T., J. Juhl, P. Nissen, T. Boesen, J.L. Karlsen, and B.P. Pedersen. 2019. Namdinator - Automatic molecular dynamics flexible fitting of structural models into cryo-EM and crystallography experimental maps. IUCrJ. 6:526–531. doi:10.1107/S2052252519007619.

Kirchhofer, A., J. Helma, K. Schmidthals, C. Frauer, S. Cui, A. Karcher, M. Pellis, S. Muyldermans, C.S. Casas-Delucchi, M.C. Cardoso, H. Leonhardt, K.P. Hopfner, and U. Rothbauer. 2010. Modulation of protein properties in living cells using nanobodies. Nat. Struct. Mol. Biol. 17:133–139. doi:10.1038/nsmb.1727.

Klar, T.A., S. Jakobs, M. Dyba, A. Egner, and S.W. Hell. 2000. Fluorescence microscopy with diffraction resolution barrier broken by stimulated emission. Proc. Natl. Acad. Sci. U. S. A. 97:8206–8210. doi:10.1073/pnas.97.15.8206.

Knockenhauer, K.E., and T.U. Schwartz. 2016. The nuclear pore complex as a flexible and dynamic gate. Cell. 164:1162–1171. doi:10.1016/j.cell.2016.01.034.

Kosinski, J., S. Mosalaganti, A. Von Appen, R. Teimer, A.L. Diguilio, W. Wan, K.H. Bui, W.J.H. Hagen, J. a G. Briggs, J.S. Glavy, E. Hurt, and M. Beck. 2016. Molecular architecture of the inner ring scaffold of the human nuclear pore complex. 352:363–365.

Kutay, U., R. Jühlen, and W. Antonin. 2021. Mitotic disassembly and reassembly of nuclear pore complexes. Trends Cell Biol. 31:1019–1033. doi:10.1016/j.tcb.2021.06.011.

Labokha, A.A., S. Gradmann, S. Frey, B.B. Hülsmann, H. Urlaub, M. Baldus, and D. Görlich. 2013. Systematic analysis of barrier-forming FG hydrogels from Xenopus nuclear pore complexes. EMBO J. 32:204–218. doi:10.1038/emboj.2012.302.

Langer, G., S.X. Cohen, V.S. Lamzin, and A. Perrakis. 2008. Automated macromolecular model building for X-ray crystallography using ARP/wARP version 7. Nat. Protoc. 3:1171–1179. doi:10.1038/nprot.2008.91.

Laudermilch, E., P.L. Tsai, M. Graham, E. Turner, C. Zhao, and C. Schlieker. 2016. Dissecting Torsin/cofactor function at the nuclear envelope: A genetic study. Mol. Biol. Cell. 27:3964– 3971. doi:10.1091/mbc.E16-07-0511.

Legland, D., I. Arganda-Carreras, and P. Andrey. 2016. MorphoLibJ: Integrated library and plugins for mathematical morphology with ImageJ. Bioinformatics. 32:3532–3534. doi:10.1093/bioinformatics/btw413.

Liebschner, D., P. V. Afonine, M.L. Baker, G. Bunkoczi, V.B. Chen, T.I. Croll, B. Hintze, L.W. Hung, S. Jain, A.J. McCoy, N.W. Moriarty, R.D. Oeffner, B.K. Poon, M.G. Prisant, R.J. Read, J.S. Richardson, D.C. Richardson, M.D. Sammito, O. V. Sobolev, D.H. Stockwell, T.C. Terwilliger, A.G. Urzhumtsev, L.L. Videau, C.J. Williams, and P.D. Adams. 2019. Macromolecular structure determination using X-rays, neutrons and electrons: Recent developments in Phenix. Acta Crystallogr. Sect. D Struct. Biol. 75:861–877. doi:10.1107/S2059798319011471.

Lin, D.H., T. Stuwe, S. Schilbach, E.J. Rundlet, T. Perriches, G. Mobbs, Y. Fan, K. Thierbach, F.M. Huber, L.N. Collins, A.M. Davenport, Y.E. Jeon, and A. Hoelz. 2016a. Architecture of the symmetric core of the nuclear pore. Science. 352: aaf1015–aaf1015. doi:10.1126/science.aaf1015.

Lin, D.H., T. Stuwe, S. Schilbach, E.J. Rundlet, T. Perriches, G. Mobbs, Y. Fan, K. Thierbach, F.M. Huber, L.N. Collins, A.M. Davenport, Y.E. Jeon, and A. Hoelz. 2016b. Architecture of the symmetric core of the nuclear pore. Science. 352. doi:10.1126/science.aaf1015.

Lobstein, J., C.A. Emrich, C. Jeans, M. Faulkner, P. Riggs, and M. Berkmen. 2016. SHuffle, a novel Escherichia coli protein expression strain capable of correctly folding disulfide bonded proteins in its cytoplasm. Microb. Cell Fact. 15:1–16. doi:10.1186/s12934-016-0512-9.

Macaulay, C., and D.J. Forbes. 1996. Assembly of the nuclear pore: Biochemically distinct steps revealed with NEM, GTPγS, and BAPTA. J. Cell Biol. 132:5–20. doi:10.1083/jcb.132.1.5.

Marshall, I.C.B., and K.L. Wilson. 1997. Nuclear envelope assembly after mitosis. Trends Cell Biol. 7:69–74. doi:10.1016/S0962-8924(96)10047-7.

Maul, G.G., H.M. Maul, J.E. Scogna, M.W. Lieberman, G.S. Stein, B.Y.L. Hsu, and T.W. Borun. 1972. Time sequence of nuclear pore formation in phytohemagglutinin-stimulated lymphocytes and in hela cells during the cell cycle. J. Cell Biol. 55:433–447. doi:10.1083/jcb.55.2.433.

McCoy, A.J. 2006. Solving structures of protein complexes by molecular replacement with Phaser. Acta Crystallogr. Sect. D Biol. Crystallogr. 63:32–41. doi:10.1107/S0907444906045975.

Miller, B.R., M. Powers, M. Park, W. Fischer, and D.J. Forbes. 2000. Identification of a new vertebrate nucleoporin, Nup188, with the use of a novel organelle trap assay. Mol. Biol. Cell. 11:3381–3396. doi:10.1091/mbc.11.10.3381.

Mosalaganti, S., A. Obarska-Kosinska, M. Siggel, R. Taniguchi, B. Turoňová, C.E. Zimmerli, K. Buczak, F.H. Schmidt, E. Margiotta, M.T. Mackmull, W.J.H. Hagen, G. Hummer, J. Kosinski, and M. Beck. 2022. AI-based structure prediction empowers integrative structural analysis of human nuclear pores. Science. 376. doi:10.1126/science.abm9506.

Murshudov, G.N., P. Skubák, A.A. Lebedev, N.S. Pannu, R.A. Steiner, R.A. Nicholls, M.D. Winn, F. Long, and A.A. Vagin. 2011. REFMAC5 for the refinement of macromolecular crystal structures. Acta Crystallogr. Sect. D Biol. Crystallogr. 67:355–367. doi:10.1107/S0907444911001314.

Muyldermans, S. 2013. Nanobodies: Natural Single-Domain Antibodies. Annu. Rev. Biochem. 82:775–797. doi:10.1146/annurev-biochem-063011-092449.

Newport, J. 1987. Nuclear reconstitution in vitro: Stages of assembly around protein-free DNA. Cell. 48:205–217. doi:10.1016/0092-8674(87)90424-7.

Nordeen, S.A., K.R. Andersen, K.E. Knockenhauer, J.R. Ingram, H.L. Ploegh, and T.U. Schwartz. 2020. A nanobody suite for yeast scaffold nucleoporins provides details of the nuclear pore complex structure. Nat. Commun. 11:1–13. doi:10.1038/s41467-020-19884-6.

Onischenko, E., J.H. Tang, K.R. Andersen, K.E. Knockenhauer, P. Vallotton, C.P. Derrer, A. Kralt, C.F. Mugler, L.Y. Chan, T.U. Schwartz, and K. Weis. 2017. Natively Unfolded FG Repeats Stabilize the Structure of the Nuclear Pore Complex. Cell. 171:904–917.e19. doi:10.1016/j.cell.2017.09.033.

Otsu, N. 1979. A threshold selection method from gray level histograms. IEEE Trans. Syst. Man. Cybern. 9:62–66. doi:10.1109/TSMC.1979.4310076.

Otsuka, S., K.H. Bui, M. Schorb, M. Julius Hossain, A.Z. Politi, B. Koch, M. Eltsov, M. Beck, and J. Ellenberg. 2016. Nuclear pore assembly proceeds by an inside-out extrusion of the nuclear envelope. Elife. 5:1–23. doi:10.7554/eLife.19071.

Otsuka, S., and J. Ellenberg. 2018. Mechanisms of nuclear pore complex assembly – two different ways of building one molecular machine. FEBS Lett. 592:475–488. 10.1002/1873-3468.12905.

Otsuka, S., A.M. Steyer, M. Schorb, J.K. Hériché, M.J. Hossain, S. Sethi, M. Kueblbeck, Y. Schwab, M. Beck, and J. Ellenberg. 2018. Postmitotic nuclear pore assembly proceeds by radial dilation of small membrane openings. Nat. Struct. Mol. Biol. 25:21–28. doi:10.1038/s41594-017-0001-9.

Peroutka, R.J., N. Elshourbagy, T. Piech, and T.R. Butt. 2008. Enhanced protein expression in mammalian cells using engineered SUMO fusions: Secreted phospholipase A 2. Protein Sci. 17:1586–1595. doi:10.1110/ps.035576.108.

Pettersen, E.F., T.D. Goddard, C.C. Huang, G.S. Couch, D.M. Greenblatt, E.C. Meng, and T.E. Ferrin. 2004. UCSF Chimera - A visualization system for exploratory research and analysis. J. Comput. Chem. 25:1605–1612. doi:10.1002/jcc.20084.

Pleiner, T., M. Bates, and D. Görlich. 2018. A toolbox of anti-mouse and anti-rabbit IgG secondary nanobodies. J. Cell Biol. 217:1143–1154. doi:10.1083/jcb.201709115.

Pleiner, T., M. Bates, S. Trakhanov, C.T. Lee, J.E. Schliep, H. Chug, M. Böhning, H. Stark, H. Urlaub, and D. Görlich. 2015. Nanobodies: Site-specific labeling for super-resolution imaging, rapid epitope-mapping and native protein complex isolation. Elife. 4:1–21. doi:10.7554/eLife.11349.

Prophet, S.M., B.S. Naughton, and C. Schlieker. 2022. p97/UBXD1 Generate Ubiquitylated Proteins That Are Sequestered into Nuclear Envelope Herniations in Torsin-Deficient Cells. Int. J. Mol. Sci. 23. doi:10.3390/ijms23094627.

Rampello, A.J., E. Laudermilch, N. Vishnoi, S.M. Prophet, L. Shao, C. Zhao, C. Patrick Lusk, and C. Schlieker. 2020. Torsin ATPase deficiency leads to defects in nuclear pore biogenesis and sequestration of MLF2. J. Cell Biol. 219. doi:10.1083/JCB.201910185.

Rasala, B., C. Ramos, A. Harel, and D.J. Forbes. 2008. Capture of AT-rich chromatin by ELYS recruits POM121 and NDC1 to initiate nuclear pore assembly. Mol. Biol. Cell. 19:3982–3996. doi:10.1091/mbc.E08.

Ribbeck, K., and D. Görlich. 2001. Kinetic analysis of translocation through nuclear pore complexes. EMBO J. 20:1320–1330. doi:10.1093/emboj/20.6.1320.

Sachdev, R., C. Sieverding, M. Flötenmeyer, and W. Antonin. 2012. The C-terminal domain of Nup93 is essential for assembly of the structural backbone of nuclear pore complexes. Mol. Biol. Cell. 23:740–749. doi:10.1091/mbc.E11-09-0761.

Schellhaus, A.K., P. De Magistris, and W. Antonin. 2016. Nuclear Reformation at the End of Mitosis. J. Mol. Biol. 428:1962–1985. doi:10.1016/j.jmb.2015.09.016.

Schindelin, J., I. Arganda-Carreras, E. Frise, V. Kaynig, M. Longair, T. Pietzsch, S. Preibisch, C. Rueden, S. Saalfeld, B. Schmid, J.Y. Tinevez, D.J. White, V. Hartenstein, K. Eliceiri, P. Tomancak, and A. Cardona. 2012. Fiji: An open-source platform for biological-image analysis. Nat. Methods. 9:676–682. doi:10.1038/nmeth.2019.

Schmidt, H.B., and D. Görlich. 2015. Nup98 FG domains from diverse species spontaneously phase-separate into particles with nuclear pore-like permselectivity. Elife. 4:1–30. doi:10.7554/eLife.04251.

Schoonderwoert, V., R. Dijkstra, G. Luckinavicius, O. Kobler, and H. van der Voort. 2013. Huygens STED Deconvolution Increases Signal-to-Noise and Image Resolution towards 22 nm. Micros. Today. 21:38–44. doi:10.1017/s1551929513001089.

Siniossoglou, S., M. Lutzmann, H. Santos-Rosa, K. Leonard, S. Mueller, U. Aebi, and E. Hurt. 2000. Structure and assembly of the Nup84p complex. J. Cell Biol. 149:41–53. doi:10.1083/jcb.149.1.41.

Stavru, F., B.B. Hülsmann, A. Spang, E. Hartmann, V.C. Cordes, and D. Görlich. 2006a. NDC1: A crucial membrane-integral nucleoporin of metazoan nuclear pore complexes. J. Cell Biol. 173:509–519. doi:10.1083/jcb.200601001.

Stavru, F., G. Nautrup-Pedersen, V.C. Cordes, and D. Görlich. 2006b. Nuclear pore complex assembly and maintenance in POM121- and gp210-deficient cells. J. Cell Biol. 173:477–483. doi:10.1083/jcb.200601002.

Stuwe, T., C.J. Bley, K. Thierbach, S. Petrovic, S. Schilbach, D.J. Mayo, T. Perriches, E.J. Rundlet, Y.E. Jeon, L.N. Collins, F.M. Huber, D.H. Lin, M. Paduch, A. Koide, V. Lu, J. Fischer, E. Hurt, S. Koide, A.A. Kossiakoff, and A. Hoelz. 2015. Architecture of the fungal nuclear pore inner ring complex. Science. 350:56–64. doi:10.1126/science.aac9176.

Stuwe, T., L. Schada Von Borzyskowski, A.M. Davenport, and A. Hoelz. 2012a. Molecular basis for the anchoring of proto-oncoprotein Nup98 to the cytoplasmic face of the nuclear pore complex. J. Mol. Biol. 419:330–346. doi:10.1016/j.jmb.2012.03.024.

Stuwe, T., L. Schada Von Borzyskowski, A.M. Davenport, and A. Hoelz. 2012b. Molecular Basis for the Anchoring of Proto-Oncoprotein Nup98 to the Cytoplasmic Face of the Nuclaer Pore Complex. J. Mol. Biol. 419:330–346. doi:10.1016/j.jmb.2012.03.024.Molecular.

Tegunov, D., and P. Cramer. 2019. Real-time cryo-electron microscopy data preprocessing with Warp. Nat. Methods. 16:1146–1152. doi:10.1038/s41592-019-0580-y.

Vera-Rodriguez, A., S. Frey, and D. Görlich. 2019. Engineered SUMO/protease system identifies Pdr6 as a bidirectional nuclear transport receptor. J. Cell Biol. 218:2006–2020. doi:10.1083/jcb.201812091.

VisionVR, A. 2020. arivis Vision4D, 3.1.1.

Vollmer, B., M. Lorenz, D. Moreno-Andrés, M. Bodenhöfer, P. De Magistris, S.A. Astrinidis, A. Schooley, M. Flötenmeyer, S. Leptihn, and W. Antonin. 2015. Nup153 Recruits the Nup107-160 Complex to the Inner Nuclear Membrane for Interphasic Nuclear Pore Complex Assembly. Dev. Cell. 1–12. doi:10.1016/j.devcel.2015.04.027.

Vollmer, B., A. Schooley, R. Sachdev, N. Eisenhardt, A.M. Schneider, C. Sieverding, J. Madlung, U. Gerken, B. MacEk, and W. Antonin. 2012. Dimerization and direct membrane interaction of Nup53 contribute to nuclear pore complex assembly. EMBO J. 31:4072–4084. doi:10.1038/emboj.2012.256.

Walther, T.C., A. Alves, H. Pickersgill, I. Loïodice, M. Hetzer, V. Galy, B.B. Hülsmann, T. Köcher, M. Wilm, T. Allen, I.W. Mattaj, and V. Doye. 2003a. The conserved Nup107-160 complex is critical for nuclear pore complex assembly. Cell. 113:195–206. doi:10.1016/S0092-8674(03)00235-6.

Walther, T.C., P. Askjaer, M. Gentzel, A. Habermann, G. Griffiths, M. Wilm, I.W. Mattaj, and M. Hetzer. 2003b. RanGTP mediates nuclear pore complex assembly. Nature. 424:689–694. doi:10.1038/nature01898.

Walther, T.C., H.S. Pickersgill, V.C. Cordes, M.W. Goldberg, T.D. Allen, I.W. Mattaj, and M. Fornerod. 2002. The cytoplasmic filaments of the nuclear pore complex are dispensable for selective nuclear protein import. J. Cell Biol. 158:63–77. doi:10.1083/jcb.200202088.

Weberruss, M., and W. Antonin. 2016. Perforating the nuclear boundary - how nuclear pore complexes assemble. J. Cell Sci. 129:4439–4447. doi:10.1242/jcs.194753.

Wiese, C., M.W. Goldberg, T.D. Allen, and K.L. Wilson. 1997. Nuclear envelope assembly in Xenopus extracts visualized by scanning EM reveals a transport-dependent “envelope smoothing” event. J. Cell Sci. 110:1489–1502.

Williams, C.J., J.J. Headd, N.W. Moriarty, M.G. Prisant, L.L. Videau, L.N. Deis, V. Verma, D.A. Keedy, B.J. Hintze, V.B. Chen, S. Jain, S.M. Lewis, W.B. Arendall, J. Snoeyink, P.D. Adams, S.C. Lovell, J.S. Richardson, and D.C. Richardson. 2018. MolProbity: More and better reference data for improved all-atom structure validation. Protein Sci. 27:293–315. doi:10.1002/pro.3330.

Wing, C.E., Fung, H.Y.J. & Chook, Y.M. 2022. Karyopherin-mediated nucleocytoplasmic transport. Nat Rev Mol Cell Biol. 23:307–328.

Wu, J., M.J. Matunis, D. Kraemer, G. Blobel, and E. Coutavas. 1995. Nup358, a cytoplasmically exposed nucleoporin with peptide repeats, Ran-GTP binding sites, zinc fingers, a cyclophilin A homologous domain, and a leucine-rich region. J. Biol. Chem. 270:14209–14213. doi:10.1074/jbc.270.23.14209.

Wühr, M., R.M. Freeman, M. Presler, M.E. Horb, L. Peshkin, S.P. Gygi, and M.W. Kirschner. 2014. Deep proteomics of the xenopus laevis egg using an mRNA-derived reference database. Curr. Biol. 24:1467–1475. doi:10.1016/j.cub.2014.05.044.

Wurm, C.A., K. Kolmakov, F. Göttfert, H. Ta, M. Bossi, H. Schill, S. Berning, S. Jakobs, G. Donnert, V.N. Belov, and S.W. Hell. 2012. Novel red fluorophores with superior performance in STED microscopy. Opt. Nanoscopy. 1:1–7. doi:10.1186/2192-2853-1-7.

Yokohama, N., N. Hayashi, T. Seki, N. Panté, T. Ohba, K. Nishii, K. Kuma, T. Hayashida, T. Miyata, U. Aebi, M. Fukui, and T. Nishimoto. 1995. A giant nucleopore protein that binds Ran/TC4. Nature. 376:1181–1185.

Zhu, X., G. Huang, C. Zeng, X. Zhan, K. Liang, Q. Xu, Y. Zhao, P. Wang, Q. Wang, Q. Zhou, Q. Tao, M. Liu, J. Lei, C. Yan, and Y. Shi. 2022. Structure of the cytoplasmic ring of the Xenopus laevis nuclear pore complex. Science. 376. doi:10.1126/science.abl8280.

Zivanov, J., T. Nakane, and S.H.W. Scheres. 2020. Estimation of high-order aberrations and anisotropic magnification from cryo-EM data sets in RELION-3.1. IUCrJ. 7:253–267. doi: 10.1107/S2052252520000081.

